# *Pseudomonas* isolates from ponds populated with duckweed prevent disease caused by pathogenic *Pseudomonas* species

**DOI:** 10.1101/2022.12.09.519836

**Authors:** E.L Baggs, F.G Stark, M.B Tiersma, K.V Krasileva

## Abstract

Duckweeds are notoriously invasive plants. They are successful in inhabiting diverse environments, despite their lack of conventional immune pathways that are essential for disease resistance in other plant species. It is unclear how duckweeds thrive in the absence of these immune pathways. In this study, we investigated the effect of bacteria from duckweeds’ natural habitat on disease progression utilizing the duckweed-*Pseudomonas* pathosystem. Through nanopore sequencing of 16S and ITS rDNA amplicons we identified duckweed-associated bacterial and fungal genera present at three environmental sites. The pond filtrate from one of the three environmental locations primed duckweed’s pathogen defenses leading to a reduction in disease symptoms. Furthermore, we were able to identify bacterial isolates from the filtrate that protect duckweed from disease symptoms upon *Pseudomonas* pathogen inoculation. The isolated protective bacteria belong to the *Pseudomonas* genus, and we demonstrated antagonistic interactions between the pathogen and beneficial strains *in vitro* and *in vivo*. The ability of our environmental isolates to protect against *Pseudomonas* pathogens appears to be plant/species specific as environmental strains showed no protective effect against *Pseudomonas* pathogens in *Arabidopsis* assays. Genome sequencing of the beneficial *Pseudomonas* strains showed the presence of several genes involved in bacterial competition. We have thus demonstrated that *Pseudomonas* species from duckweeds natural habitat can successfully antagonize other plant pathogens.

## Introduction

Plants are holobionts that host a complex assemblage of microbes, which interact with one another and the plant (Hassani et al., 2018). The outcomes of these interactions can be affected by environmental fluctuations, ecological factors, priority effects, host genotype, and microbial genotypes (Henry et al., 2021; Kohl, 2020; Vishwakarma et al., 2020). Within the microbiome, fungi and bacteria are known to interact in a variety of ways, which enables the maintenance of stable communities (Coyte et al., 2015; Hassani et al., 2018; Paasch & He, 2021). To form a stable community, microbes can learn to live in mutualism, however, often they must outcompete other microbes through either ecological competition or interference competition (Coyte et al., 2015; Hassani et al., 2018; Paasch & He, 2021). Ecological competition includes overcoming or imposing nutrient limitations, niche exclusion, and quorum sensing disruption (Bauer et al., 2018; Ghoul & Mitri, 2016; Hibbing et al., 2010). In contrast, interference competition involves direct damage to competitor cells (Bauer et al., 2018; Coyte & Rakoff-Nahoum, 2019). Interference competition can be further subdivided into contact-dependent toxin delivery systems, including Type III, Type IV, and Type VI secretion systems, or contact-independent toxin systems, such as bacteriocin/tailocin and secondary metabolites (Coyte & Rakoff-Nahoum, 2019). Interference competition between plant-associated bacteria is poorly understood. Many studies demonstrate an outcome rather than a mechanism of biocontrols; interference mechanisms such as T6SS were only discovered relatively recently (Hood et al., 2010; Schwarz et al., 2010); and microbiome interactions are complex, requiring interdisciplinary approaches at the intersection of plant pathology, microbiome research, and ecology.

The microbiome of plants traditionally considers the terrestrial plant microbiome which has been characterized in a number of plant species, primarily focusing on the distinct rhizosphere and phyllosphere ecological niches (Kwak et al., 2018; Lundberg et al., 2012; Morella et al., 2020; Peiffer et al., 2013; Trivedi et al., 2020). It remains unclear whether many established plant microbiome principles that are derived from terrestrial plant experiments hold true in the understudied aquatic plant microbiome. Studies have begun to investigate the relative abundance of different bacteria phylum and genera associated with the duckweed microbiome (Acosta et al., 2020; O’Brien et al., 2020, 2022; Yoneda et al., 2021). Previous work has focused on utilizing duckweed for bioremediation (Acosta et al., 2020; Inoue et al., 2022; O’Brien et al., 2020, 2022; Yoneda et al., 2021), whilst our understanding of bacterial pathogens and their interaction with the aquatic plant microbiome is still limited. Additionally, the fungal constituents of the duckweed microbiome are yet to be fully understood, although many studies have investigated duckweeds as potential sources of antifungal compounds (Das et al., 2012; Effiong & Sanni, 2009; Gülçin et al., 2010). Typically, within the plant microbiome, there are plant growth-promoting bacteria and fungi which have been shown to promote the growth of both terrestrial and aquatic plant species (Hossain et al., 2007, 2017; Ramakrishna et al., 2019; Suzuki et al., 2014; Toyama et al., 2022). Microbiome-mediated mechanisms for promoting growth include: facilitating nutrient acquisition, nitrogen fixation, modulation of phytohormones, and competition with other microbes, including pathogens (Ishizawa et al., 2019, 2020; Olanrewaju et al., 2017; Shalev et al., 2022; Suzuki et al., 2014; Toyama et al., 2022; Yamakawa et al., 2018; Yoneda et al., 2021).

Plant pathogens belong to dozens of fungal, bacterial, and oomycota genera. Bacterial pathogens from the *Pseudomonas* and *Xanthomonas* genera are globally distributed and contain species highlighted as within the top five economically and scientifically important bacterial phytopathogens (Mansfield et al., 2012). *Pseudomonas syringae* pv. *tomato* (*Pst*) DC3000 and *Arabidopsis thaliana* constitute a model pathosystem (Katagiri et al., 2002; Xin & He, 2013). Once inside the plant, *Pst* DC3000 multiplies to high population density in the substomatal cavity and mesophyll (Katagiri et al., 2002). In order to thrive inside the leaf, the pathogen creates its own niche through the use of effectors to induce water soaking, creating a humid, photosynthate-rich inter-cellular environment (Hernandez & Lindow, 2019). *Pseudomonas syringae* pv. *syringae* B728a (*Pss* B728a) is another pathogenic *Pseudomonas* isolated from *Phaseolus vulgaris* (Hirano et al., 1995) with well-studied virulence mechanisms (Feil et al., 2005; Hernandez & Lindow, 2019; Hirano et al., 1999; Lacombe et al., 2010). *Pss* B728a typically has a longer epiphytic phase on the host plant surface, thought to be due to its greater stress tolerance (Feil et al., 2005; Hirano & Upper, 2000). Disease symptoms following *Pss* B728a infection can be due to colonization of either the leaf surface or the apoplast (Hirano & Upper, 2000). Duckweed species have been shown to be susceptible to *Pss* B728a, *Pst* DC3000, and several pathogenic *Xanthomonas* strains (Baggs et al., 2022). Economically and scientifically important fungal pathogens use an arsenal of strategies to infiltrate the plant and cause disease. Fungal pathogens are also among the dominant causal agents of plant disease with the potential to decimate entire plant populations (Doehlemann et al., 2017). Strategies employed by fungal pathogens include: plant polymer and cell wall degrading enzymes, specialized infection structures such as the appressorium/ haustorium, and virulence factors such as effectors known to evade or manipulate the plant’s immune response (Dean et al., 2012).

Duckweeds have recently been developed as an alternative model system to *A. thaliana* when studying bacterial pathogens (Baggs et al., 2022). The small size of duckweeds, short generation time, small proteome, and low maintenance inputs make them an ideal system for rapid hypothesis testing (Acosta et al., 2021). In contrast to the majority of angiosperms, including *A. thaliana*, duckweeds are missing a conserved immune pathway consisting of EDS1, PAD4, and RNLs (Baggs et al. 2020; Baggs et al. 2022; Lapin et al. 2019). Contrary to the reduced copy number of the NLR intracellular immune receptor gene family, the antimicrobial protein family characterized by the presence of the MiAMP1 domain was expanded and differentially expressed upon pathogen challenge in duckweeds (Acosta et al., 2021; An et al., 2019; Baggs et al., 2022). Despite the overlap between duckweed species and several plant pathogen ranges (Cai et al., 2011; Gutiérrez-Barranquero et al., 2019; Potnis et al., 2015), natural duckweed populations show sparse signs of visible disease (Xu et al., 2015). Duckweed pathogenic fungi include *Tracya lemnae* (Vanky, 1981) and *Olipidium amoebae* (Fisch, 1884; Gaumann, 1928), but fungal community ecology has yet to be described in duckweed. The ubiquitous nature of *Pseudomonas* pathogens and model duckweed pathosystem *Spirodela polyrhiza - Pst* DC3000 (Baggs et al., 2022) prompted us to investigate whether natural populations of duckweed are in contact with bacterial pathogens. Furthermore, we were interested in how the natural environment could affect the outcome of duckweed populations exposed to pathogenic bacteria.

To date, it is unclear how plant species from aquatic environments, or plants without EDS1 defense pathways, would be able to survive environmental pathogen pressure. We explored the role of the water filtrates and the plant microbiome in protection of duckweed species against *Pseudomonas* pathogens. Our results confirm that microbiome-mediated protection against disease is present in an aquatic plant environment. Furthermore, we discovered two bacterial strains that confer disease protection for duckweed species challenged with *Pseudomonas* pathogens. We show that the environmental bacterial isolates reduce *Pst* DC3000 colony forming units in the absence of the plant host, with the genomes of the protective strains containing an arsenal of bacterial interference mechanisms. However, the inhibition seen *in vitro* is insufficient to prevent *Pseudomonas* from causing disease in *A. thaliana*. Together, our findings suggest a key role of the microbiome in the protection of duckweed from disease.

## Results

### Community structures of duckweed-associated bacteria and fungi differ from those associated with terrestrial plants

To understand how natural duckweed (*Lemnaceae*) populations protect themselves from pathogens, we assessed the impact of their aquatic environment on disease progression. On February 27th 2020, we identified and sampled both water and fronds from three populations of *Lemnaceae* at the University of California Berkeley Botanical Garden (UCB-BG) (Fig 1a). Sites will be referred to herein by their UCB-BG bed IDs: 404, 405, and 923. The sampling was then repeated on November 3rd 2020 for further studies including microbiome analysis. Each site had healthy duckweed populations with little visible to no visible disease symptoms (Fig 1b). Duckweed and water samples were taken at three distinct locations within each site for further analysis (Fig 1c, d). A subset of environmental *Lemnaceae* fronds (fused leaf and stem) were sterilized to allow aseptic propagation, however, only fronds from site 405 were able to survive this stringent sterilization process. Genotyping of the duckweed from the sample site 405, herein referred to as BG405, using the AtpF marker (Wang et al., 2010) identified it as most closely related to *Landoltia punctata* strain DW2701-4 (100% nucleotide identity for AptF (File S1)).

**Figure 1.**
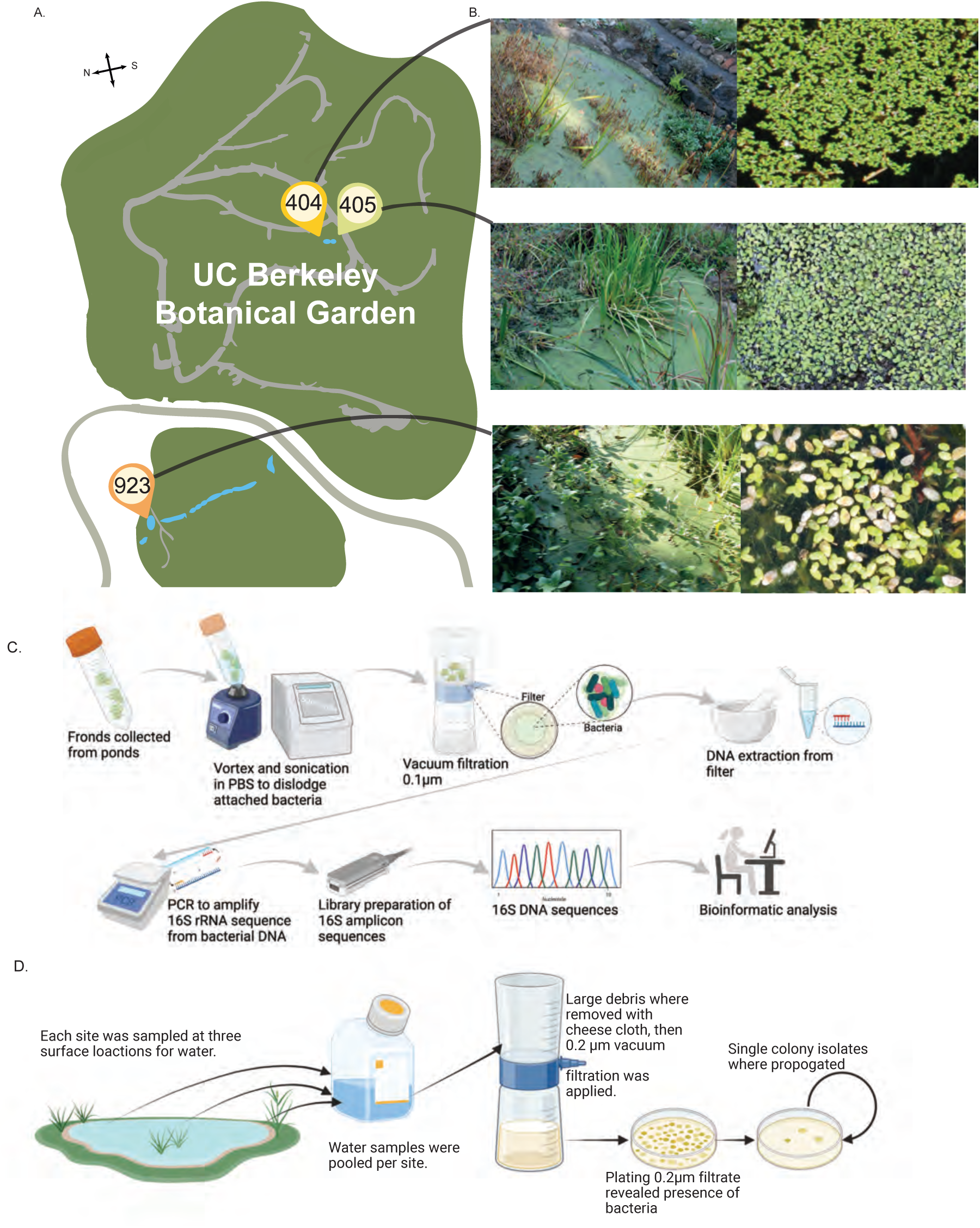
Schematic of location and sample processing for duckweed and its microbiome. A. Schematic map of the UC Berkeley Botanical Garden sampling location. Blue indicates water bodies, gray shows pathways and point locators show specific ponds sampled. B. Images of the three ponds at time of sampling. C. Experimental procedure for the 16S rRNA amplicon sequencing. D. Experimental procedure for water sampling from each pond.

To identify in a culture-independent manner which bacteria and fungi live in close association with duckweed, we washed microbial communities from environmental fronds into Phosphate-Buffered Saline (PBS). The solution was passed through a 0.1 μm filter, and DNA was then extracted from the filter (Fig 1c). Five biological replicates of duckweed-associated microbial DNA were extracted from each of the three sites. Followed by library preparation and nanopore minION sequencing. Taxonomic assignments were made for the bacterial full-length (V1-V9) 16S rDNA amplicons against the SILVA databases. Bacterial richness at the family and genus level, before and after normalization, was highest at site 404 and lowest at site 923 (Table 2). Genus richness was not saturated when considering biological replicates within sites separately but was when replicates were pooled by site (Fig S1). Shannon diversity index and Simpson diversity index applied to family-level taxon assignments showed that, despite slightly lower taxa richness, site 405 had the highest level of diversity of bacterial families among the three sites.

**Table 1:** A subset of genomic features identified by antiSMASH 5.0 and SecReT6 in the genome of 5BF in comparison to Pst DC3000 (NC_004578.1) and Pss B728a (NC_007005.1). For the full table see Table S15.

| Product of most similar known cluster | Genomic feature | Genomic Location 5BF | Genomic Location 5BT | Present in *Pst* DC3000 or *Pss* B728a | Role in interference competition | Reference |
| --- | --- | --- | --- | --- | --- | --- |
| Pyoverdin, fluorescent siderophore | NRPS | 859,747-910,922 | 2,580,964 - 2,602,394 | *Pss* B728a, *Pst* DC3000 | Pyoverdine is a siderophore which provides iron that enables faster bacterial growth. It can also translocate into mammalian hosts, where it binds and extracts iron and triggers autophagy. | (Kang et al., 2018; Meyer et al., 1996) |
| Type VI secretion il | Contact dependent bactericidal effector secretion system | 2,050,334-2,069,478 | 890,104-918,813 | *Pst* DC3000 | Although related Type VI secretion il were found in *Pst*, they both showed little homology or synteny with the Type VI secretion il system in 5BF or 5BT and one was not intact (Fig S30-32). | (Barret et al., 2011; Chien et al., 2020) |
| Viscosin, antimicrobial lipopeptide | NRPS | 3,015,416-3,115,302 | None | None | Viscosin is a lipopeptide shown to act as a biosurfactant and inhibit *Streptomyces scabies* (Fig S29) | ((Bonnichsen et al., 2015) (Pacheco-Moreno et al., 2021)) |

**Table 2:** Family and Genus level diversity statistics for the three sites sampled after SRS normalization.

| Bacteria | Diversity metric | site 404 | site 405 | site 923 |
| --- | --- | --- | --- | --- |
| Family | Richness | 85-91 | 86-97 | 53-73 |
|  | Shannon | 2.88-2.99 | 3.23-3.40 | 2.48-2.84 |
|  | Simpson | 0.86-0.88 | 0.91-0.94 | 0.82-0.90 |
| **Fungi** |  |  |  |  |
| Family | Richness | 268 | 229 | 262 |
|  | Shannon | 1.53 | 1.34 | 2.55 |
|  | Simpson | 0.46 | 0.39 | 0.77 |

All reads which passed quality filters were able to be assigned to bacterial orders (Table S1). For each successive taxonomic rank below family an increasing percentage of reads were unclassified, with between 8-21% of reads unclassified at the genus level. The relative abundance of bacterial order classifications across the three sample sites revealed a stable pattern of dominant taxa across sites. Burkholderiales are the most abundant order in all but one site where it was the second most abundant (Fig. 2). Family-level classification showed that Comamonadaceae was the only family with more than 10% classified reads across all sites (Fig. S2, Table S2). The only other family that accounted for more than 3% of classified reads in all replicates was Sphingomonadaceae. The overlap between the core terrestrial bacterial microbiome and the duckweed bacterial microbiome is consistent with prior studies (Acosta et al., 2020; Inoue et al., 2022; O’Brien et al., 2022). Most other families were assigned less than 5% of classified reads at any given site, although the stability of relative abundance varied across taxa. The relative abundance was quite stable (±5%) across all samples for families such as Sphingomonadaceae and Chitinophagacea. Conversely, other families such as Pseudomonadaceae showed high variance in abundance from 0.4% in biological replicate 3 at site 405 to 15% at biological replicate 3 at site 923. The trend of high variability in relative abundance of Pseudomonadaceae was reproduced with *Pseudomonas* at the genus level (Fig 2a, Table S3) accounting for 0.3-2.3% of total abundance among genera in site 405 but 2.6-16% at site 923. Family-level classification of the Lemnaceae-associated microbiome shows there is variability in the dominance and stability of bacterial families across sampled sites.

**Figure 2.**
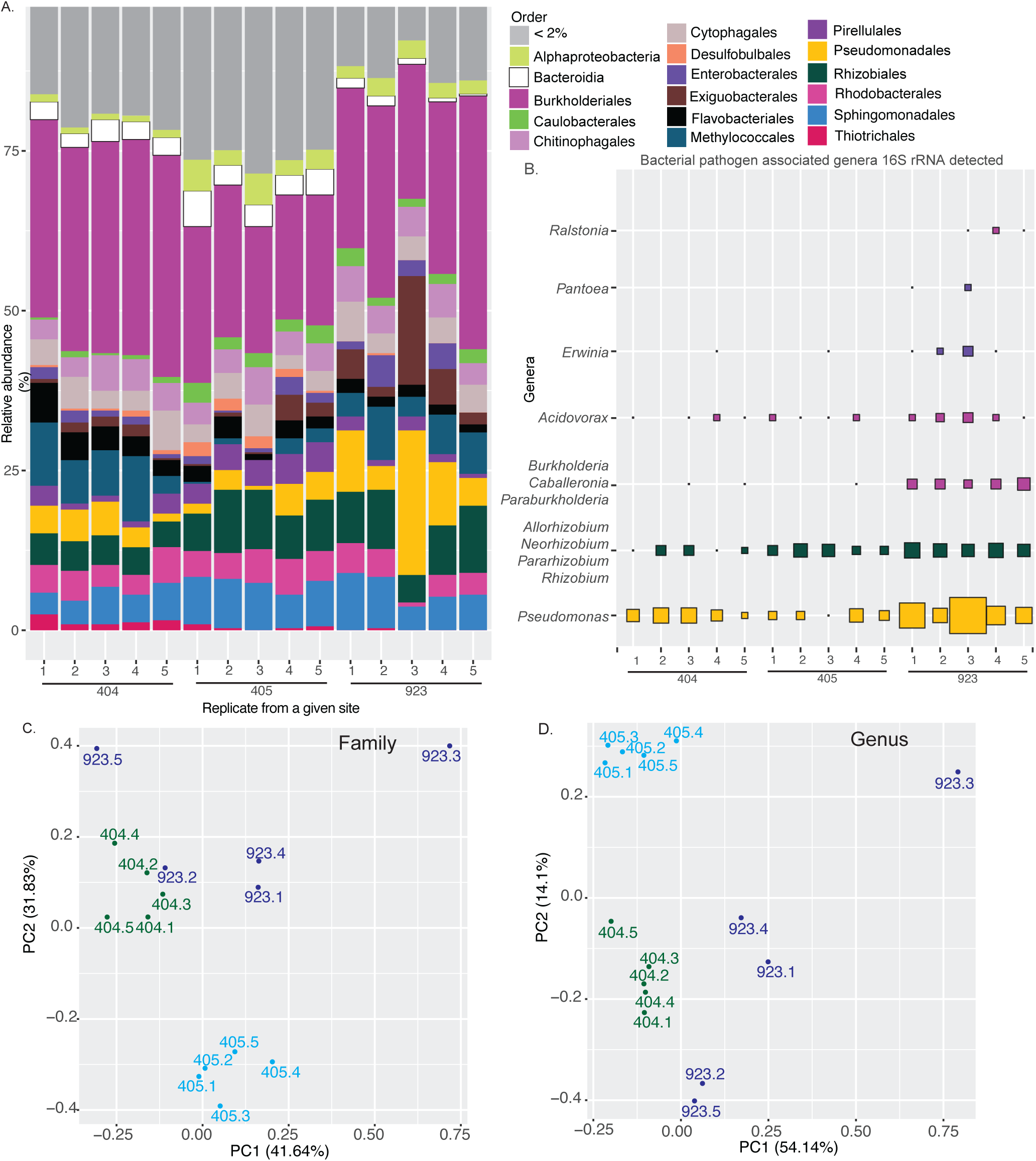
Microbiome composition at the UC Botanical Garden sites. A. Percentage of the 16S rDNA amplicons corresponding to SILVA 16S rRNA family classification for taxa which account for >1% of total classified reads for at least one site. The size of of the bar is proportional to percent of total classified 16S rDNA amplicons. B The percentage of ITS amplicons corresponding to plant pathogen associated ITS present at atleast one site. The size of the squares is proportional to percent of total classified ITS rDNA amplicons for that genera. C. Biplot of sites given bacterial family count data. D. As in B but upon genus level classification.

Despite the close physical proximity of sites 404 and 405, biplots showed clear separation of all sites given their bacterial family (PC1 41.6%, PC2 31.8%) or genera compositions (PC1 54.14%, PC2 14.1%) (Fig 2b, c). Despite further geographic distance, sites 404 and 923 clustered together when considering family-level classification driven by them having higher levels of Comamonadaceae, Methylomondaceae, Pseudomonadaceae, and Exiguobacteraceae than site 405 (Fig. S3a). To investigate this clustering further, we looked into the genera-level separation between sites. At this higher resolution, sites 923 and 404 clustered separately with the relative abundance of Pseudomonas, *Rhodoferax*, and unknown bacteria being the main drivers separating clusters (Fig. S3b). Comparison of the duckweed bacterial microbiomes across sites showed evidence of stable taxa that remain at a similar level across all sites along with a collection of bacteria whose abundance is highly variable, including *Pseudomonas* species.

Fungal taxa richness was highest at site 404 and lowest at site 405 (Table 2). Whilst site 923 had the highest Shannon diversity and Simpson diversity when considering genus and family-level taxonomic assignment. Phyla level classification revealed a large percentage of the ITS amplicons at each site (43-63%) didn’t belong to fungal taxa but instead were assigned unidentified or to non-fungal phyla (Table S4). Within the reads assigned to a phyla, the relative abundance of chytridiomycota as the first or second most abundant phyla ((33-40% of reads classified at phyla level) Table S5) suggests the fungal community of aquatic plants differs from terrestrial plants where ascomycota and basidiomycota are typically the dominant community members (Bergelson et al., 2019; Fuentes et al., 2020; Qian et al., 2019). However, Ascomycota was the second most abundant phyla (19-45% of reads) while Basidiomycota was the third most abundant phyla (9-22% of reads). Unlike the bacterial communities across the three sites, there was not one clearly dominant fungal order (Fig. 3a, Table S6). At site 923 Capnoideales taxa were the most prevalent whilst at sites 405 and 404 the highest abundance orders were Polyproales and Hypocreales. Fungi from the Capnoideales and Hypocreales are among the top 10 in relative abundance across all three sites, but this is not the case for Polypropales. The Hypocreales and Capnodiales were core members of the microbiome of *Arabis alpina* root (Almario et al., 2017) and *Arabidopsis thaliana* leaves (Almario et al., 2022) respectively. Tremellales, Helotales, and Pleosporales are also orders which show conserved high relative abundance orders across terrestrial plants and duckweed.

**Figure 3.**
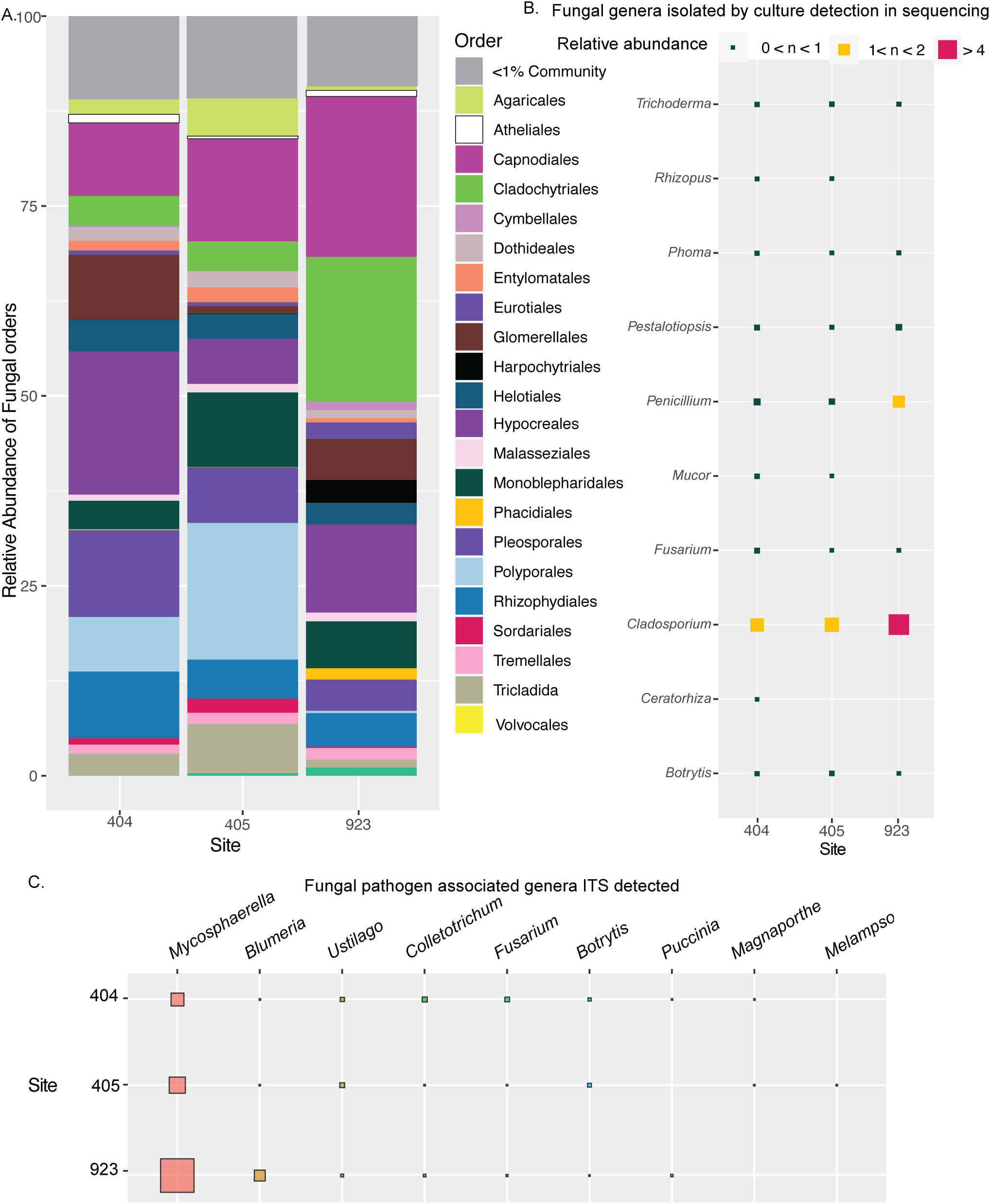
Percentage abundance of fungi from ITS amplicon sequencing at the three sites at UC Berkeley Botanical Garden. A. Percentage of the ITS amplicons corresponding to different fungal orders.The size of the colored bar is proportional to percent of total classified ITS amplicons for that order. B. The percentage of ITS amplicons corresponding genera which where morphotyped from samples at the UC Berkeley Botanical Garden. C.Percentage of the ITS amplicons corresponding to plant pathogen associated fungal genera present at atleast one site. The size of the squares is proportional to percent of total classified ITS rDNA amplicons for that genera.

In order to cross-reference our ITS amplicon sequencing data, we sought to identify fungal isolates that we were able to recover from the material collected at the UC-BG (Fig 3b). Fungal isolates were cultured either from the 0.2 μM PES filter that pond water was passed through or from surface sterilized duckweed populations as described in the methodology. Fungi were grouped into 19 distinct morphotypes through colony phenotype and microscopy then assigned to genera based on 5.8S Fun - ITS4 sequencing (Table S7). From our sequencing data we were able to recover reads assigned to all the morpho-type based taxa (Table S8-10) with the exception of *Galactomycetes* genera, however, reads were assigned to the family (Dipodascaceae) which contains *Galactomycetes* (Table S8).

### The Duckweed microbiome suggests common pathogenic Bacterial and Fungal genera co-occur

The ubiquitous nature of *Pseudomonas* pathogens and the susceptibility seen in the model Lemnaceae pathosystem *Spirodela polyrhiza - Pst* DC3000 (Baggs et al., 2022) prompted us to investigate the relative abundance of genera that contain the most common plant bacterial and fungal pathogen species (Fig 3c, Table S11). We considered the genera taxonomic level, SRS plots suggested saturation of diversity at the genera taxonomic level when considering all biological replicates from one pond, but not if each biological replicate was considered individually (Fig S4). Therefore, an absence in a particular sample is likely to be due to insufficient sampling depth unless the absence is consistent across all samples. We mined our samples for evidence of the presence of 13 bacterial pathogen genera (though without experimental validation we can’t be sure if all members of the genus are pathogens). Site 923 had the highest relative abundance of bacteria pathogen-associated genera within its community composition. This was most evident for the taxa Pantoea, Ralstonia, and Burkholderia-Caballeronia-Parburkholderia. Sites 404 and 405 had more similar relative abundance of pathogen-associated genera although site 405 typically had a lower relative abundance of Pseudomonas but higher Allorhizobium-Neorhizobium-Pararhiuzobium-Rhizobium. The relative abundance of *Pseudomonas* genera was the highest at site 923 at between 2.6-7.8%, and the lowest at site 405 with 0.32-2.3%. Previous work has shown *L. punctata* was susceptible to *Pseudomonas spp*. which are pathogens or endophytes on other plant species (Baggs et al., 2022). Reads assigned to these genera could come from species with a variety of lifestyles ranging from pathogenic to beneficial. We were unable to identify any reads classified to 4 of the 13 bacterial genera (*Streptomycetes*, *Clavibacter, Pectobacterium, Spiroplasma*, and *Xylella*). We cannot preclude the possibility that this is a technical artifact due to the bias of molecular biology methods used rather than the absence of any species belonging to these genera.

All of the nine pathogen-associated fungal genera were recovered from at least one site although often at very low relative abundance (Fig. 6b, Table S12). Notably, two of the morphotyped taxa assignments overlap with fungal pathogen-associated taxa. Site 923 had the highest relative abundance of pathogen-associated genera within its community composition at 3.6% compared to 0.577% and 0.78% for sites 404 and 405, respectively. The higher percentage of pathogen-associated genera at site 923 was largely driven by 3.27% of its total community being *Mycosphaerella*. Although *Mycosphaerella* species are responsible for *Mycosphaerella* blight on a number of plant species, there are also some species within the genera that are endophytes of brown algae (Fries, 1979). Further metagenomics and laboratory studies would be needed to estimate the exact disease burden from bacterial and fungal pathogens.

### Environmental bacteria from pond filtrate protect against disease-causing Pseudomonas on duckweeds

The collected pond water samples were used to investigate the effect of the natural environment on the health of duckweeds. To assay the health implications of the environment on duckweed, we used the *Pst* DC3000 and *S. polyrhiza* pathosystem (Fig 4a) (Baggs, Tiersma, Abramsom, et al., 2022). *Pst* DC3000 has previously been shown to be virulent on *S. polyrhiza* and cause induction of black lesions (Baggs et al., 2022). To reduce the complexity of biological compounds in the pond water, we used filtration. Sterile fronds of the *S. polyrhiza* were primed for 24 hrs in the 0.2 μm filtrate from each site, fronds were then transferred to fresh media and inoculated with *Pst* DC3000. The *S. polyrhiza* plants primed with buffer alone showed symptoms of stunted growth and black lesions upon *Pst* DC3000 treatment comparable with previous descriptions of infection (Baggs et al., 2022). Duckweed primed with site 404 filtrate, when treated with *Pst* DC3000, had symptomatic black lesions comparable to buffer (Fig S5). In contrast, *polyrhiza* primed with site 404 filtrate and then treated with *Pst* DC3000 *hrcC*, a mutant unable to deploy effectors, showed signs of increased frond growth compared to buffer priming. Strikingly, site 405 filtrate priming led to a dramatic decrease in disease symptoms on *S. polyrhiza* (Fig 4b). The decrease in symptoms upon priming in site 405 filtrate could be reproduced when using pond water samples from February 2020 which was collected as part of a pilot study for the November experiment (Fig S5). However, once the 0.2 μm filtrate was boiled (Fig S6) or upon subsequent filtration through a 0.1 μm filter (Fig 4b) the priming effect was lost. Previous work has shown that several bacterial taxa can pass through a 0.2 μm filter but not 0.1 μm (Hahn, 2004). *S. polyrhiza* primed with site 923 filtrate and then inoculated with either *Pst* DC3000 or *Pst* DC3000 *hrcC* showed visible disease symptoms comparable with buffer priming following pathogen treatment (Fig S5).

**Figure 4.**
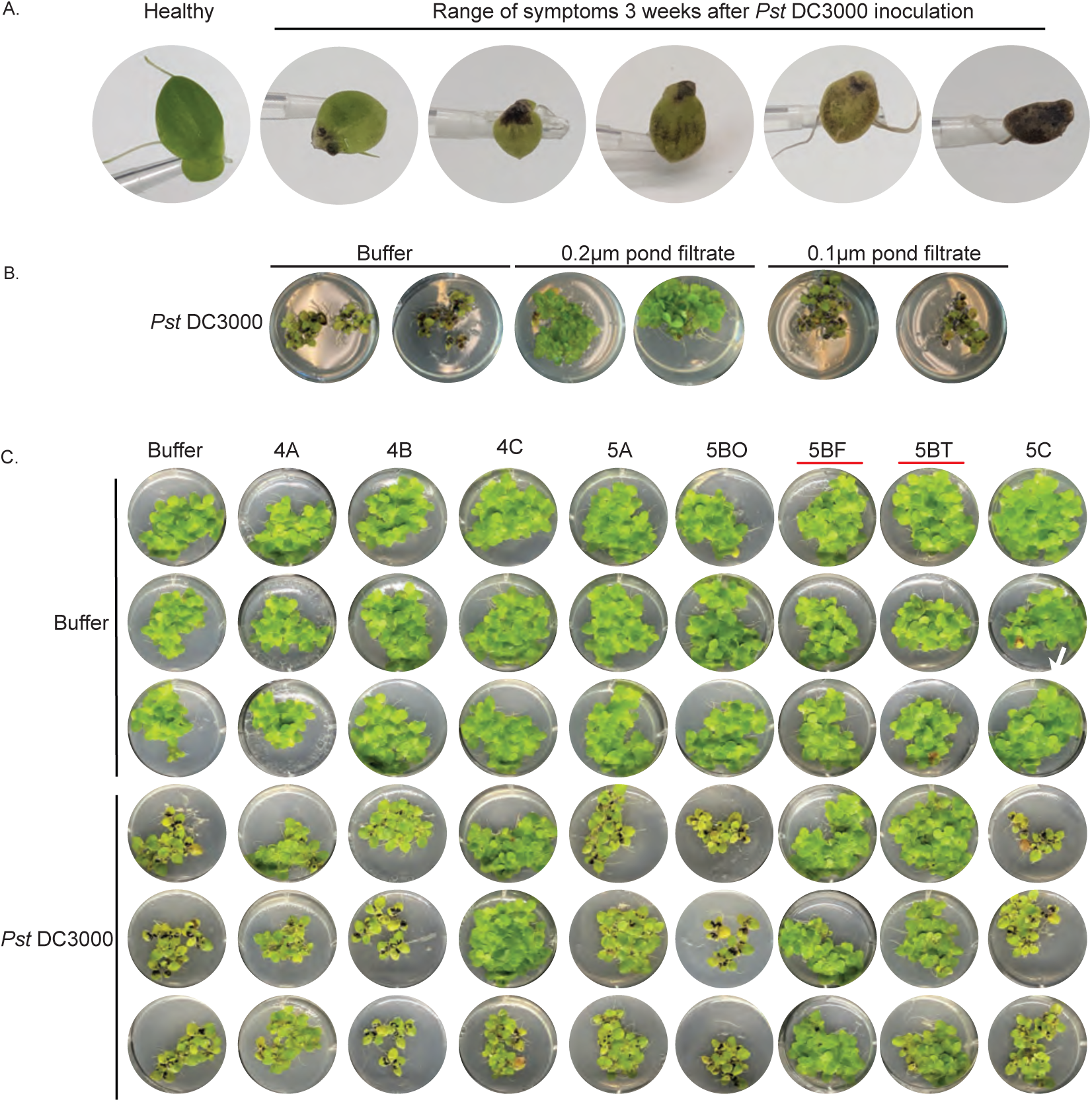
Pond water filtrate and bacterial isolates protection of S. polyrhiza against the pathogen Pst DC3000. A. Depiction of healthy duckweed and range of symptoms upon Pst DC3000 infection. B. Pond water 0.2 μm and 0.1 μm filtrate from site 405, two replicates are shown per treatment (full experiment has been independently replicated 2 times, see Sfig 1). C. Bacterial strains isolated from pond site 404 or 405, as indicated by the first number of the isolate identifier, three replicates within experiment are shown per treatment (full experiment has been independently replicated three times, see Sfig 3 and Sfig 4).

To test if environmental bacteria were contributing to the protective effect of the 0.2 μm filtrates we plated the undiluted filtrates and monitored for microbial growth. Without dilution, we were able to isolate 8 strains that were purified via single colony isolation. After establishing pure colonies, we inoculated *S. polyrhiza* with these isolates individually and together with *Pst* DC3000 (Fig 4c, Fig S7-10). Purified environmental isolates had diverse effects when introduced to *S. polyrhiza*, ranging from no visible effect (5A, 5BO, 5C), to reduced *S. polyrhiza* growth (4A), to a beneficial effect when co-inoculated with pathogenic *Pst* DC3000 (5BF, 5BT). Isolate 5BF showed consistent protection against *Pst* DC3000 infection across replicates, whilst isolate 5BT’s protective effect was variable between replicates. The 16S rDNA genotyping of isolates (Table S13) 5BF and 5BT revealed that they were distinct *Pseudomonas* species with the highest similarity to *Pseudomonas poae* (98.02%) and *Pseudomonas kielensis* (99.21%), respectively.

We also tested whether the protective effect of 5BF and 5BT was specific to a single plant species or pathogen isolate. Another pathogenic *Pseudomonas*, *Pss* B728a, when inoculated on its own, has been previously shown to cause white chlorosis of fronds of *L. punctata* 5635 (Baggs et al., 2022). When we inoculated *Pss* B728a together with isolates 5BT or 5BF, we observed reduced or absent chlorosis on *L. punctata* 5635 (Fig S11-13). Together the results suggest that bacterial isolates 5BF and 5BT can provide disease protection for *S. polyrhiza* and *L. punctata* 5635 against at least two *Pseudomonas* pathogens that have distinct lifestyles and virulence profiles.

We were interested in whether the duckweed strain *L. punctata* BG405, that we isolated and sterilized from site 405, would be naturally resistant to *Pseudomonas* infection. Previous work showed severe infection symptoms could be observed after infection of *L. punctata* 5635 with *Pss* B728a, whilst fewer symptoms were seen upon *Pst* DC3000 infection (Baggs et al., 2022). We therefore investigated *L. punctata* BG405 symptoms upon infection with Pss B728a alone or with 5BF or 5BT. Stunted growth and disease symptoms were evident for the *L. punctata* BG405 after inoculation with *Pss* B728a (Fig S14-17). Furthermore, the occurrence of disease symptoms caused by *Pss* B728a were suppressed in the presence of 5BF and 5BT. This suggests that natural populations of *L. punctata* BG405 are susceptible to *Pss* B728a; however, naturally occurring bacteria from the same ponds can suppress the pathogenicity of *Pss* B728a.

### Genomes of the protective isolates 5BF and 5BT reveal the presence of several natural products and Type VI secretion machinery

To compare the protective *Pseudomonas* isolates to other *Pseudomonas* pathogens virulent on *S. polyrhiza* or *L. punctata* 5635 (Baggs et al., 2022), we sequenced and assembled the genomes of 5BF and 5BT (Fig S18). BUSCO analysis revealed 93.5% and 100% of BUSCOs were present as complete single-copies in the 5BF and 5BT genomes (Table S14). Genomic sequences of 5BF and 5BT allowed us to assign them more confidently to *Pseudomonas* species. The genome sequence of 5BT has 99.5% average nucleotide identity (ANI) to the type specimen genome of *Pseudomonas kielensis* [99109] in NCBI GeneBank, with 90.2% total genome coverage. Further analysis indicated that the *Pseudomonas* sp. with the most similar genome to 5BF was *P. poae* with OrthoANI of 88.9%, this is however insufficient to consider them as the same species (95%) (Lee et al., 2016). Therefore, isolate 5BF and 5BT will herein be referred to as *Pseudomonas* nov. 5BF and *Pseudomonas kielensis* 5BT.

We next annotated gene clusters known for their role in interference competition. In the 6.7 Mbp genome of isolate 5BF, we identified through antiSMASH (Blin et al., 2019) 13 genomic islands that have homology to known secondary metabolite clusters (Table 1, Table S15). Genome annotation further supported the presence of a significant repertoire of interference competition islands with 220 genes classified within the stress response, defense, and virulence subsystem (Fig S19). Type VI, iII, iIII, or iv secretion systems were found in *P.* nov. 5BF or *P. kielensis* 5BT but are not present in *Pst* DC3000 or *Pss* B728a (Table S15). In summary, the genomes of isolate *P.* nov. 5BF and *P. kielensis* 5BT revealed the presence of several genomic features that are candidates for mediating interference competition.

### *P.* nov. 5BF and *P. kielensis* 5BT isolates localize to the same niche as *Pst* DC3000 and inhibit growth of *Pst* DC3000 in *S. polyrhiza* upon co-inoculation

To investigate how *P.* nov. 5BF and *P. kielensis* 5BT protect *S. polyrhiza* species from pathogenic *Pseudomonas,* we first examined *in planta* niche occupancy of the two isolates after flood inoculation. *S. polyrhiza* 5 days post-inoculation (dpi) had populations of *P.* nov. 5BF (Fig 5a-c) and *P. kielensis* 5BT (Fig S24) on the surface, and small colonies of *P.* nov. 5BF were identified within the stomatal cavity. The presence of *P.* nov. 5BF in the substomatal cavity would suggest that it may use stomata to enter the plant; the same niche occupied by *Pst* DC3000 (Baggs et al., 2022).

**Figure 5.**
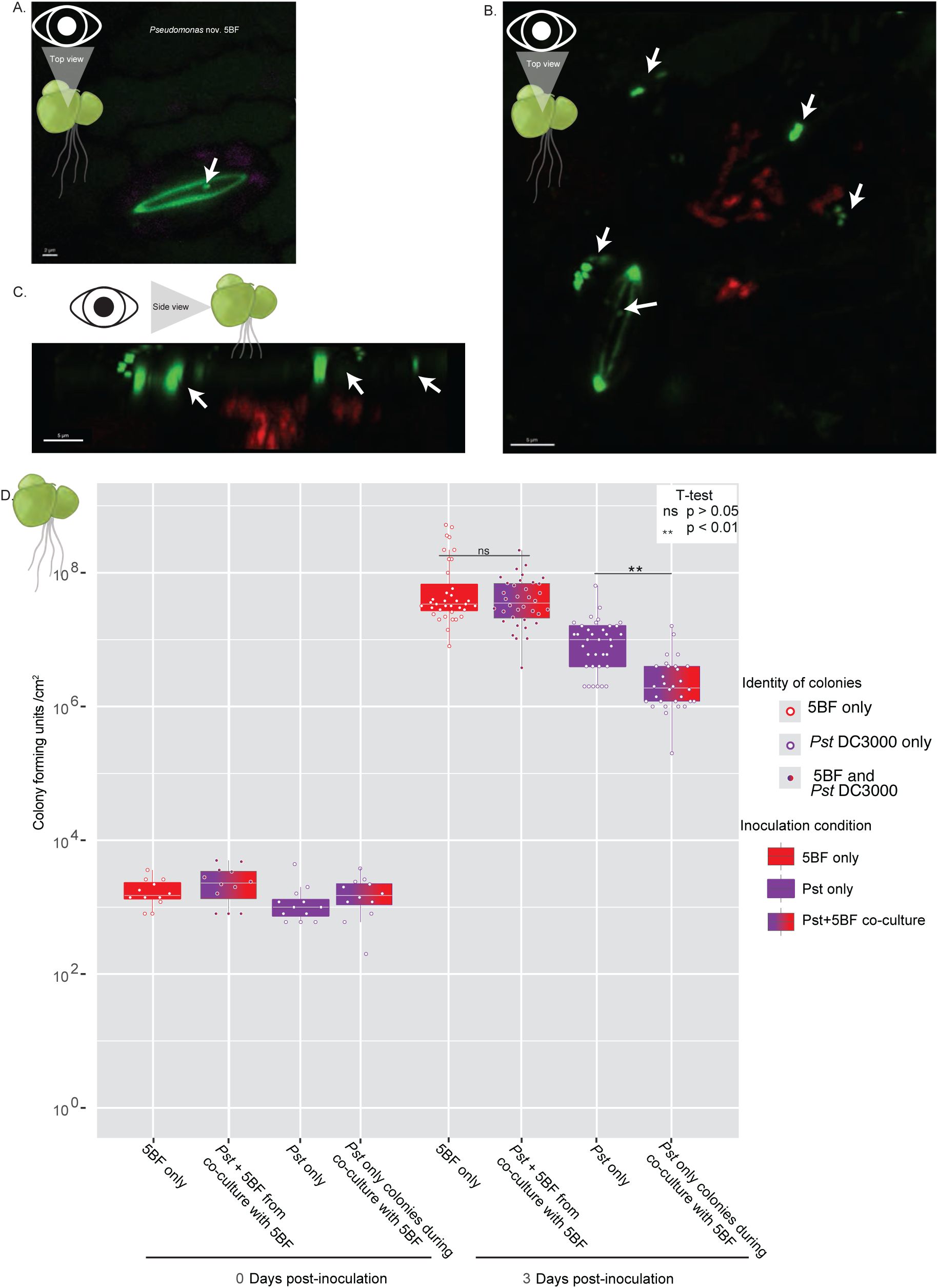
Confocal microscopy of 5BF on *Spirodela polyrhiza* fronds and growth points for *Pseudomonas spp*. A. *S. polyrhiza* frond surface, pink false color shows chlorophyll fluorescence and green indicates SytoBC stained 5BF bacteria. Autofluorescent stomata cell wall can also be seen in green B. *S. polyrhiza* frond surface, *Pst* Dc3000 dsRed is depicted in red and *P. nov.* 5BFis shown in green, autofluorescent stomata cell wall can also be seen in green. C. Side view constructed from Z-stack of *Pst* DC3000 and *P. nov* 5BF colored as in B. D. Quantification of bacterial colonies of *Pseudomonas* sp. in *S. polyrhiza.* P-values shown are from t-test are after Holm-Bonferroni adjustment ((n = 4 each day 0 condition, n = 12 for each day 3 condition except *Pst DC3000* only from co-culture n=10, for each biological replicate in n, three technical replicates were performed) *Table S19)*

To understand why co-inoculation of *P.* nov. 5BF with *Pst* DC3000 led to an absence of symptoms on *S. polyrhiza* (Fig 3), we quantified the colony-forming units (CFU) of these *Pseudomonas* isolates when inoculated alone or together. After inoculation of *P.* nov. 5BF alone at a standard low bacterial load, it was able to grow exponentially on *S. polyrhiza* to a high density of ∼10^8^ CFU/cm^2^ (Fig 5d, Fig S25, Table S10). Co-inoculation of *Pst* DC3000 and *P.* nov. 5BF, when measured at 3 dpi, resulted in a significant decrease (t-test; Holm–Bonferroni adjusted p ! 0.01) in the number of recovered *Pst* DC3000 colonies. The presence of *P.* nov. 5BF appears to inhibit the normal growth of *Pst* DC3000. The visible symptoms of the fronds not sampled for the growth curve were imaged at 10 dpi and supported the conclusion that even at low standard inoculum *P.* nov. 5BF co-inoculation reduces visible symptoms induced upon *Pst* DC3000 treatment (Fig S26).

We then investigated if the 16S rDNA of the protective strains 5BF and 5BT, isolated from pond 405, were present in our amplicon sequence data. To do this we aimed to separate conserved SNPs likely present in the gDNA from those that were technical artifacts generated during PCR and amplicon sequencing. Technical artifact SNPs would likely be randomly distributed in amplicon reads and would be absent from genomic DNA sequence due to high coverage allowing them to be removed during assembly. Alignments were made for sequences with high similarity to isolate 5BF 16S rDNA and isolate 5BT (Fig 6a, b, Fig. S27, Table S16). There were several SNPs present in 5BF that were absent from the closest reference, the 16S rDNA sequence *Pseudomonas poae*. For 7 of 8 positions at which there were reference-specific SNPs, the consensus among aligned reads was nucleotides discrete to *P. nov* 5BF (Table S17,18). The amplicon sequencing therefore supports the presence of the novel species *P.* nov 5BF rather than *P. poae* at the sample sites. There were no discrete SNPs at the 16S rDNA to distinguish 5BT from the bacterial reference strain of *Pseudomonas kielensis.* We were able to recover reads from sites 404 and 923 that passed the threshold of similarity to isolate *P. kielensis*. The false negative of absence of *P. kielensis* from site 404 where it was isolated is not surprising given the sequencing depth is not sufficient for saturation of species-level diversity and that the filtrate from which it was recovered was the product of concentration. The recovery of the reads with high identity to 5BF 16S rDNA, suggests the species is not specific to site 405, which showed protective effects *in vitro,* but rather is likely present at all three sampled sites within the UC Berkeley Botanical Garden.

**Figure 6.**
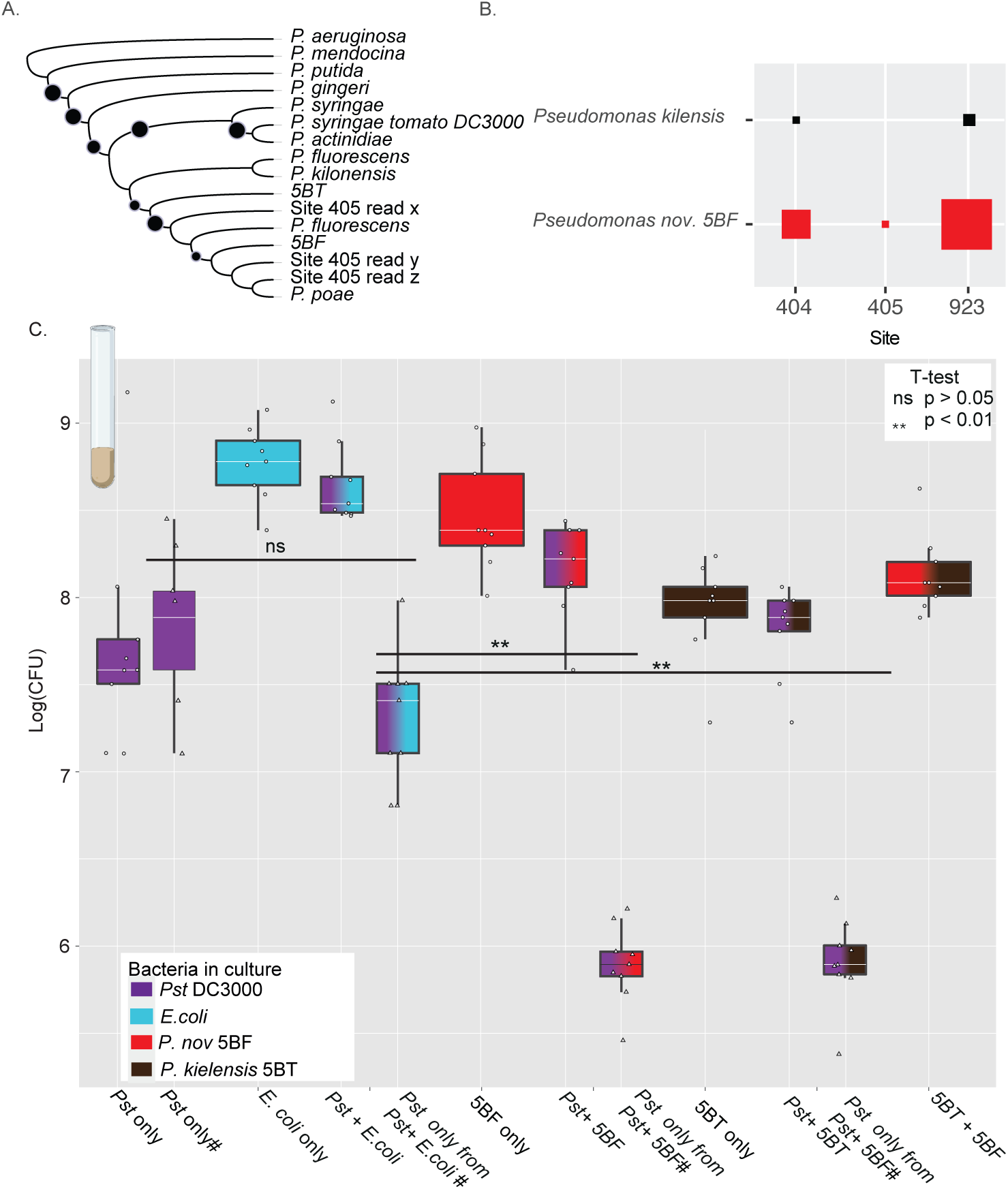
*In vitro* interactions between pseudomonads in pond samples and model bacterial species. A. Maximum likelihood phylogeny showing relationship of read hits at site 405. Dots indicate bootstrap values greater than 70. B. Number of reads to *P. nov* 5BF and *P. kilensis* based on SNP matrix.C. Bacterial colony forming units on LB with no antibiotics or LB supplemented with rifampicin and kanamycin to select for *Pst* DC3000 as indicated by #, the only isolate resistant to both antibiotics. P-values shown are from t-test are after Holm-Bonferroni adjustment ((n = 4 for each treatment condition, for each biological replicate in ‘n’ three technical replicates were performed) Table S20)((n = 4 for each treatment condition, for each biological replicate in ‘n’ three technical replicates were performed) Table S20)

### *In vitro* liquid culture inoculation experiments reveal negative growth effect of environmental isolates on *Pst* DC3000

To study the interaction between *Pst* DC3000 and 5BF further, we investigated the potential for growth inhibition in *in vitro* co-cultures. *Escherichia coli* was included as a negative control as it should not inhibit *Pst* DC3000 growth. *E. coli* co-culture with *Pst* DC3000 did not cause a significant reduction in *Pst* DC3000 colony forming units (CFU) after 24 hours (t-test, Holm–Bonferroni adjusted p = 0.0798) (Fig 7, Table S20). However, co-culture with either *P.* nov. 5BF or *P. kielensis 5*BT under similar conditions resulted in a significant reduction in CFU of *Pst* DC3000 (t-test, Holm–Bonferroni adjusted p-values: 0.00346, 0.00061). The reduction in CFU upon co-culture of *Pst* DC3000 with *P.* nov. 5BF or *P. kielensis* 5BT suggests there could be interference competition between *Pst* DC3000 and *P.* nov. 5BF, as well as *P. kielensis* 5BT.

**Figure 7.**
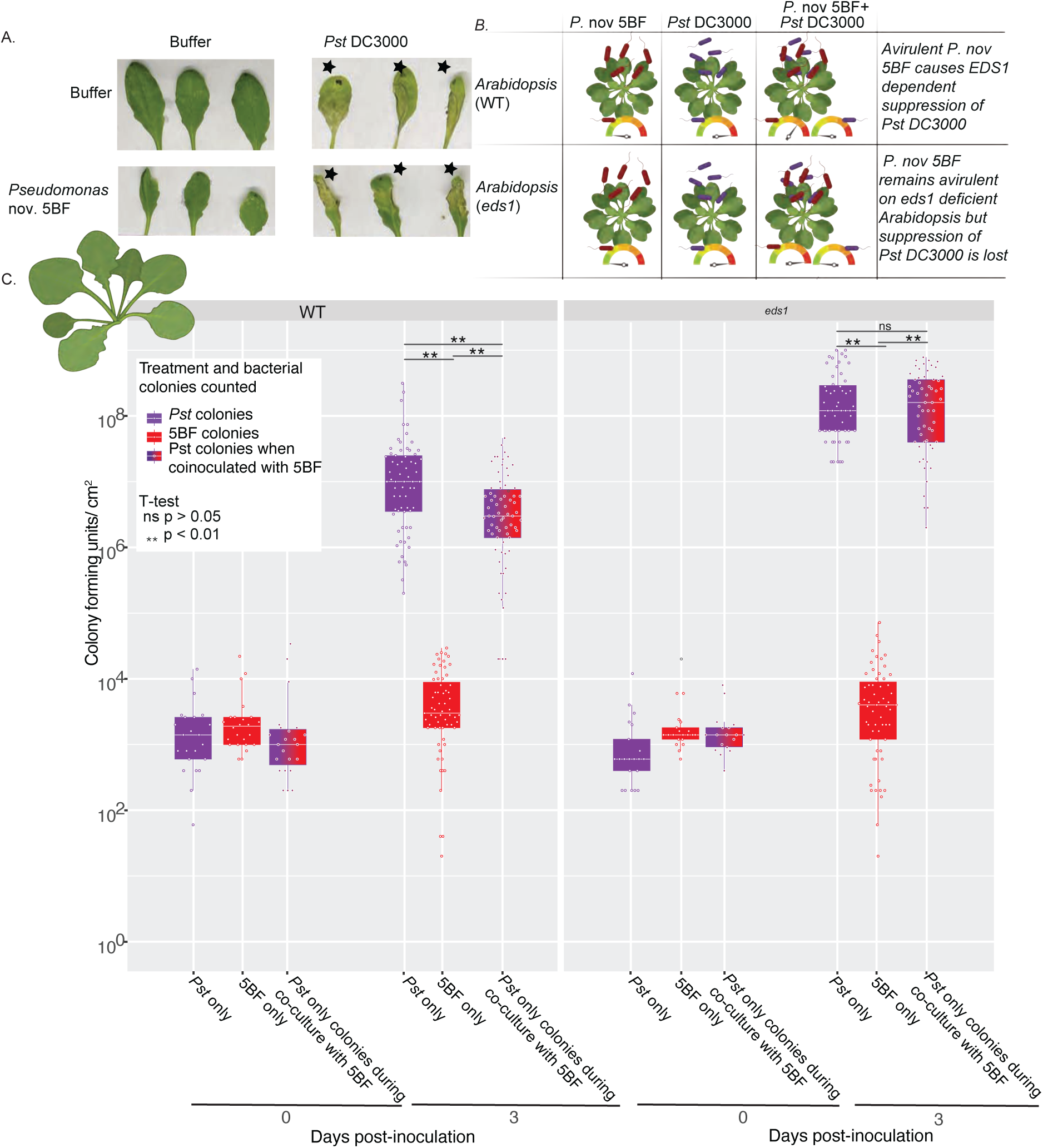
*Arabidopsis thaliana* after inoculation with *Pseudomonas spp*. A. Images of the three leaves per plant 48 hours after syringe infiltration with a standard high inoculum bacterial solution. Black stars indicate leaves that appeared to fully collapse. B. Quantification of CFU of *Pseudomonas spp*. in *A. thaliana* WT and eds1 knockout line when inoculated alone and together. P-values shown are from t-test after Holm-Bonferroni adjustment ((n=7 for WT plants each treatment condition and n=6 for each eds1 treatment condition, for each biological replicate in ‘n’ three technical replicates wereperformed) Table S21).

Given the potential for co-localization in the shared environment within the plant, we next assessed contact-dependent interactions of both isolates through a cross-streak assay (Fig S28). *P.* nov. 5BF and *P. kielensis* 5BT each showed a pattern of growth along the juxtaposed *Pst* DC3000 colony streak, irrespective of which strain was streaked first. In addition, there was no zone of clearing between 5BF or 5BT and *Pst* DC3000. In contrast, cross streaks of *P.* nov. 5BF and *P. kielensis* 5BT showed an absence of colony establishment at the intersection of the streaks, a pattern consistent with antimicrobial activity.

### *A. thaliana* does not support 5BF growth resulting in only minor EDS-1 dependent growth suppression of *Pst* DC3000

To further interrogate the mechanisms of disease protection provided by *P.* nov. 5BF and *P. kielensis* 5BT, we decided to test the strains on *A. thaliana*. When infiltrated with a standard high bacterial load of *P.* nov. 5BF and *P. kielensis* 5BT, *A. thaliana* leaves showed signs of small yellow lesions (Fig 8a). To test whether the disease inhibition of *Pst* DC3000 could be replicated in *A. thaliana*, we then inoculated leaves by syringe infiltration with *P.* nov. 5BF and *Pst* DC3000. In contrast to *S. polyrhiza*, we saw no suppression of disease symptoms; instead, we observed leaf collapse for leaves infiltrated with *P.* nov. 5BF and *Pst* DC3000. Often, *Pst* DC3000 infiltrations alone showed lower symptoms at 48 hours post-inoculation than when co-inoculated with *P. nov.* 5BF (Fig. 8a, Fig S29-32).

To understand the dynamics of *Pst* DC3000 and *P.* nov. 5BF interaction in *A. thalina*, we performed growth points at 3 dpi, following syringe infiltration of leaves with a standard low bacterial inoculation. In contrast to *S. polyrhiza*, following inoculation of *P.* nov. 5BF alone, *P.* nov. 5BF populations did not grow beyond the level at which it was infiltrated into *A. thaliana* by at 3 dpi. *Pst* DC3000 was able to grow to much higher levels than *P.* nov. 5BF in the leaf (Fig 7b, Fig S33, Table S21). Surprisingly, given the increased severity of symptoms upon high inoculum co-inoculation of *Pst* DC3000 and *P.* nov 5BF, we observed a small but significant (t-test, Holm-Bonferroni adjusted p ≤ <0.01) decrease in the number of *Pst* DC3000 colonies at 3 dpi compared with a low standard inoculum.

Furthermore, we were interested in whether the presence of the EDS1 pathway may play a role in the differential protection of *P.* nov. 5BF after co-inoculation with a high bacterial inoculum across species. To investigate this, we repeated the growth points in an *eds1* knockout line of *A. thaliana. P.* nov. 5BF was still unable to grow above starting inoculum levels in the *eds1* mutant plants as in wild-type (Fig 7, Fig S33, Table S21). The absence of *EDS1* alone is not sufficient to explain the difference between *S. polyrhiza* and *A. thaliana* in the growth of *P.* nov. 5BF, as in *A. thaliana eds1 P.* nov. 5BF was unable to grow to high density and did not significantly suppress *Pst* DC3000 CFU/cm^2^. Co-inoculation with *P.* nov 5BF in the *eds1* knockout did not suppress the growth of *Pst* DC3000, suggesting EDS1-dependent interaction in Arabidopsis unlike the suppression seen between *P. nov* 5BF and *Pst* DC3000 in duckweed and *in vitro*.

## Discussion

Here, we characterized the *Landoltia sp.* microbiome at three different locations within 1km of each other. Analysis of the pond water filtrate showed a differential protective effect produced by the 0.2 μm filtrate. From this filtrate we were able to identify two isolates with disease protective effects, which, through 16S rDNA and full genome sequencing, we revealed were *Pseudomonas* species. Genome analysis highlighted the presence of numerous bacterial interference competition genomic islands including type VI secretion systems and viscosin antimicrobial lipopeptide. *In vitro* co-cultivation showed direct inhibition of the number of *Pst* DC3000 colony forming units by *P.* nov. 5BF. However, the suppression of *Pst* DC3000 infection was not apparent in *A. thaliana eds1* mutants.

Several *Pseudomonas* species have been established as biocontrol agents due to their ability to suppress pathogen infection, either directly or through induced systemic resistance (ISR) (Cheng et al., 2017; Haney et al., 2018; Pacheco-Moreno et al., 2021). In contrast to site 405 0.2 μm filtrate, filtrates from sites 404 and 923 provided little or no protection against *Pst* DC3000 symptoms. The variation in disease suppression between pond filtrates could be considered similar to terrestrial plant disease-suppressive soils (Pacheco-Moreno et al., 2021; Schlatter et al., 2017). Despite the variation in disease-suppressive pond filtrate, Lemnaceae fronds from all sites appeared healthy. This could be for numerous reasons, such as the insufficient recovery of protective bacterial isolates by our methods, larger protective isolates being unfiltrable through the 0.2 μm filter, or protective bacteria’s incapability for disease protection *in vitro*.

Protection from disease can occur when specific members of the plant microbiome trigger host systemic resistance mechanisms such as systemic acquired resistance (SAR) and ISR (Cheng et al., 2017; Duke et al., 2017). Mechanisms such as SAR and ISR require the plant host to rapidly produce diffusible molecules to trigger defense responses upon infection (Duke et al., 2017; Fu & Dong, 2013; Pieterse et al., 2014). However, the definitions of SAR and ISR immunity cannot easily be applied to the duckweed Lemnaceae family given their reduced immune system, clonal reproduction, and body plan consisting of a frond and roots. Furthermore, studies would be needed to establish a role of SAR and ISR in *P.* nov. 5BF-induced resistance. However, if ISR or SAR is involved in disease protection, it is unlikely to be through SA upregulation since it has been shown that exogenous SA does not protect *S. polyrhiza* from *Pst* DC3000 infection (Baggs et al. 2022). A number of *Pseudomonas* species have previously been suggested as biocontrol agents (Bano & Musarrat, 2003; Duke et al., 2017; Ligon et al., 2000). However, it remains to be seen if disease protection by P. nov. 5BF would be transferable beyond Lemnaceae. Studies have shown that PGPB growth promotion can work upon inoculation of both *Lemna minor* and the dicot plant lettuce (Suzuki et al., 2014). The absence of protection against disease upon *P.* nov. 5BF co-inoculation with *Pst* DC3000 on *A. thaliana* plants and only minor growth suppression suggests that *Pseudomonas* nov. 5BF proliferation is required for the levels of antagonism with *Pst* DC3000 observed in duckweed. It remains unclear if the mechanism of protection would be the same or different from those established by the commensal disease-protective *Pseudomonas* sp. of *A. thaliana* (Shalev et al., 2022).

In contrast to ISR and SAR, disease protection can also be a byproduct of direct bacteria interaction (Bano & Musarrat, 2003; Harting et al., 2021; Ligon et al., 2000; Pacheco-Moreno et al., 2021; Sun et al., 2017)). The results of our *in vitro* co-inoculation experiments suggest that in the tested environments, *P. kielensis* 5BT and *P.* nov. 5BF can suppress the growth of *Pst* DC3000. A previous study of the Lemnaceae microbiome provided evidence of preferential recruitment of bacterial isolates, which were hypothesized to ameliorate plant stressors (Acosta et al., 2020; O’Brien et al., 2022). Further studies would be needed to test whether *P.* nov. 5BF or *P. kielensis* 5BT is preferentially recruited to protect Lemnaceae from pathogen stress.

Microbiome characterization of closely located Lemnaceae populations at the UC Berkeley Botanical Garden revealed bacterial taxa had a more stable abundance relative to the more dynamic composition of the fungal microbiome. Consistent with prior work, the Lemnaceae bacterial microbiome was composed of similar taxa to those present in terrestrial plants (Acosta et al., 2020; O’Brien et al., 2020). Members of Comamonadaceae, the most abundant family associated with the Lemnaceae in this study, have been previously identified as core taxa in groundwater samples (Deja-Sikora et al., 2019; Inoue et al., 2022) and are enriched in disease-protective rhizospheres (Wen et al., 2020). Sphingomonadaceae include common members of terrestrial plant microbiomes, with species within the family acting as plant growth-promoting bacteria (Innerebner et al., 2011; Luo et al., 2019). There were numerous amplicons from *Pseudomonas* species, despite fronds appearing healthy and previous work showing a number of *Pseudomonas* pathogens capable of infecting sterile duckweed. We were able to identify 16S rDNA amplicon sequences that likely derived from the same species as *P.* nov. 5BF and *P. kielensis* 5BT. The fungal microbiome is more distinct from terrestrial plants with a higher proportion of the duckweed microbiome consisting of Chytridomycota (Almario et al., 2017; Bergelson et al., 2019; Fuentes et al., 2020). The higher abundance of Chytridomycota in an aquatic environment is not surprising given their presence typically in freshwater and marine habitats. However, it is interesting to consider how the increased abundance of this phyla may affect interactions within the microbiome and with the plant. Further experiments would be required to understand the relationship between duckweed and these fungal genera, though this data provides a starting point for further exploration of fungal pathogens that could be studied using the duckweed model system.

It remains unclear how duckweed’s association with isolated bacteria is established and the subsequent mechanism that facilitates increased resistance among duckweed fronds. Interestingly, the pond from which *P. nov* 5BF *and P. kielensis* 5BT were isolated had a lower percentage of *Pseudomonas* than sites 404 and 923. For growth inhibition between *Pseudomonas* species, there are a number of established mechanisms, including bacteriocins, type VI secretion systems, and secondary metabolite productions. In the genome of *P.* nov. 5BF, we did not detect complete bacteriocins or bacteriophages known for targeting other *Pseudomonas* (Carim et al., 2021). However, we were able to detect the presence of two type VI secretion systems and 13 secondary metabolites that could mediate bacterial interaction. It would be interesting for future research to determine the molecular mechanism of *Pst* DC3000 inhibition by *P.* nov. 5BF as this understanding could be further leveraged for improving biocontrol systems in agriculture.

## Materials and Methods

### Sampling of healthy populations of duckweed to characterize associated bacteria

With the permission of the UC Berkeley Botanical Garden, we sampled three pond locations that had large Lemnaceae populations (site 404: 37° 52’ 25.2” N, 122° 14’ 15.3” W; site 405: 37° 52’ 25.0” N, 122° 14’ 15.4” W; site 923: 37° 52’ 31.9” N 122° 14’ 23.9” W). We collected surface pond water samples on two separate dates, February 27^th^, 2020, and November 3^rd^, 2020. Three 166 ml samples spread across each site were taken and combined into the sample for that location for both pond water and duckweed fronds. At each site five 50 ml tubes were filled with duckweed fronds. Similar to Acosta et al. 2020. (Acosta et al., 2020): the water sample from each bed was split into three 166 ml samples and passed through sterilized miracloth to remove solids, and subsequently passed through a 0.2 μM PES 500 mL filter (Corning 431097) to collect the majority of the microbes on the filters. The resulting 0.2 μM filtrate was used for priming assays and culturing of bacteria. The filters were then cut into fourths with a sterile scalpel and one-fourth was plated immediately on antibiotics to observe water-associated fungi while the rest of the filter was stored at −80°C. Duckweed samples were processed for culture-dependent and independent analyses as described below.

### Duckweed microbiome culture-dependent analyses

To recover bacterial isolates, 1 ml of 0.2 μm water filtrate from sites 404 and 405 was plated onto several LB plates and incubated at room temperature for several days. Visually distinct colonies were identified, and a single colony of each distinct morphology was re-streaked onto a fresh plate. This was repeated three times or until no distinct differences in colony morphology were visible. Single colony-derived cultures were given isolate identifiers that consisted of the last number of the bed identifier from which they were isolated, followed by a unique combination of letters. It is of note that some isolates showed pathogenicity after initial isolation (Fig. S7-9); however, this effect was lost upon cryostorage in glycerol. Single colony-derived cultures were assigned a genus using the 16S rDNA (V1-V9).

Fungal isolates were cultured either from the 0.2 μM PES filter that pond water was passed through or from surface sterilized duckweed populations. For each pond which consisted of three 166 ml samples, a total of six filters were plated. Two one-fourth sections of each 0.2 μM PES water filter were plated on individual Malt Extract Agar (MEA) plates containing kanamycin, gentamicin, and rifampicin. Fungal isolates growing from the edge of the filter were transferred to a new plate and morphology groups were assigned. Two isolates from each morphology group were assigned genera using 5.8S Fun - ITS4. To observe fungi in closer association to Lemnaceae, ∼3 mL of fronds from each pond were washed with sterilized water over miracloth and separated from debris. These were then split into three ∼1 ml aliquots and underwent a surface sterilization procedure similar to (Thomson & Dennis, 2013) in order to select for endophytic fungi. Plants were submerged in 10% v/v bleach for 30 seconds, transferred into 70% v/v ethanol for 30 seconds, washed with sterile water, and then placed on sterile miracloth in a biosafety cabinet to dry. Each side of every frond was then touched to the surface of a MEA plate containing kanamycin, gentamicin, and rifampicin as a control to make sure surface sterilization was complete. The duckweed fronds were then split into pieces with a sterile scalpel to expose the inside of the tissue. For each pond a total of 50 duckweed fronds were plated across 6 15 mm x 100 mm Petri dishes containing MEA media with kanamycin, gentamicin, and rifampin. Fungi that visually came out of internal tissue were then re-plated and morphology groups were assigned. Two isolates from each morphology group were assigned genera using 5.8S Fun - ITS4.

### Duckweed microbiome culture-independent sequencing and analyses

Fronds from the environmental pond sampling were separated from water using sterile miracloth and then rinsed with ddH_2_O. Fronds were then transferred to a falcon tube containing 40 ml of PBS buffer using sterilized tweezers. To wash the microbial cells off the leaves, sonication (frequency 40 kHz) was performed for 6 min in an ultrasonic cleaning bath, followed by shaking at 200 r min^-1^ for 20 min at 30°C. Sonication (frequency 40 kHz) was then continued for 3 min. The plant suspension was filtered through miracloth to remove fronds, and the suspension was filtered through a 0.1 µm membrane (CPLabSafety, FX-146-5113-RLS) by vacuum filtration. The filter membrane was removed and stored at −80°C until DNA extraction. Filters were divided into two equal parts using a sterile razor blade. One-half of the filter was used for bacterial DNA extraction using the FastDNA spin Kit (MPBio, 116540600), with the CLS-TC buffer. The extracted DNA was quantified using a Qubit fluorometer. DNA was present in all samples from the environmental duckweed samples, but DNA extractions from laboratory sterile *S. polyrhiza* gave results of nucleic acids below-detectable limits.

DNA extracts were aliquoted and diluted to 2.5ng in 5 µl of water for 16S rDNA amplicon PCR and barcode integration. To the DNA, 1.5 µl of ddH_2_O and 3 µl of each of the following primers at 2.5 µM concentration (Table S22) were added. 12.5 µl of 2X KAPA HiFi HotStart ReadyMix (Roche, 07958960001) was added before vortexing and spinning down. The PCR cycle conditions were as follows: initial denaturation at 95°C for 3 minutes, 27 cycles of denaturation at 95°C for 30 seconds, primer annealing (the ramp rare for the annealing step to ≤ 3°C/s.) at 57°C for 30 seconds, and template extension at 72°C for 60 seconds.

To amplify the ITS region for fungal classification 5.5 ul of ddH_2_O and 2 µl of each of the following primers at 2.5 µM concentration (Table S23) were added to the 3uL of DNA at a 4ng/uL concentration. 12.5 µl of 2X KAPA HiFi HotStart ReadyMix (Roche, 07958960001) was added before vortexing and spinning down. The PCR cycle conditions were as follows: initial denaturation at 95°C for 3 minutes, 34 cycles of denaturation at 95°C for 30 seconds, primer annealing (the ramp rate for the annealing step to ≤ 2.5°C/ s.) at 59°C for 30 seconds, and template extension at 72°C for 60 seconds (Taylor et al., 2016).

To check for DNA amplification, 1ul of the PCR amplified sample was run on a 0.75% agarose gel, and PCRs were then pooled together. Amplicons were size selected using 0.6X AMPure XP bead (Beckman Coulter, A63880) purification. For library preparation, the Nanopore Genomic DNA by ligation (SQK-LSK-110) - Flongle branch was followed.

The bioinformatic workflow used for oxford nanopore 16S and ITS rDNA amplicon sequencing is available on GitHub (https://github.com/erin-baggs/DuckweedMicrobes.git). In brief, the workflow consists of Fast5 files from nanopore that were converted to fasta using guppy Version 4.2.2+effbaf8, reads were demultiplexed using porechop v 0.2.4 (https://github.com/rrwick/Porechop) utilizing the custom barcodes (Table S23). Then an adapted version of the PuntSeq V1 protocol (https://github.com/d-j-k/puntseq version 1)(Urban et al., 2021) was used for read mapping and species abundance quantification for bacterial 16S and ITS rDNA.

For analysis of bacterial communities all reads were aligned to the SILVA database using minimap2. All reads that passed quality filters could be assigned classification to at least the class level (Table S24). The SRS R package (Beule & Karlovsky, 2020) was used to calculate diversity index values and downsample. Saturation of order, family, and taxa level richness required pooing the biological replicates together (Fig S20). Due to uneven numbers of reads per site, the read counts were normalized by scaling with ranked subsampling (Table S1-6) (Beule & Karlovsky, 2020). Some reads could be assigned to a site by the forward barcode but could not be demultiplexed to an individual replicate as reads were missing reverse barcodes. Among these reads for site 404 were reads assigned to *Xanthomonas* and *Streptomycetes*, whilst at 923 reads assigned as *Xanthomonas* were recovered. For *Xanthomonas* genera, reads were only identified from site 404 at very low relative abundance; the absence of a reverse barcode on *Xanthomonas* classified reads meant it could not be assigned to a particular replicate.

Taxonomic assignments for ITS rDNA amplicons (5.8S Fun - ITS4) were made by mapping with minimap2 against the UNITE database (Nilsson et al., 2019). Read processing and rarefaction were otherwise performed as with the 16S rDNA analysis. Due to uneven numbers of reads per site, the read counts were normalized by scaling with ranked subsampling (Table S8-9) (Beule & Karlovsky, 2020). Sub-sampling richness curves suggest that fungal genus taxa richness was saturated at all 3 sites after ∼80,000 reads respectively including the reads which do not map to the fungal kingdom (Fig S4). Reads that were assigned unidentified were removed as a majority of these represented non-fungal kingdoms (Table S25).

### Genome sequencing and assembly

Bacterial cultures were grown to OD_600_ = 0.6 in liquid culture at 28 °C within a shaking incubator (20.5 g). Monarch High Molecular Weight DNA Extraction Kit for tissue (T3060L) was used to extract DNA. For library preparation, Genomic DNA by ligation (SQK-LSK-110) - Flongle branch was followed. Resulting libraries were then run-on consecutive days on two flongle flow cells. Base-calling was done using guppy (Version 4.2.2+effbaf8). Assembly was constructed using flye v2.8.3-b1695 on read fasta files (Kolmogorov et al., 2020). Medaka v.1.0.3 was used for genome polishing (https://github.com/nanoporetech/medaka). This was confirmed using orthoANI (Lee et al., 2016) to assess the similarity of isolate 5BF to *Pseudomonas poae,* the bacterial species with the highest similarity across the 16S rDNA locus.Genome assemblies were submitted to The National Center for Biotechnology Information Genbank and TrueBac™ID (Ha et al., 2019), both identified the 5BF isolate to be a novel *Pseudomonas* specie and 5BT as *Pseudomonas kielensis*.

Genome assembly statistics and annotation was performed utilizing PATRIC v3.22.0 (Brettin et al., 2015). BUSCO version 5.3.0 with the bacteria_odb10 database was used to assess the quality of genome assembly. The *Pst* DC3000 (NC_004578.1) (Buell et al., 2003) and *Pss* B728a (NC_007005.1) (Feil et al., 2005) genomes used for comparison were downloaded on 11/27/21 from NCBI. Type VI Secretion System prediction was carried out using SecReT6 v.3 (Li et al., 2015) prediction on the nucleotide genome assembly (Li et al., 2015). BLASTN of the two T6SS operons from *P.* nov. 5BF against the NCBI nucleotide collection (nr/nt) showed the operons that displayed homology to other *Pseudomonas* species T6SS. To identify secondary product genomic clusters antiSMASH5.0 was used (Blin et al., 2019), and Phaster v. Dec 2022 (Arndt et al., 2016) was used to identify phage in the genome (Arndt et al., 2016).

### Bacterial 16S PCR genotyping

Not all isolates survived well upon serial passaging. For colonies we were able to successfully isolate, a single bacterial colony was resuspended in 6.5 µl of H_2_O. Added to this was 3ul of each of the following primers at 2.5 µM concentration (Fwd p8-16S_For_bc1005: /5Phos/GCATCCACTCGACTCTCGCGTAGRGTTYGATYMTGGCTCAG, Reverse p16-16S_Rev_bc1033 /5Phos/GCATCAGAGACTGCGACGAGARGYTACCTTGTTACGACTT). 12.5 µl of 2X KAPA HiFi HotStart ReadyMix (Roche, 07958960001) was added before vortexing and spinning down. The PCR cycle conditions were as follows: initial denaturation at 95°C for 3 minutes, 37 cycles of denaturation at 95°C for 30 seconds, primer annealing (the ramp rate for the annealing step to ≤ 3°C per second) at 57°C for 30 seconds, and extension at 72°C for 60 seconds. PCR product was run on a 0.75% agarose gel to confirm amplification. PCR product excised from gel was extracted using Zymoclean gel DNA recovery kit (Zymogen, D4007) and mixed with primers before Sanger sequencing at the UC Berkeley DNA Sanger Sequencing Facility.

### Duckweed sterilization and genotyping

Duckweed fronds collected from the ponds were sterilized by submerging in 10% bleach and agitating on a rotor for ∼2 min until whitening was visible along the frond edges. The fronds were then rinsed with sterile deionized water three times and transferred to fresh Schenk and Hildebrandt basal salt media. The fronds were allowed to regenerate over 2 months in the growth chamber under the conditions listed above. Only fronds from site 405 survived sterilization. DNA was extracted from new growth using Plant DNAeasy extraction kit (Qiagen, 69104). For genotyping, 100ng of DNA was resuspended in 10 µl of H_2_O, followed by PCR (GoTaq 5x Buffer 10 μl, 10 mM dNTPs 1.25 μl, 10 μM Primer 1 1.25 μl, 10 μM Primer 2 1.25 μl, MgCl2 6.25 μl, H2O 19.75 μl, GoTaq 0.25 μl). The following primers were used: aptF-atpH-Fwd ACTCGCACACACTCCCTTTCC, aptF-aptH-Rev GCTTTTATGGAAGCTTTAACAAT (Wang, et al 2010). PCR conditions were as follows: initial denaturation at 95°C for 2 minutes, 42 cycles of denaturation at 95°C for 1 minute, annealing at 53°C for 1 minute, extension at 72°C for 0.5 minutes, and 1 cycle final extension at 72°C for 5 minutes. PCR product was run on a 0.75% agarose gel to check for amplification. PCR product excised from gel was extracted using Zymoclean gel DNA recovery kit (Zymogen, D4007) and mixed with primers before sanger sequencing at the UC Berkeley DNA Sanger Sequencing Facility.

### Plant growth conditions

Duckweed fronds are propagated every 3-4 weeks by transferring 3 mother fronds to a new well with fresh media from a stock plate. Schenk and Hildebrandt basal salt media (Sigma-Aldrich, S6765-10L) (0.8% agarose, pH 6.5) was used to grow plants in 6 or 12 well plates. Plates were then transferred to a growth chamber set to 23 °C with a diurnal cycle of (16hr light (400 lux)/ 8hr dark). *A. thaliana* was grown in sunshine mix 4 in a growth chamber set to 23 °C/19 °C with a diurnal cycle of (16hr light (300 lux)/ 8hr dark).

### Duckweed priming

Sterile *S. polyrhiza* was primed by placing around 30 duckweed fronds into a 50ml beaker with around 10 ml of 0.2 µM rifampicin or 0.1µM PES filtered pond water (and in one case boiled filtered pond water) from the three separate ponds and sterile PBS for the negative control. The duckweed fronds remained suspended in pond water for 24 hours at room temperature and ambient light. Primed duckweed was then transferred onto a sterile miracloth taped over a beaker by pouring the entire solution over the miracloth. Using pipette tips, exactly 5 fronds were transferred into each well of a six well plate containing Schenk and Hildebrandt basal salt media (Sigma-Aldrich, S6765-10L) (0.8% agarose, pH 6.5). The duckweed was then inoculated using vacuum infiltration at 80 PSI with the pathogen *Pst* DC3000, or a *Pst* DC3000 *hrcC* mutant as described in the pathogen inoculation section below.

### Pathogen inoculation

Bacterial strains were grown in LB media supplemented with the following concentrations of antibiotics: *P. syringae* pv. tomato DC3000 (Matthysse et al., 1996; Mudgett & Staskawicz, 1999) with 10 μg/ml; kanamycin (Km) and 50 μg/ml; rifampicin (Rif), P. *syringae* pv. tomato DC3000 dsRed (Rufián et al., 2022) with Rif 50 μg/ml and 20 μg/ml; gentamicin (Gm) and *P. syringae* pv. syringae B728a (Feil et al., 2005) with Rif 50 μg/ml. Plates were washed with 1ml of 10mM MgCl_2_ to remove bacteria. The OD_600_ was then measured and adjusted to OD_600_ = 0.1 with 10mM MgCl_2_.

### Growth curves

For duckweed standard high inoculum experiments, a total of 500 µL of bacterial inoculation solution (OD_600_ = 0.1) was pipetted onto three duckweed fronds. The plate was then either placed straight into the incubator or placed in a vacuum (0.8 PSI) for 10 minutes.

*In vitro* cultures were started with a single colony of each bacterium and allowed to grow in LB with no selection. Cultures were sampled after 24 hours, serial dilutions were made of cultures, and dilutions were plated on LB without antibiotics to quantify total bacterial growth and on antibiotic media to select for *Pst* DC3000 alone.

For standard high inoculum experiments with *A. thaliana*, the leaves were syringe-infiltrated with OD_600_ = 0.1 solution until water soaking was visible across the full leaf surface. For standard low inoculum experiments, 25 ul of OD_600_ = 0.1 solution was diluted in 25ml of 10mM MgCl_2_, infiltrations were then carried out as above with standard high inoculum. On day 0 and day 3, leaves and fronds were harvested, photographed, and sectioned. The tissue was then homogenized with 3mm glass beads in a biospec mini-beadbeater at 2,000 rpm with a vital distance of 1.25 inches. Serial dilutions were made and plated on appropriate selective media. Two days after plating, colonies were counted.

### Microscopy

For microscopy, flood inoculation with 500 ul solutions of inoculum (OD_600_ = 0.1) was used. Whole fronds were staged in water on slides and covered with a glass coverslip at 7 dpi. The slides were imaged on a Zeiss 710 LSM confocal microscope with either the 20x (water), 63x (oil), or 100x (oil) objectives. To image bacteria on duckweed fronds, bacteria were stained with 10ul of 1x Syto™ BC Green Fluorescent Nucleic Acid Stain (Thermo Scientific, S34855). To further investigate whether *Pst* DC3000 and *P. nov*. 5BF occupy the same niche, we used *Pst* DC3000 dsRed (Rufián et al., 2022) and SytoBC-stain for bacteria in general. Utilizing the Zeiss LSM900 with Airyscan 2 and then Airy Process using Zen Blue, we were able to identify *P.* nov. 5BF and *Pst* DC3000 in the same field of view, though the *Pst* DC3000 appeared embedded in crypts whilst *P.* nov. 5BF was on the surface. The Syto™BC-stain was unable to penetrate the crypt regions as can be seen by the absence of overlapping fluorescent labeling of *Pst* DC3000 that should also be stained by Syto™BC.

### 16S rDNA alignments for species designation

Given the accuracy of 98.3% per bp of the R9.4.1 flow cells used for sequencing, we expected that within our ∼1,400 bp sequences those with fewer than 23 nucleotide mismatches could correspond to reference 16S rDNA for strains of interest. We used BLASTn to identify reads that passed the threshold of nanopore error corresponding to other known *Pseudomonas* species including pathogens, endophytes, and biological controls (File S2). We were unable to recover any reads that passed the threshold for 9 *Pseudomonas* species including *Pst* DC3000 and *Pseudomonas aeruginosa*. At site 923, reads were recovered with high similarity to *Pseudomonas fluorescens, a species* that includes known plant pathogenic and protective strains. We were able to recover reads from sites 404 and 923 that passed the threshold of similarity to isolate *P. kielensis* using *P. kielensis* 5BT as reference.

To identify the presence of bacteria of the same species as *P.* nov. 5BF required further filtering due to the high similarity of 16S rDNA of *P.* nov. 5BF and *P. poae.* After the initial filtering, the same reads were often mapped to both species. To distinguish the two species, we performed RNA structure-informed alignment using SSU-align (Nawrocki, 2009) of the 16S rDNA region reads and reference sequences (File S3). The alignments were then inspected for SNP differences between *P. poae* and *P.* nov 5BF. The double-mapped amplicon reads which mapped to both *P. poae* and *P.* nov 5BF sequences were checked as to whether the SNPs follow *P.* nov 5BF’s sequences (Table S14,15). For 7 of 8 positions at which there were reference-specific SNPs, the majority of the sequences shared nucleotides with *P.* nov 5BF. The majority of SNPs between reads and *P.* nov 5BF reference sequences were low frequency and even those found in more than 4/37 reads were widely distributed across variable and conserved regions of the 16S rDNA gene (Fig S21). The pattern of distribution of SNPs between and among sequences is largely characteristic of technical errors. The recovery of reads with high identity to *P.* nov 5BF and *P. kielensis* 5BT 16S rDNA, suggests the species are ubiquitous across the sampled sites at the UC Berkeley Botanical Garden.

## Author contributions

E.L.B. and F.G.S designed and undertook the sampling at the UC Berkeley Botanical Garden. F.G.S conducted pond water filtration. E.L.B and F.G.S conducted phytopathogen assays. E.L.B isolated bacteria from environmental samples. E.L.B and M.B.T conducted *L. punctata* microbiome DNA extraction. E.L.B and F.G.S undertook library preparation, amplicon sequencing and amplicon sequencing analysis. E.L.B undertook bacterial growth curves and was responsible for DNA extraction, library preparation, sequencing, and annotation of the 5BF and 5BT genomes. E.L.B, F.G.S and K.V.K designed the study. E.L.B prepared figures and wrote a complete draft of the manuscript. All authors contributed to writing and editing of the final manuscript.

## Data availability

Sequencing reads for 16S, and ITS rDNA and microbial genomes can be found within the NCBI BioProjects PRJNA785658. The genomes and annotation of *Pseudomonas* nov. 5BF and Pseudomonas *kielensis* 5BT are available through the IMG/JGI database with the respective IMG IDs 2931463386 and 2935452895.

## Acknowledgements

We would like to thank The UC Berkeley Botanical Garden for allowing us to sample the ponds in their grounds. We would also like to thank Dr. Denise Schichnes and Dr. Steven Ruzin for their expert advice and help with confocal microscopy. Research reported in this publication was supported in part by the National Institutes of Health (NIH) S10 program under award number 1S10RR026866-01. Many thanks to China Lunde for propagation of duckweed during the pandemic and Daniel ChoAhn for his work with the duckweed project team. We thank China Shaw, Wei Wei, Pierre Joubert, Kyungyong Seong, and Chandler Sutherland for their helpful comments and diligent reading of drafts. We thank Dr. Wilfried Haerty and all members of the Krasileva lab for helpful discussions. This research has been supported by the Gordon and Betty Moore Foundation (8802), the Innovative Genomics Institute, and the NIH Director’s Award (052239-002).

## Supplementary Figures

Figure S1. Rarefaction curves for 16S rDNA amplicons.

Figure S2. Bacterial family classification at the UC Botanical Garden sites.

Figure S3. Principal component analysis of bacterial microbiome composition at the UC Botanical Garden sites.

Figure S4. Rarefaction curves for fungal ITS rDNA amplicons.

Figure. S5 *Spirodela polyrhiza* priming with 0.2 μm water filtrate following pathogen inoculation.

Figure S6. *Spirodela polyrhiza* priming with 0.2 μm water filtrate before and after boiling followed by pathogen inoculation.

Figure S7. Experiment 2 *Spirodela polyrhiza* co-inoculated with bacteria isolated from the 0.2 μm water filtrate and *Pst* DC3000.

Figure S8 Experiment 3 *Spirodela polyrhiza* co-inoculated with bacteria isolated from the 0.2 μm water filtrate and *Pst* DC3000.

Figure S9. Experiment 4 *Spirodela polyrhiza* co-inoculated with bacteria isolated from the 0.2 μm water filtrate and *Pst* DC3000.

Figure S10. Experiment 5 *Spirodela polyrhiza* co-inoculated with bacteria isolated from the 0.2 μm water filtrate and *Pst* DC3000.

Figure S11. Experiment 1 *Landoltia punctata* 5635 inoculated with pathogens and environmental bacteria.

Figure S12. Experiment 2 *Landoltia punctata* 5635 inoculated with pathogens and environmental bacteria.

Figure S13. Experiment 3 *Landoltia punctata* 5635 inoculated with pathogens and environmental bacteria.

Figure S14. Experiment 1 Isolate 5BF’s protection of *L. punctata* BG405 from symptomatic Pss B728a infection.

Figure S15. Experiment 2 *Landoltia* BG405 inoculated with pathogens and environmental bacteria.

Figure S16. Experiment 2 *Landoltia* BG405 with pathogens and environmental bacteria.

Figure S17. Experiment 3 *Landoltia* BG405 with pathogens and environmental bacteria.

Figure S18. Circos plot of *Pseudomonas nov*. 5BF and *Pseudomonas kielensis* 5BT genome assemblies.

Figure S19. Pie-chart of the number of subsystems within superclasses for *Pseudomonas* species of interest.

Figure S20. Comparison of isolate *Pseudomonas nov.* 5BF type i1 T6SS with *Pst* DC3000 type i1 T6SS.

Figure S21. NCBI phylogeny of the top 100 BLASTN hits queried with the *Pseudomonas nov.* 5BF type i1 T6SS locus.

Figure S22 NCBI phylogeny of the top 50 BLASTN hits to the *Pseudomonas nov.* 5BF type i3 T6SS locus

Figure S23. Synteny between *Pseudomonas nov.* 5BF and *Pseudomonas fluorescens* SBW25 at the viscosin secondary metabolic islands

Figure S24. Confocal microscopy of Z-stack of 5BT on *Spirodela polyrhiza* fronds.

Figure S25. Growth curve split by experiment *Spirodela polyrhiza* co-inoculated with low bacterial load of *Pseudomonas nov.* 5BF and or *Pst* DC3000.

Figure S26. Experiments 1-5 *Spirodela polyrhiza* co-inoculated with low bacterial load of *Pseudomonas nov.* 5BF and or *Pst* DC3000.

Figure S27. Consensus sequence of nanopore reads and reference 16S rDNA sequence from secondary structure informed sequence alignment.

Figure S28. *Pseudomonas* interaction streak replicates.

Figure S29. High OD syringe infiltration of *A. thaliana* leaves with *Pseudomonas spp*.

Figure S30. High OD syringe infiltration of *A. thaliana* leaves with *Pseudomonas spp*.

Figure S31. High OD syringe infiltration of *A. thaliana* leaves with *Pseudomonas spp*.

Figure S32. High OD syringe infiltration of *A. thaliana* leaves with *Pseudomonas spp*.

Figure S33. Low OD syringe infiltration of *A. thaliana* leaves with *Pseudomonas spp*.

Figure S34. Model of bacterial plant interactions

## Supplementary Tables

Table S1 - Percentage abundance of orders across samples for 16S rDNA amplicon sequencing.

Table S2 - Percentage abundance of families across 16S rDNA amplicon sequencing.

Table S3 - Percentage abundance of genera across 16S rDNA amplicon sequencing

Table S4 - Phyla classification of ITS rDNA amplicons with unidentified reads included in relative abundance.

Table S5 - Phyla classification of ITS amplicons with unidentified reads removed.

Table S6 - Percentage abundance of orders across sites for ITS amplicon sequencing.

Table S7 - Culture dependent isolation, morphotyping and genotyping of fungi.

Table S8 - Percentage abundance of families across sites for ITS amplicon sequencing.

Table S9 - Percentage abundance of genus across sites for ITS amplicon sequencing.

Table S10 - Relative abundance of ITS amplicons corresponding to morphotyped fungal taxa.

Table S11 - Relative abundance of 16S rDNA amplicons associated with bacterial pathogens not including unidentified bacteria.

Table S12 - Relative abundance of ITS amplicons associated with fungal pathogens not including unidentified ITS sequences.

Table S13 - Isolates from pond water 16S rDNA best hit.

Table S14 - BUSCO genome scores.

Table S15 - Genomic features identified in duckweed associated isolates that play a role in interbacterial competition in other species.

Table S16 - Number of reads after normalization at sites belonging to *P. nov* 5BF and *P. kilensis* species.

Table S17 - Discriminatory SNP positions between 16S rDNA sequence of *P. nov* 5BF and *P. poae*.

Table S18 - Table of all sites were there is a SNP/indel in more than 3 reads within the *P. nov* 5BF and *P. poae* alignment

Table S19 - Colony forming unit counts for duckweed growth curves.

Table S20 - Colony forming unit counts for in vitro growth curves.

Table S21 - Colony forming unit counts for Arabidopsis thaliana growth curves.

Table S22 - Barcodes for 16S rDNA amplicon sequencing.

Table S23 - Barcodes for ITS rDNA amplicon sequencing.

Table S24 - Class classification across sites before SRS normalization.

Table S25 - UNITE database kingdom classification for ITS amplicons.

## Supplementary Files

File S1 - AtpF genotyping of site 405 duckweed isolate.

File S2 - 16S rDNA nucleotide sequences for bacterial reference isolates and *Pseudomonas* nov. 5BF and *Pseudomonas kielensis* 5BT.

File S3 - 16S rDNA nucleotide sequence alignment for *Pseudomonas* nov. 5BF and raw reads.

