## Supplemental Figures for "*Pseudomonas* isolates from ponds populated with duckweed prevent disease caused by pathogenic *Pseudomonas* species"

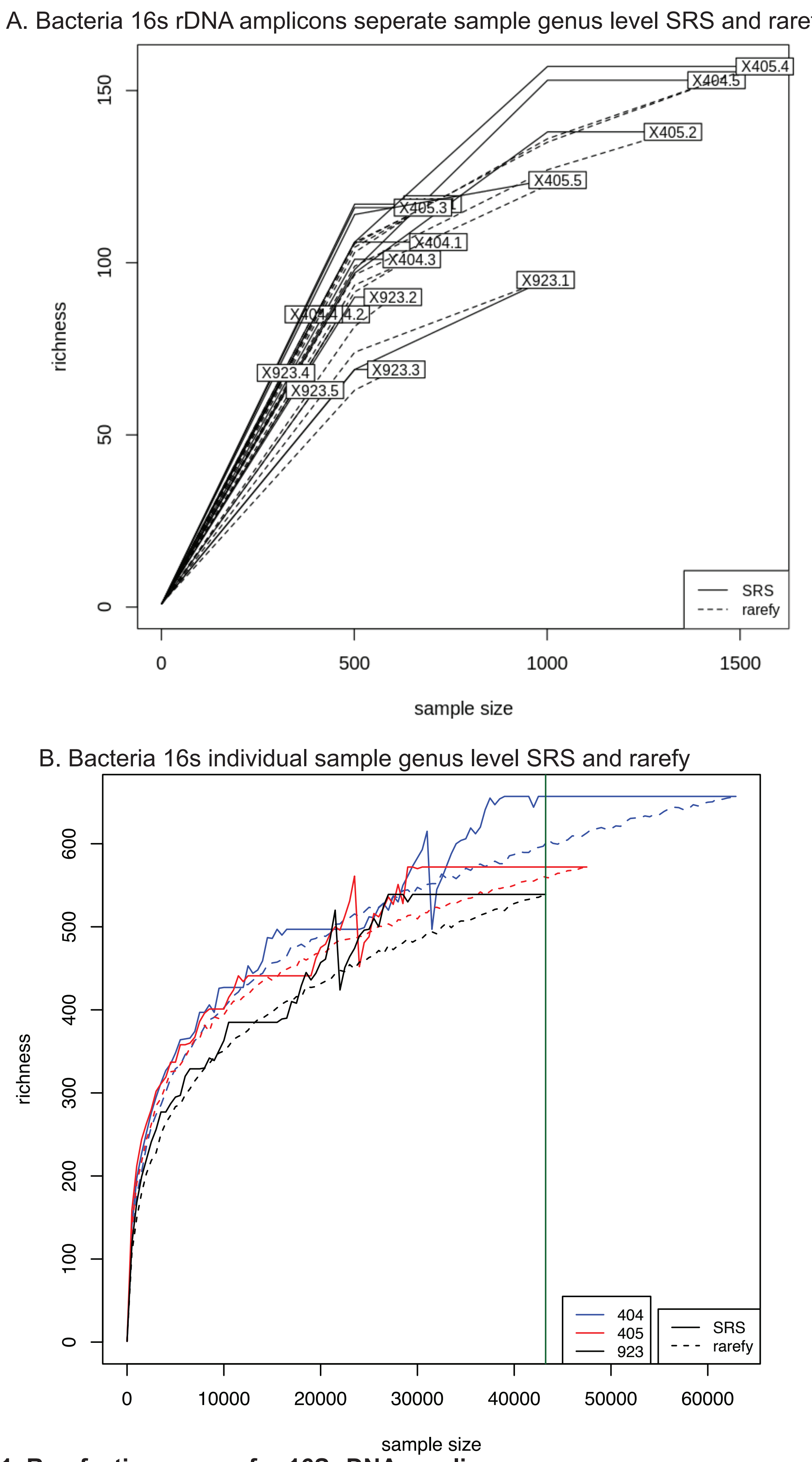


**Figure S1. Rarefaction curves for 16S rDNA amplicons.**

A. Bacteria 16s rDNA amplicons separate samples classified at genus level and subsampled with SRS and rarefy.B. Replicates from A are combined by site  Black- site 923, blue - site 404 and red - site 405. .

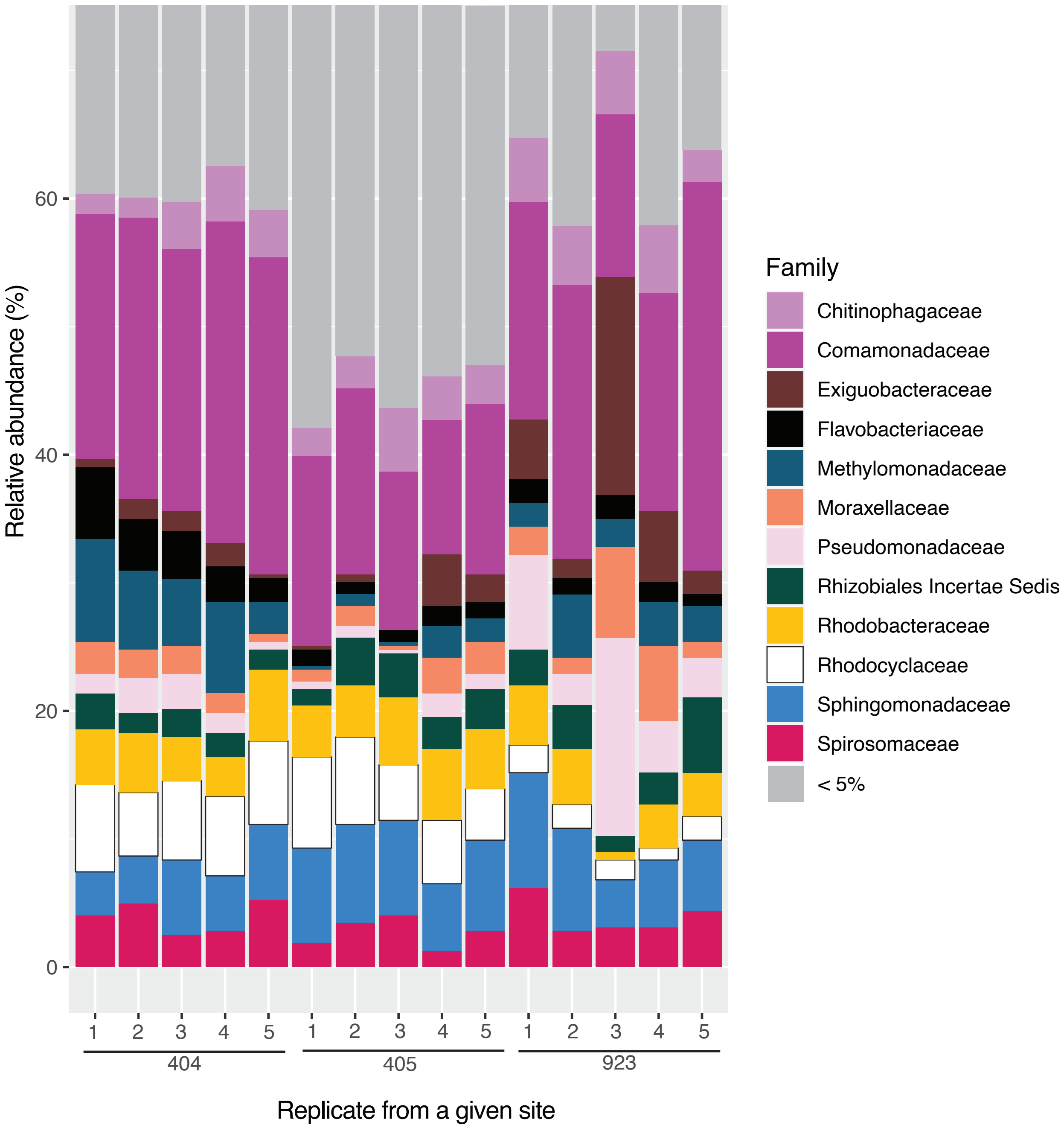


**Figure S2. Bacterial family classification at the UC Botanical Garden sites.**
A. Percentage of the 16S rDNA amplicons corresponding to SILVA 16S rRNA family classification for taxa which account for >1% of total classified reads for at least one site. The size of of the bar is proportional to percent of total classified 16S rDNA amplicons. B. Biplot of sites given bacterial order count data. C. As in B but upon genus level classification.

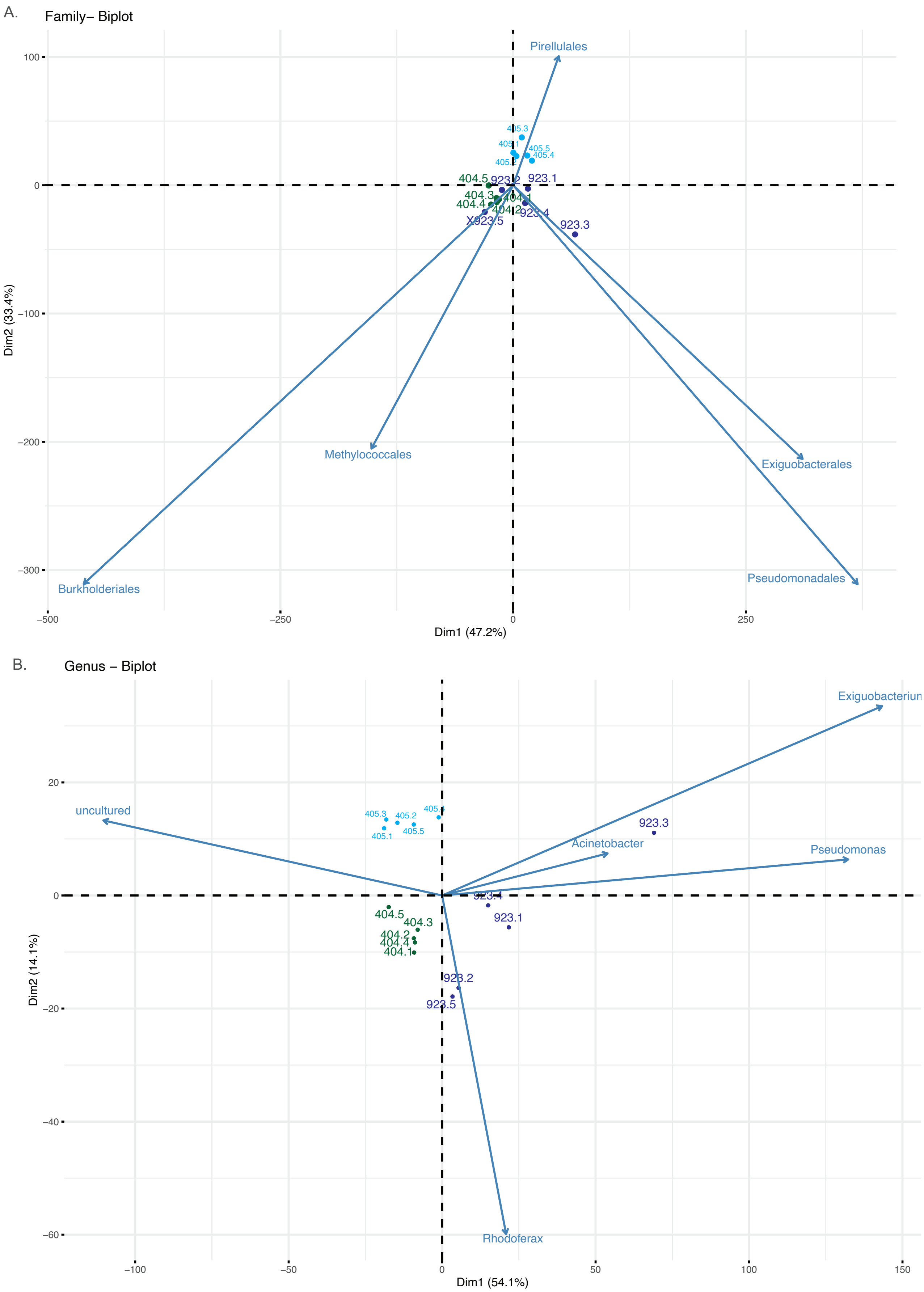


**Figure S3. Principal component analysis of bacterial microbiome composition at the UC Botanical Garden sites.**
Biplot of sites given bacterial family count data, arrows are shown for the top 5 components. Green - site 404, light blue - site 405 and dark blue - site 923 C. As in A but upon genus level classification counts.

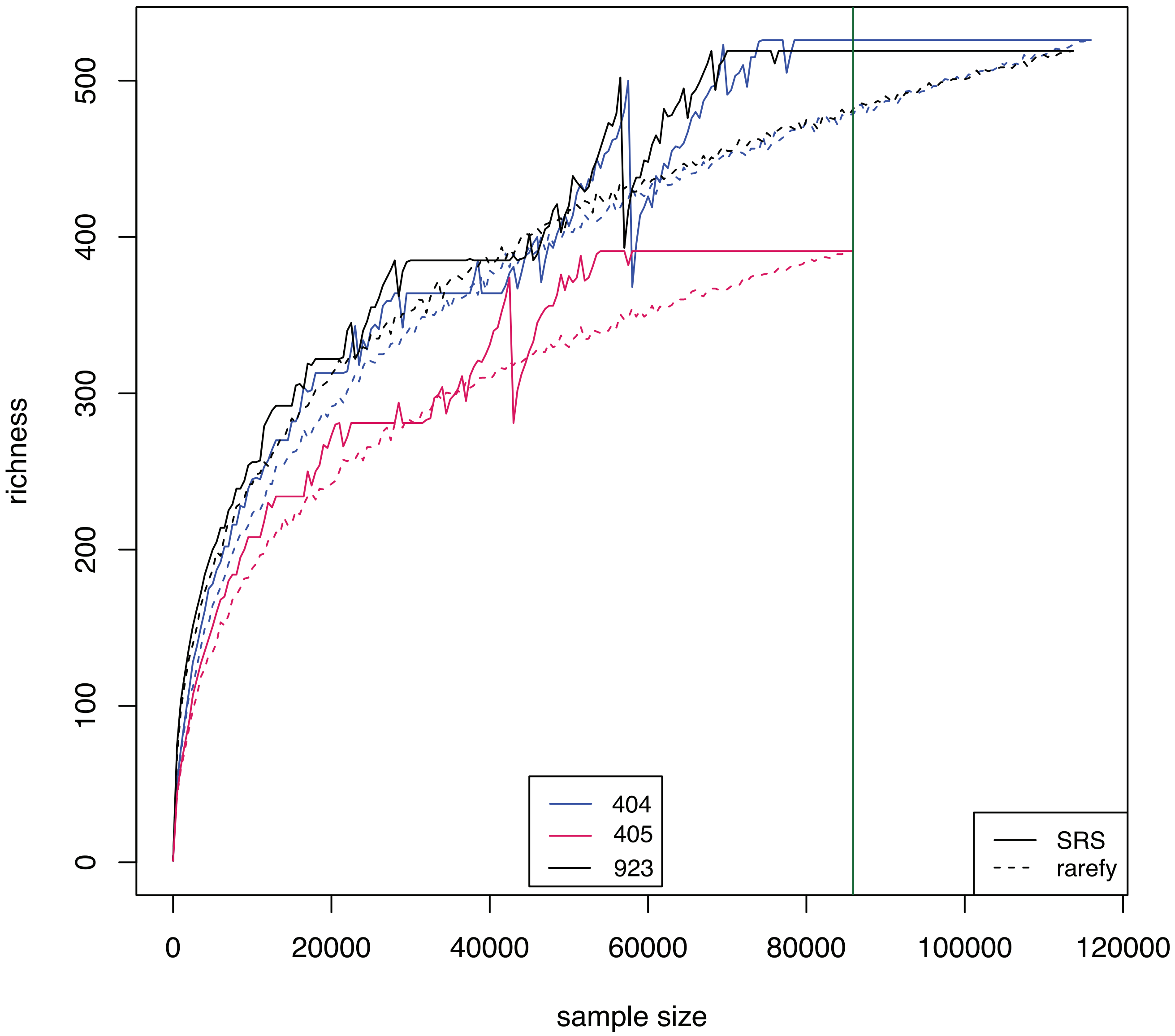


**Figure S4. Rarefaction curves for fungal  ITS amplicons.**
A. Fungal ITS amplicons separate samples pooled per site classified at genus level  and normalized with SRS and rarefy. Black- site 923, blue - site 404 and red - site 405.

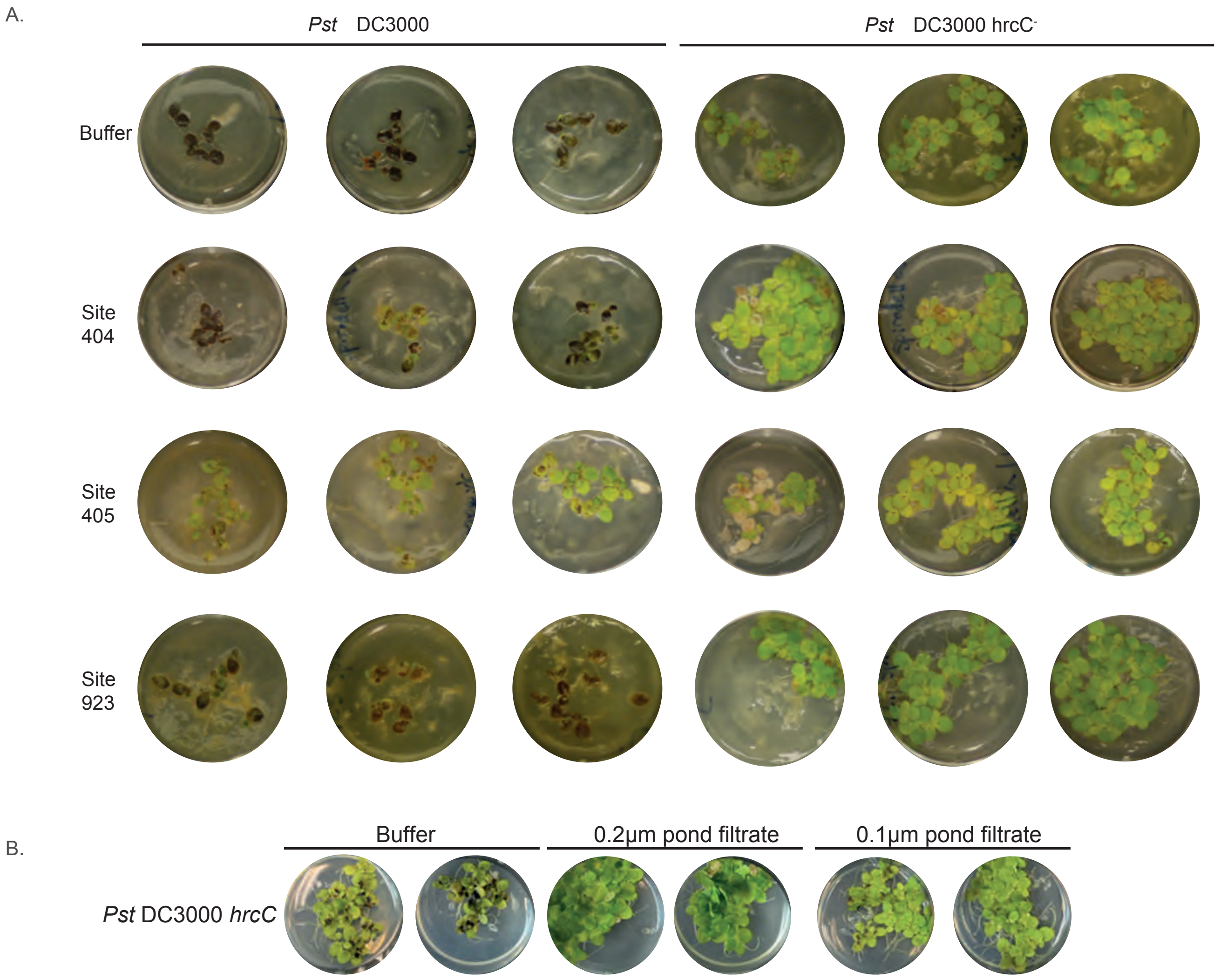


**Figure. S5 *S. polyrhiza* priming with 0.2 µm water filtrate following pathogen inoculation.**
A. Images 13 days post inoculation of duckweed primed for 24 hours in 0.2 µm filtrate from the three sites and subsequently treated with *Pst* DC3000 and *Pst* DC3000 hrcC-. B. Pond water 0.2 µm and 0.1 µm filtrate from site 405, two replicates are shown per treatment when inoculated with *Pst* DC3000 hrcC-.

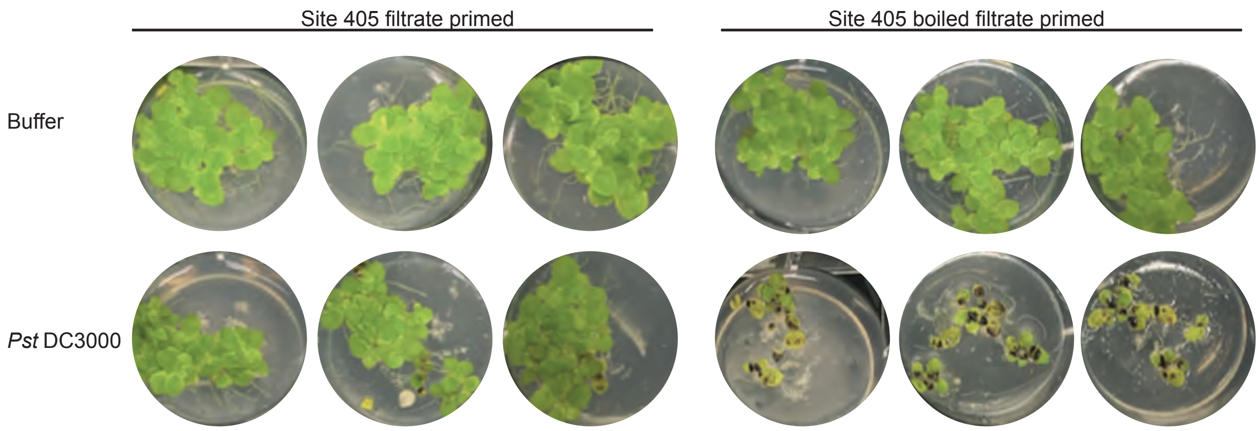


**Figure S6. *Spirodela polyrhiza* priming with 0.2 µm water filtrate before and after boiling followed by pathogen inoculation.**
Images 10 days post inoculation of duckweed primed for 24 hours in 0.2 µm filtrate from the site 405 with subsequent treatment with *Pst* DC3000 or 10mM MgCl2 buffer control.

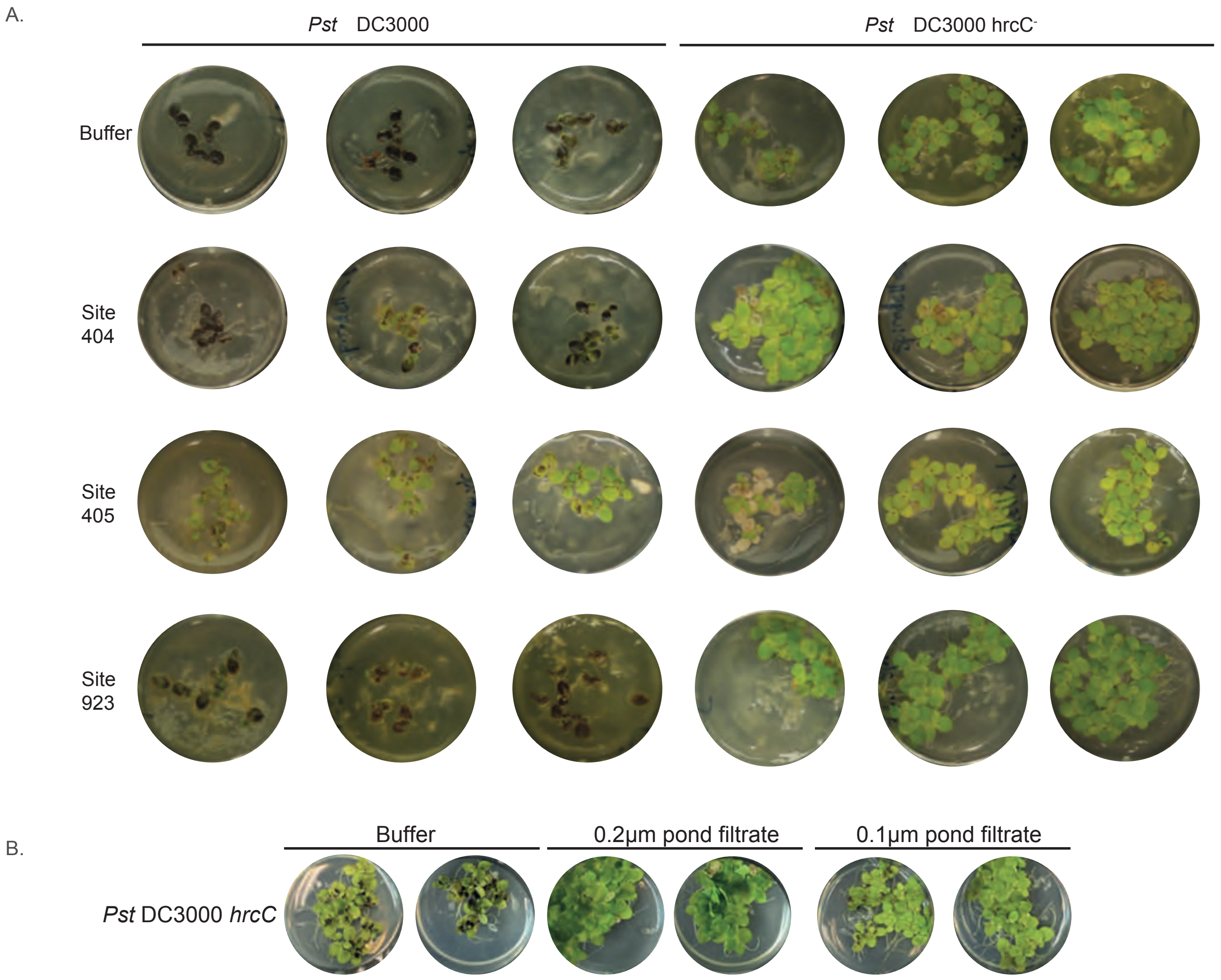


**Figure. S5 *S. polyrhiza* priming with 0.2 µm water filtrate following pathogen inoculation.**
A. Images 13 days post inoculation of duckweed primed for 24 hours in 0.2 µm filtrate from the three sites and subsequently treated with *Pst* DC3000 and *Pst* DC3000 $hrcC^{-}$. B. Pond water 0.2 µm and 0.1 µm filtrate from site 405, two replicates are shown per treatment when inoculated with *Pst* DC3000 $hrcC^{-}$.

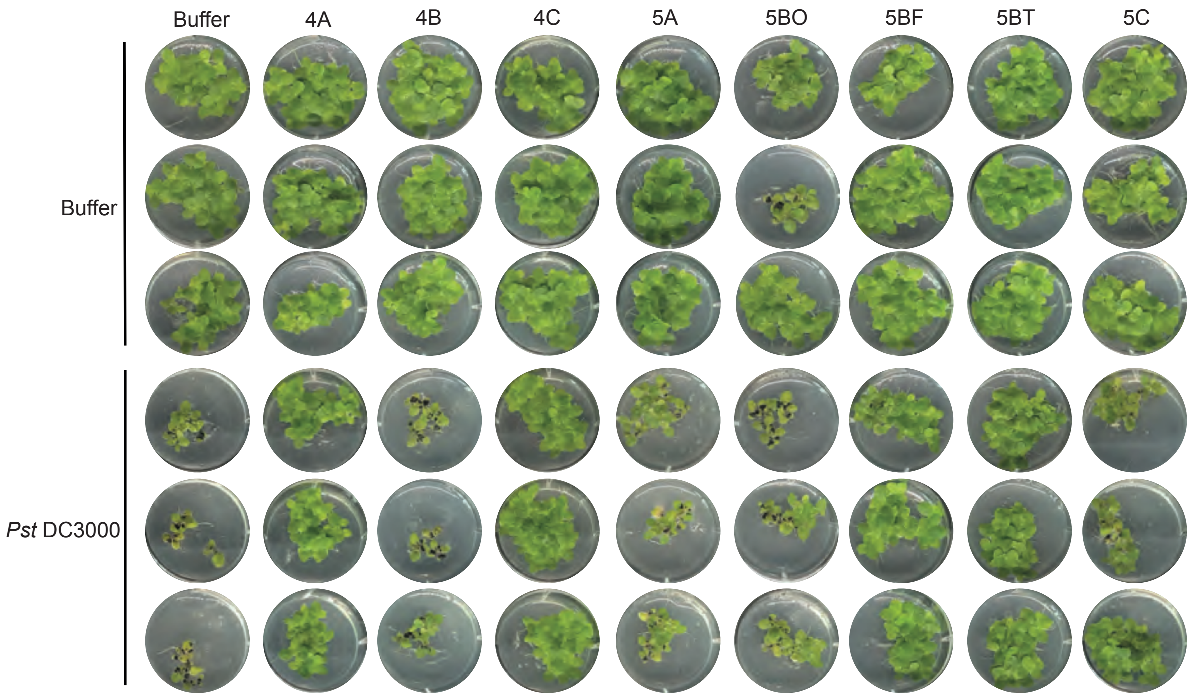


**Figure S7. Experiment 2 *Spirodela polyrhiza* co-inoculated with bacteria isolated from the 0.2 µm water filtrate and *Pst* DC3000.**
Images 3 weeks after co-inoculation of *S. polyrhiza* with *Pst* DC3000 and bacteria isolated from 0.2 µm water filtrate. Each well is an individual replicate.

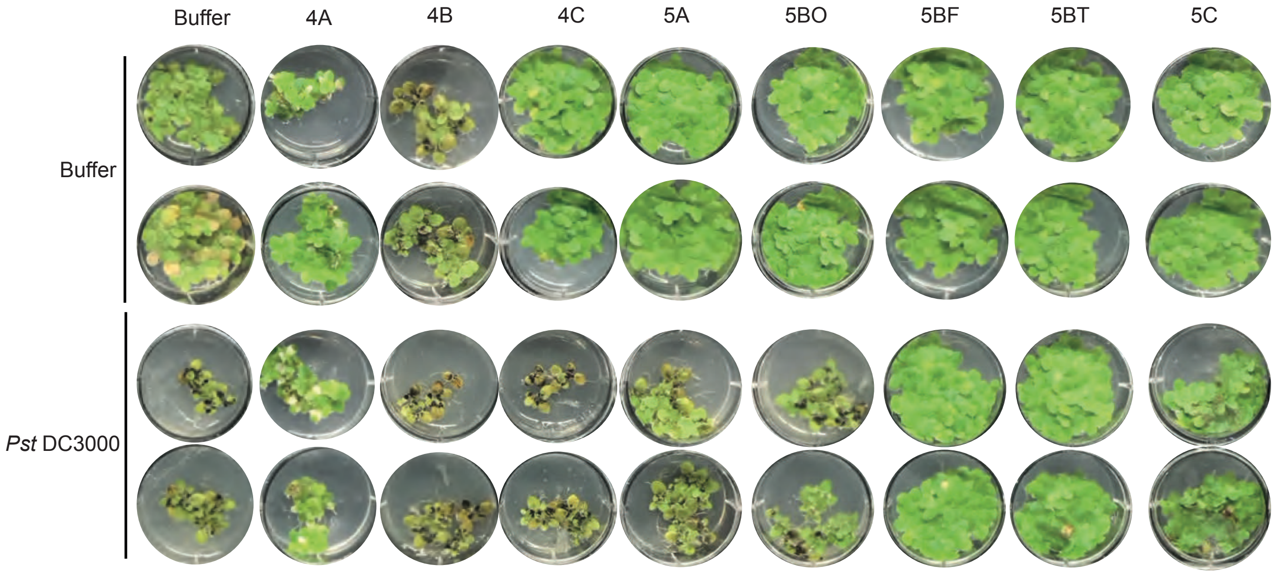


**Figure S8 Experiment 3 *Spirodela polyrhiza* co-inoculated with bacteria isolated from the 0.2 µm water filtrate and *Pst* DC3000.**
Images 3 weeks after co-inoculation of *S. polyrhiza* with *Pst* DC3000 and bacteria isolated from 0.2 µm water filtrate. Each well is an individual replicate.

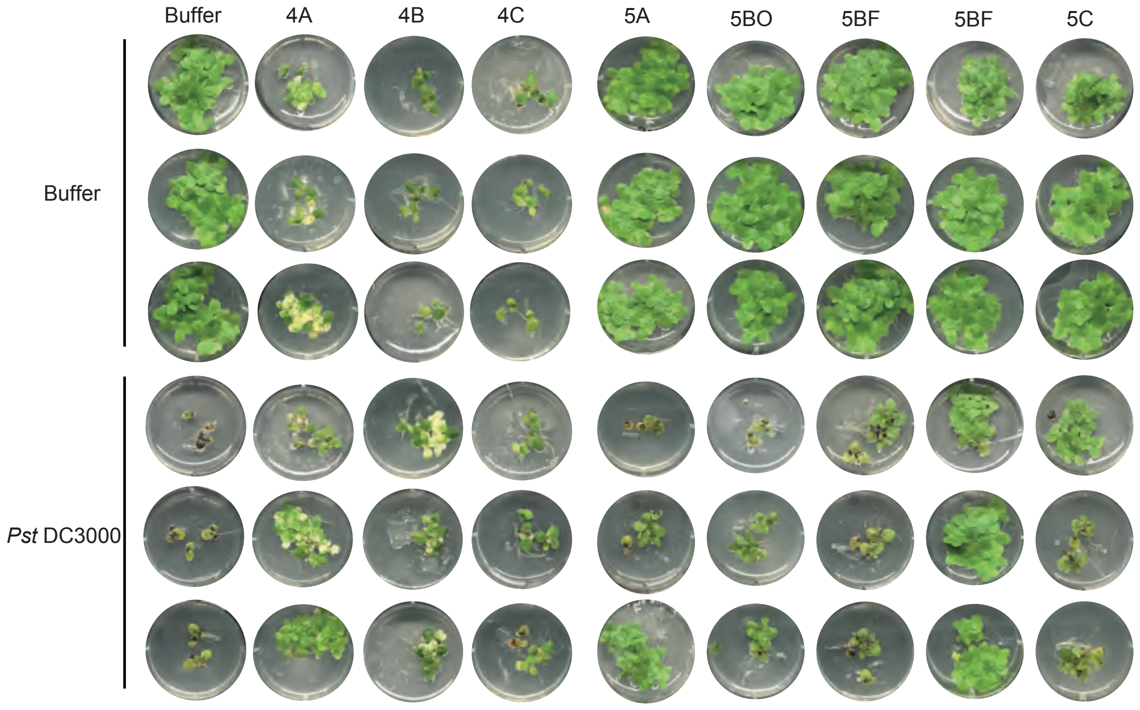


**Figure S9. Experiment 4 *Spirodela polyrhiza* co-inoculated with bacteria isolated from the 0.2 µm water filtrate and *Pst* DC3000.**
Images 3 weeks after co-inoculation of *S. polyrhiza* with *Pst* DC3000 and bacteria isolated from 0.2 µm water filtrate. Each well is an individual replicate.

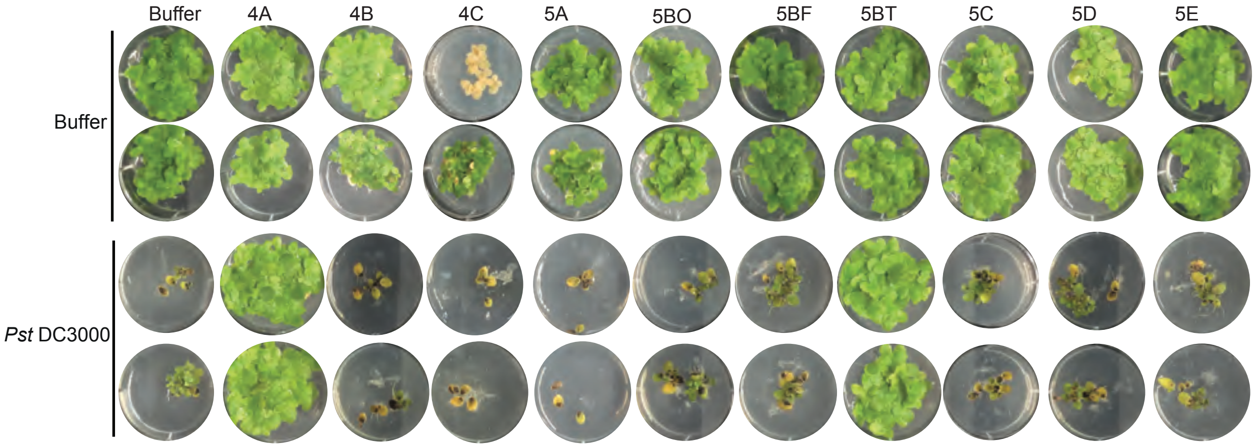


**Figure S10. Experiment 5 *Spirodela polyrhiza* co-inoculated with bacteria isolated from the 0.2 μm water filtrate and *Pst* DC3000.**
Images 3 weeks after co-inoculation of *S. polyrhiza* with *Pst* DC3000 and bacteria isolated from 0.2 μm water filtrate. Each well is an individual replicate.

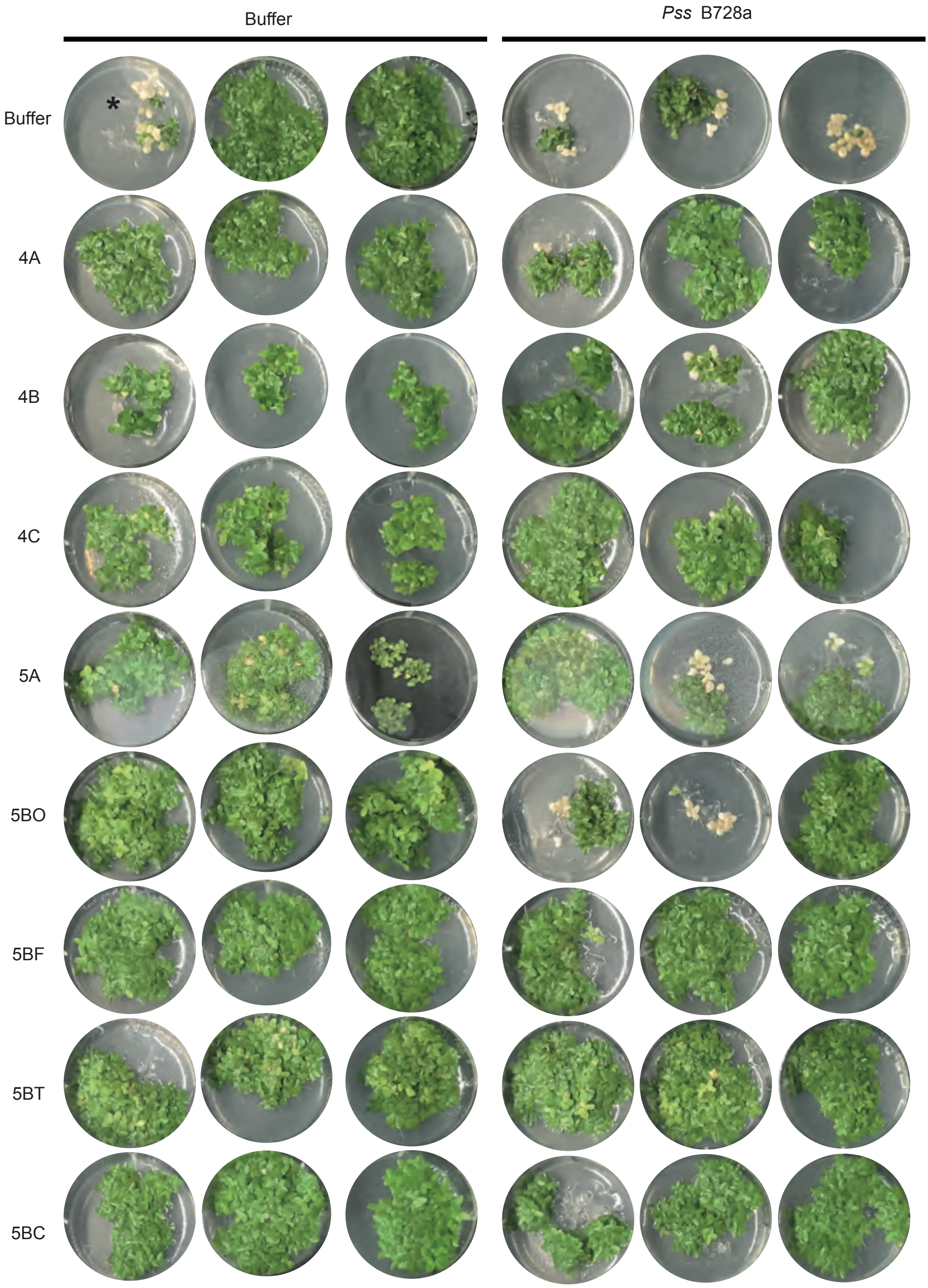


**Figure S11. Experiment 1 *Landoltia punctata* 5635 inoculated with pathogens and environmental bacteria.**
Images 3 weeks post-inoculation of *L. punctata* 5635 inoculated with a combination of bacterial strains. Individual wells are separate biological replicates. Asterix indicates cross-contamination from a neighboring well.

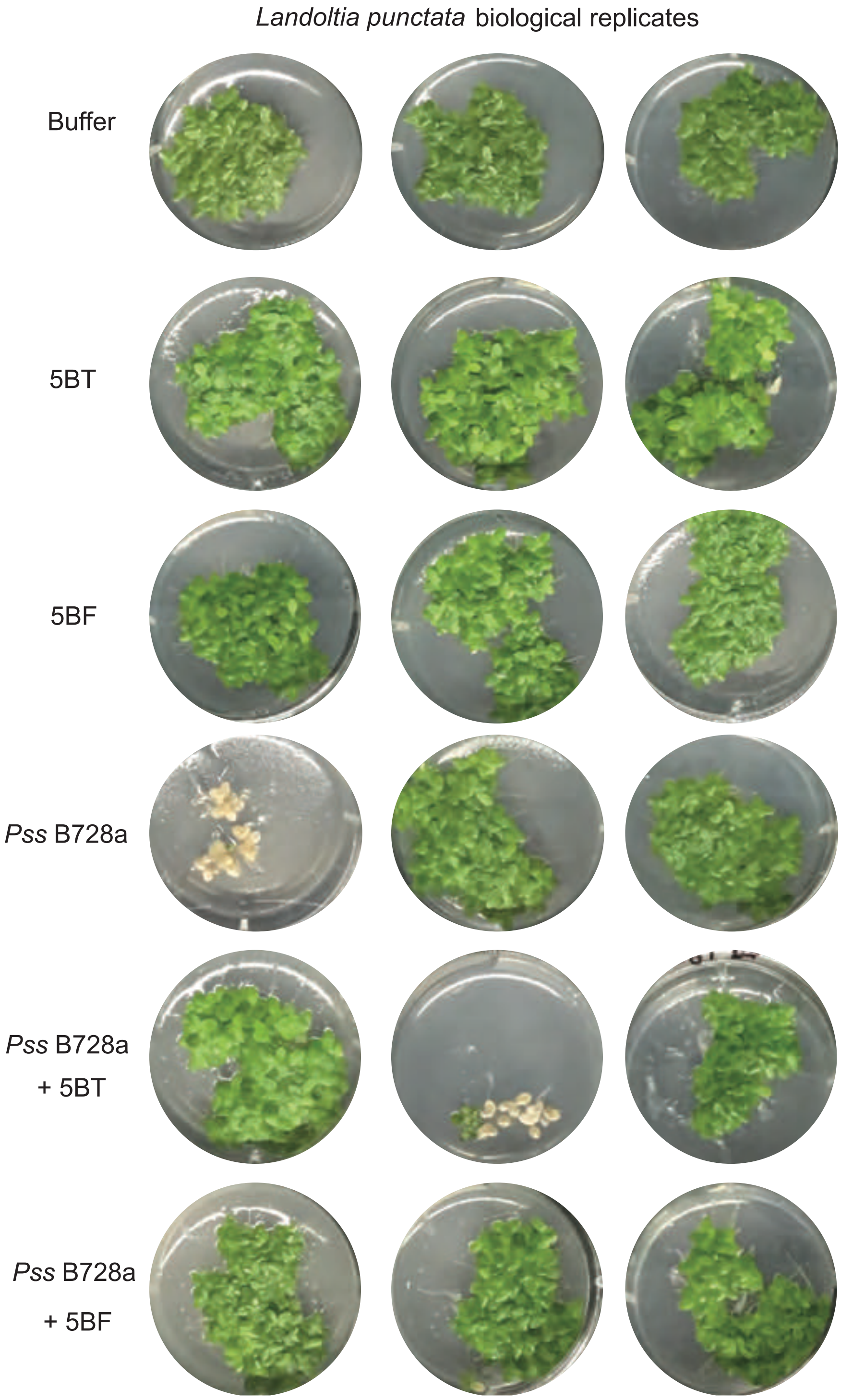


**Figure S12. Experiment 2 *Landoltia punctata* 5635 inoculated with pathogens and environmental bacteria.**
Images 3 weeks post-inoculation of *L. punctata* 5635 inoculated with combination of bacterial strains. Individual wells are separate biological replicates.

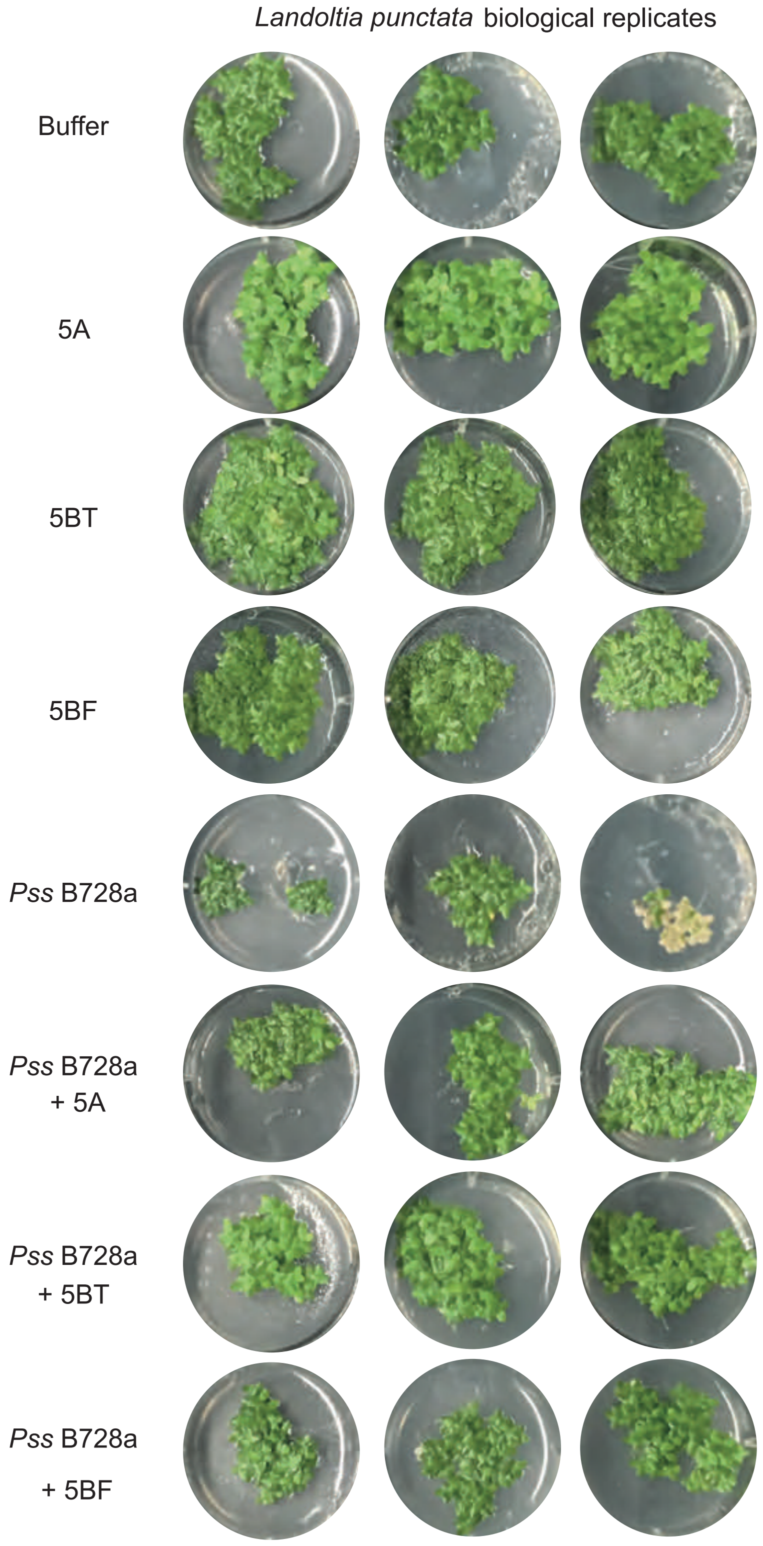


**Figure S13. Experiment 3 *Landoltia punctata* 5635 inoculated with pathogens and environmental bacteria.**
Images 3 weeks post-inoculation of *L. punctata* 5635 inoculated with combination of bacterial strains. Individual wells are separate biological replicates.

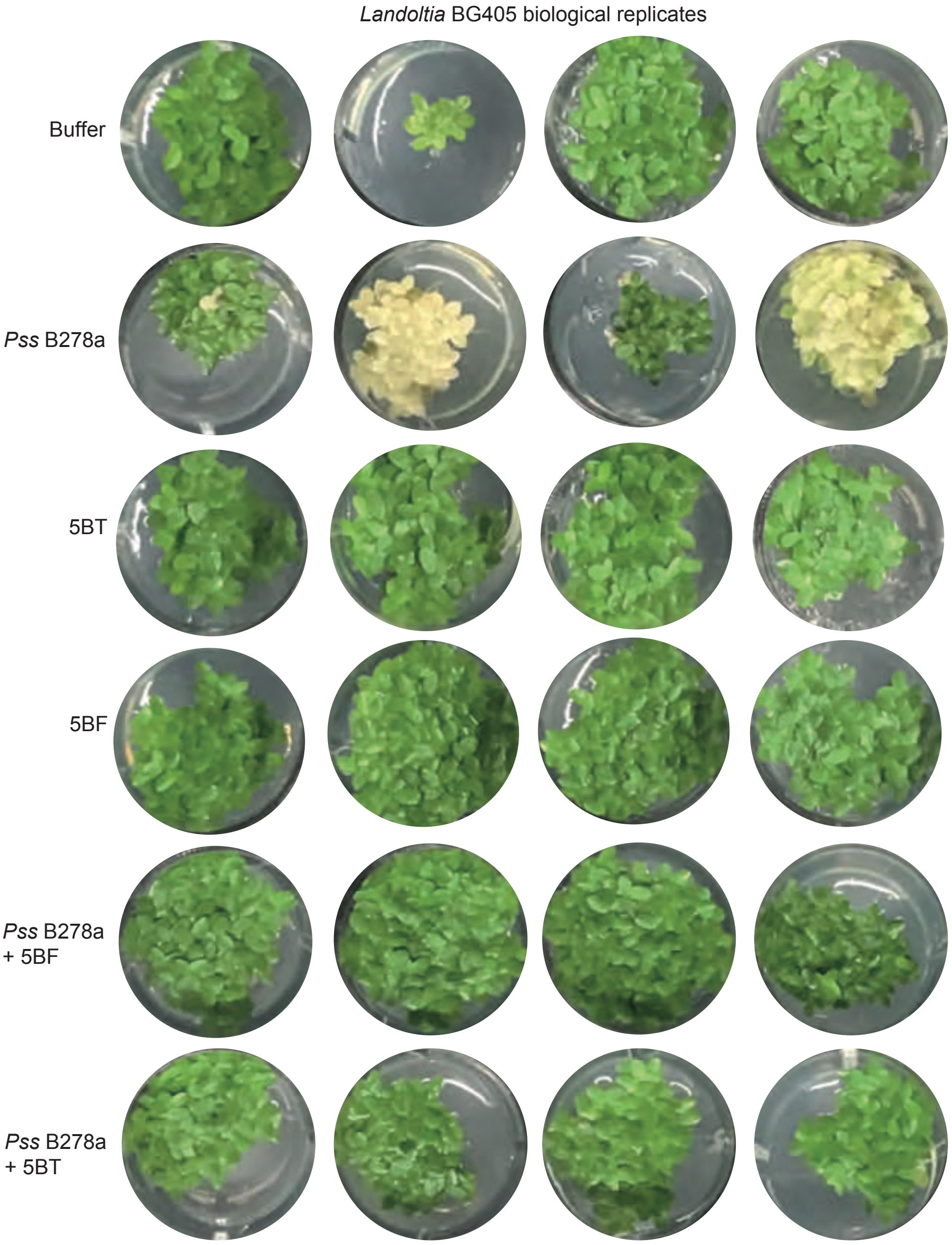


**Figure S14. Experiment 1 Isolate 5BF's protection of *L. punctata* BG405 from symptomatic *Pss* B728a infection.**
Images 3 weeks after co-inoculation of *L. punctata* with *Pss* B728a and bacteria isolated from 0.2 µm water filtrate. Four replicates are shown per treatment (full experiment has been independently replicated 2 times, see Sfig 9). Asterix indicates potential well contamination with *Pss* B728a.

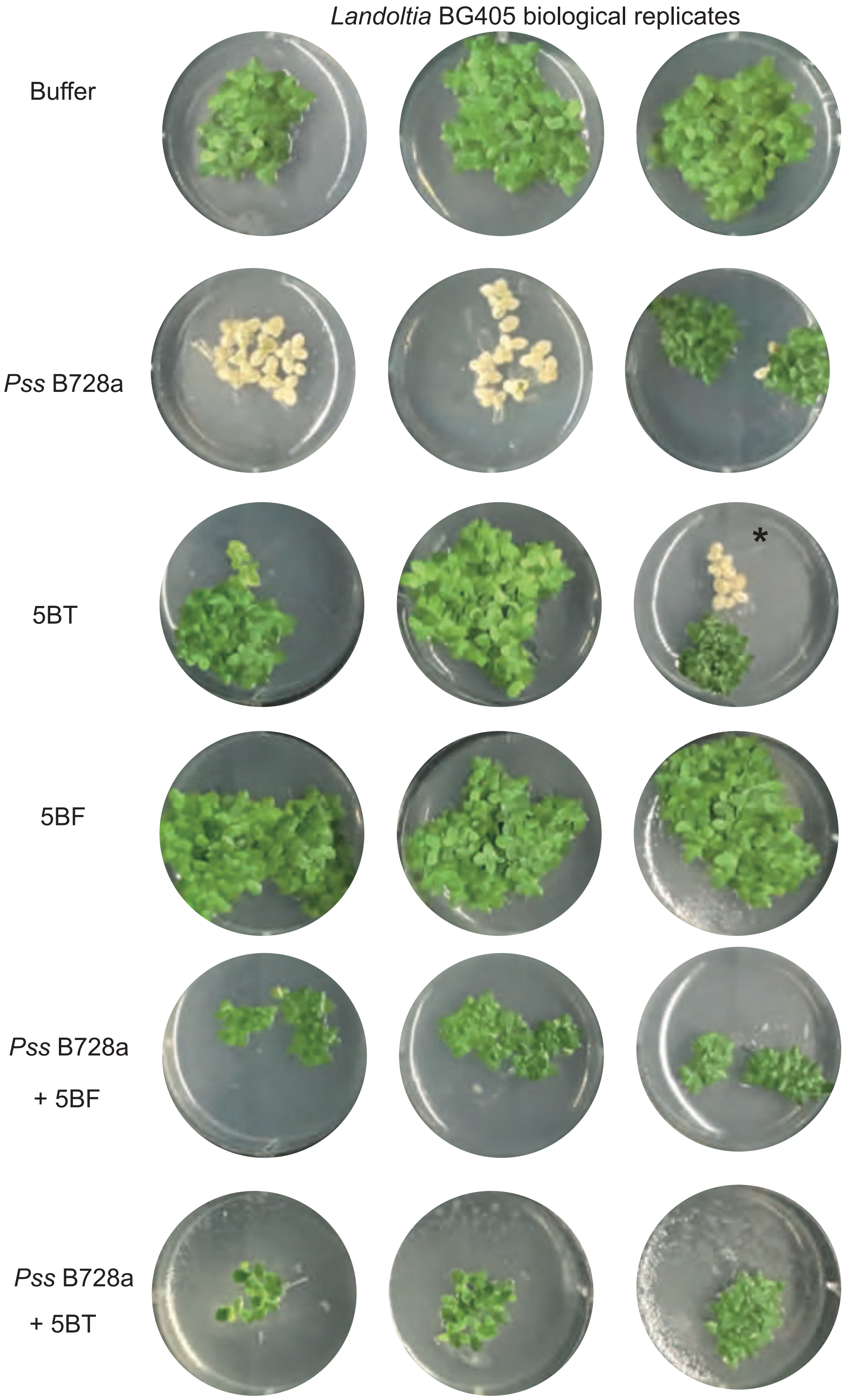


**Figure S15. Experiment 2 *Landoltia* BG405 inoculated with pathogens and environmental bacteria.**
Images 3 weeks post-inoculation of *Landoltia* BG405 inoculated with a combination of bacterial strains. Individual wells are separate biological replicates.

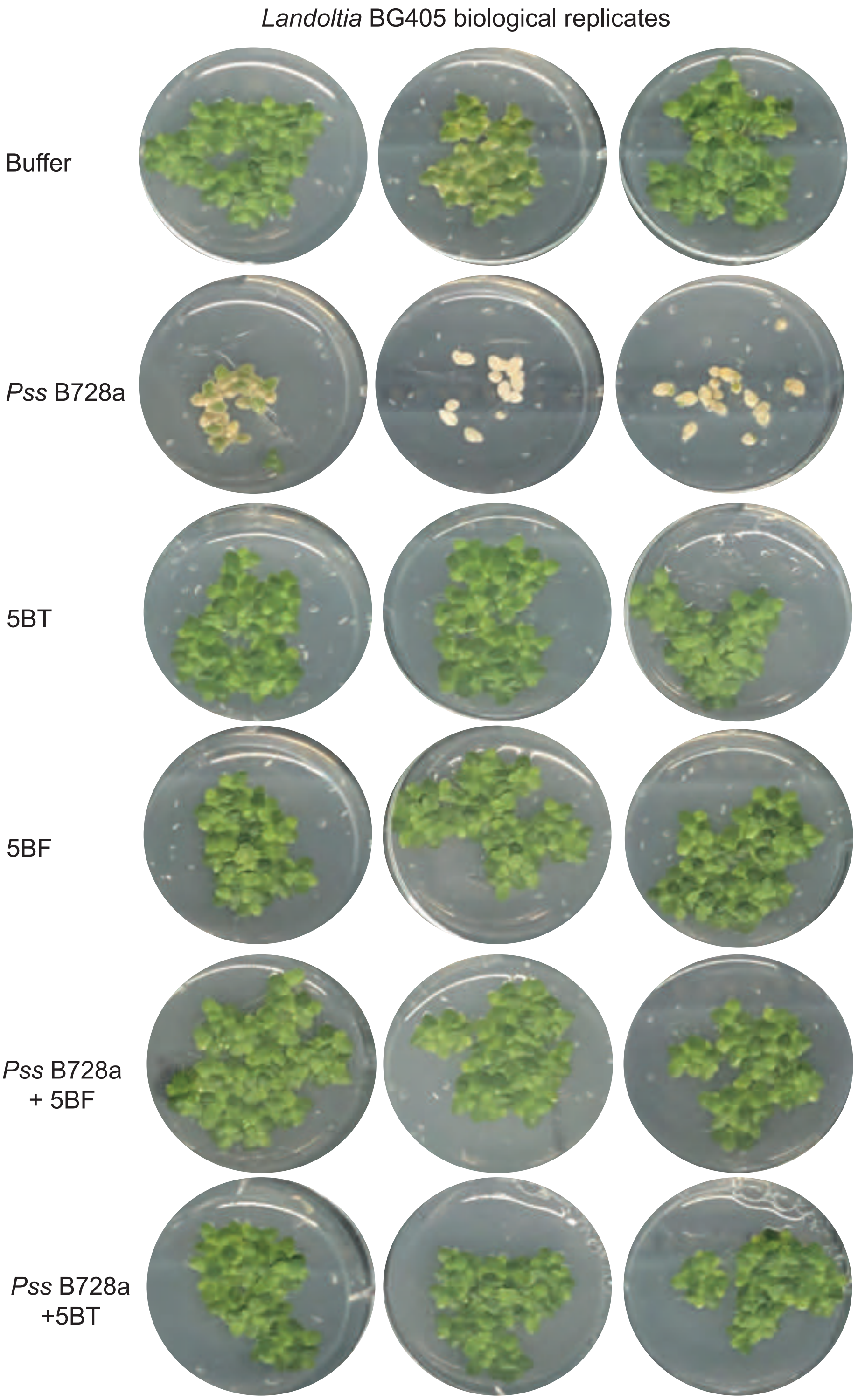


**Figure S16. Experiment 2 Landoltia BG405 with pathogens and environmental bacteria.** Images 3 weeks post-inoculation of Landoltia BG405 inoculated with a combination of bacterial strains. Individual wells are separate biological replicates.

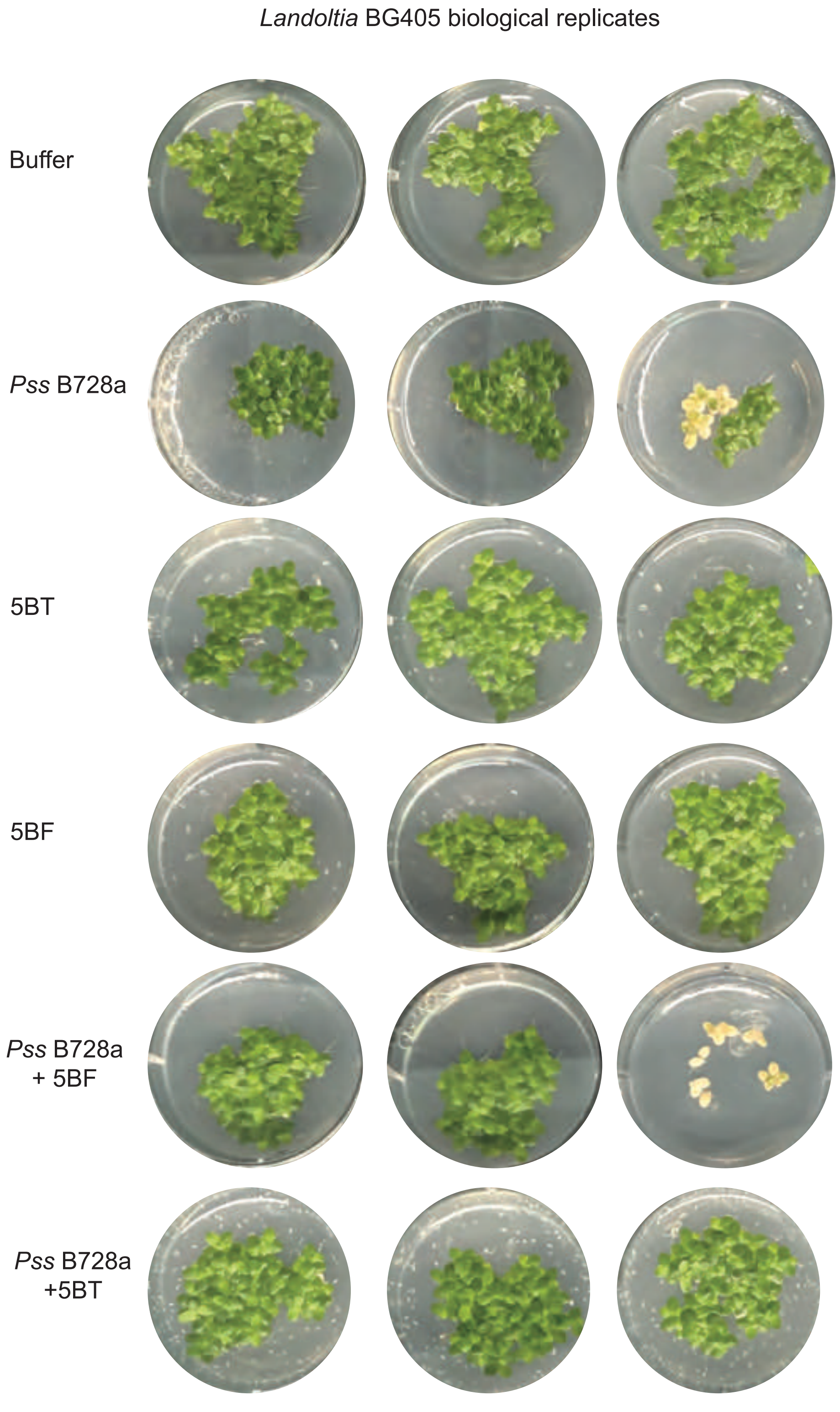


**Figure S17. Experiment 3 Landoltia BG405 with pathogens and environmental bacteria.** Images 3 weeks post-inoculation of Landoltia BG405 inoculated with a combination of bacterial strains. Individual wells are separate biological replicates.

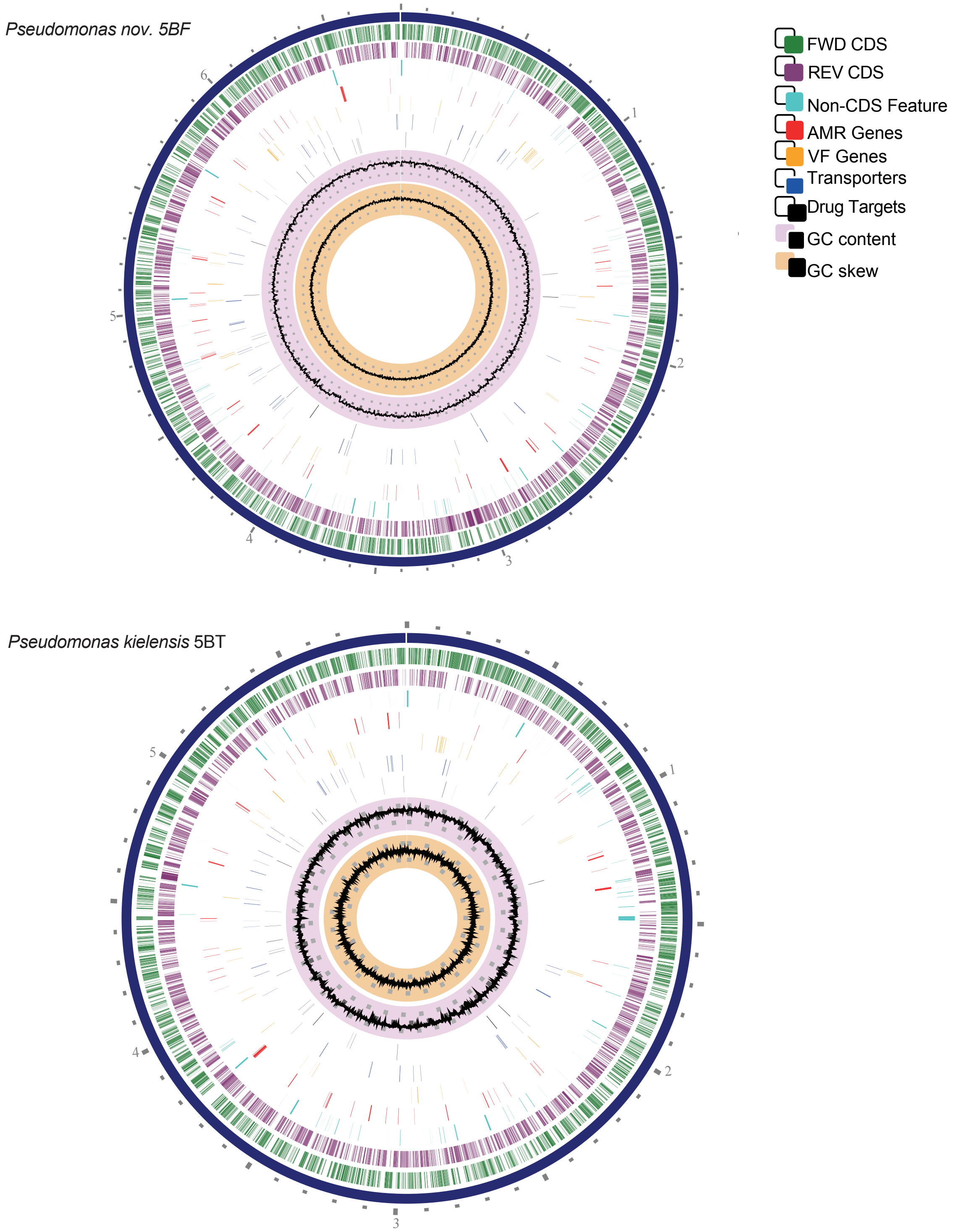


**Figure S18. Circos plot of Pseudomonas nov. 5BF and Pseudomonas kielensis 5BT genome assemblies.**

For genome plots in A and B the from innermost to outermost the rings indicate: GC skew, GC content, putative drug targets, transporters, virulence factor genes, antimicrobial resistance genes, non-coding sequence feature, reverse coding sequence, forward coding sequence and chromosomal location.

A.

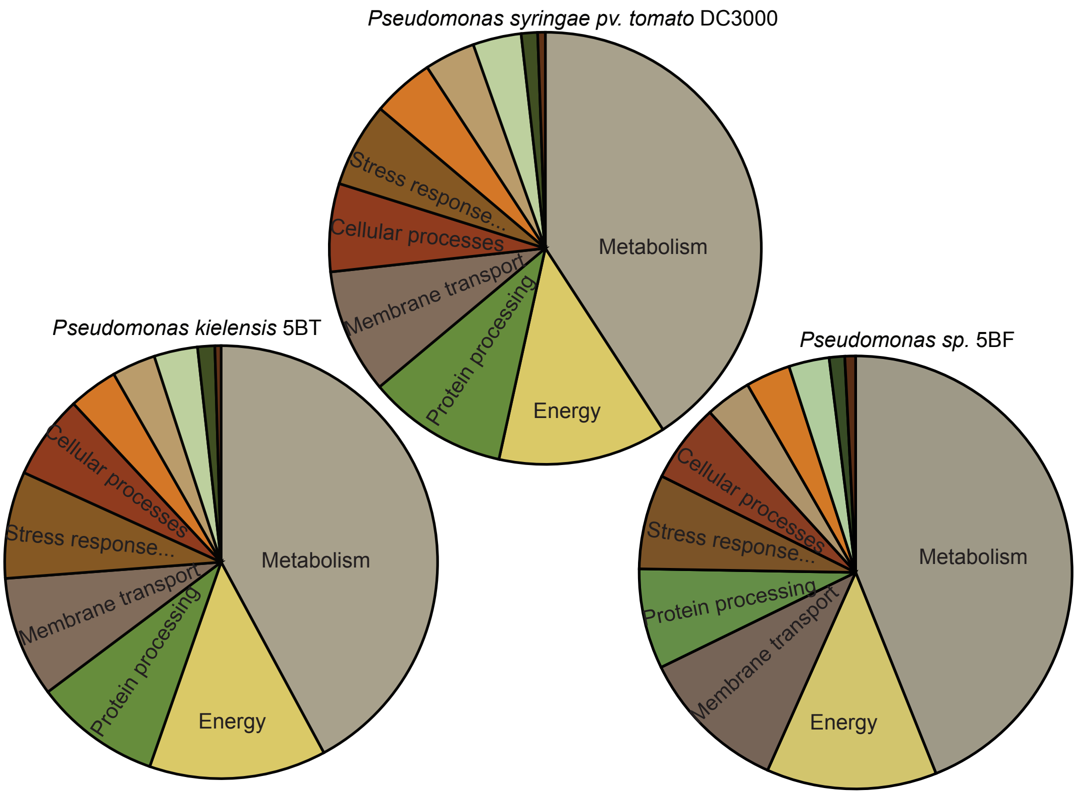


B.

| No. subsystems | DC3000 | 5BT | 5BF |
|---|---|---|---|
| Metabolism | 112 | 112 | 123 |
| Energy | 31 | 30 | 29 |
| Membrane transport | 30 | 29 | 31 |
| Protein processing | 43 | 43 | 41 |
| Stress response, defense, virulence | 30 | 34 | 33 |
| Cellular processes | 17 | 16 | 18 |
| Cell envelope | 11 | 8 | 9 |
| DNA processing | 20 | 18 | 19 |
| RNA processing | 14 | 15 | 15 |
| Miscellaneous | 4 | 7 | 6 |
| Regulation and cell signaling | 4 | 3 | 5 |

C.

| No. genes | DC3000 | 5BT | 5BF |
|---|---|---|---|
| Metabolism | 936 | 1087 | 1368 |
| Energy | 289 | 341 | 396 |
| Membrane transport | 214 | 233 | 346 |
| Protein processing | 240 | 243 | 233 |
| Stress response, defense, virulence | 145 | 206 | 220 |
| Cellular processes | 151 | 162 | 184 |
| Cell envelope | 87 | 84 | 107 |
| DNA processing | 106 | 95 | 104 |
| RNA processing | 83 | 84 | 94 |
| Miscellaneous | 28 | 33 | 36 |
| Regulation and cell signaling | 13 | 12 | 25 |

**Figure S19. Pie-chart of number of subsystems within superclasses for *Pseudomonas* species of interest.**
A. Patric genome annotation of the proportion of subsystems assigned to the major superclasses. Colors of pie sectors correspond to table B along with the absolute numbers of subsystems. C. Provides the numbers of genes that are assigned to each superclass for the three species.

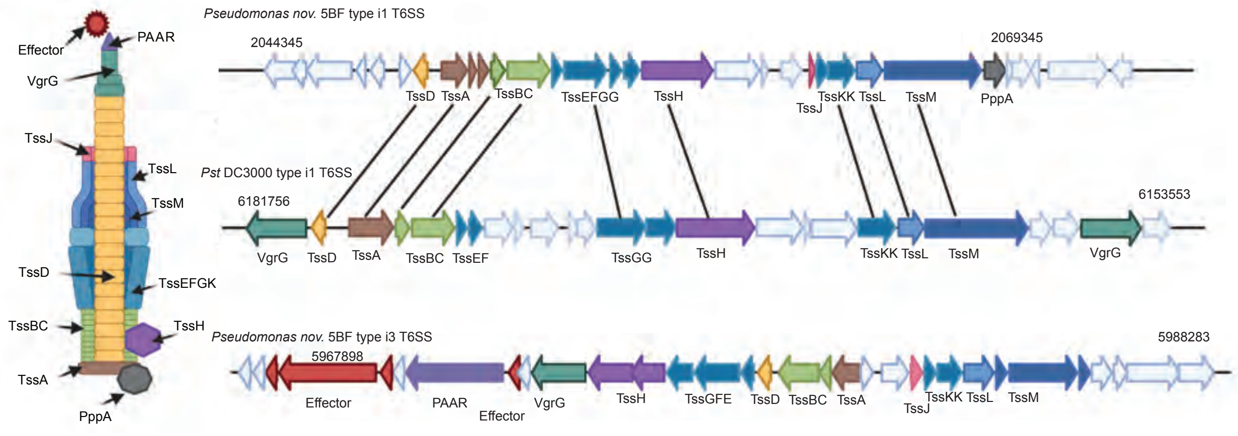


**Figure S20. Comparison of isolate *Pseudomonas nov.* 5BF type i1 T6SS with *Pst* DC3000 type i1 T6SS.**

A. SecReT6 comparison of synteny and gene identity between *Pseudomonas nov.* 5BF Type i1 T6SS (2044345..2069345) (top) and *Pst* DC3000 type i1 T6SS (6149242..6189243) (bottom). Colored arrows indicate T6SS component genes which correspond to the T6SS diagram. White arrows are genes which are not part of the T6SS apparatus. Gene identifiers are given above or below the genes. B. Depiction of *Pseudomonas nov.* 5BF T6SS i3 locus (5967898..5988283) with arrows as in A.

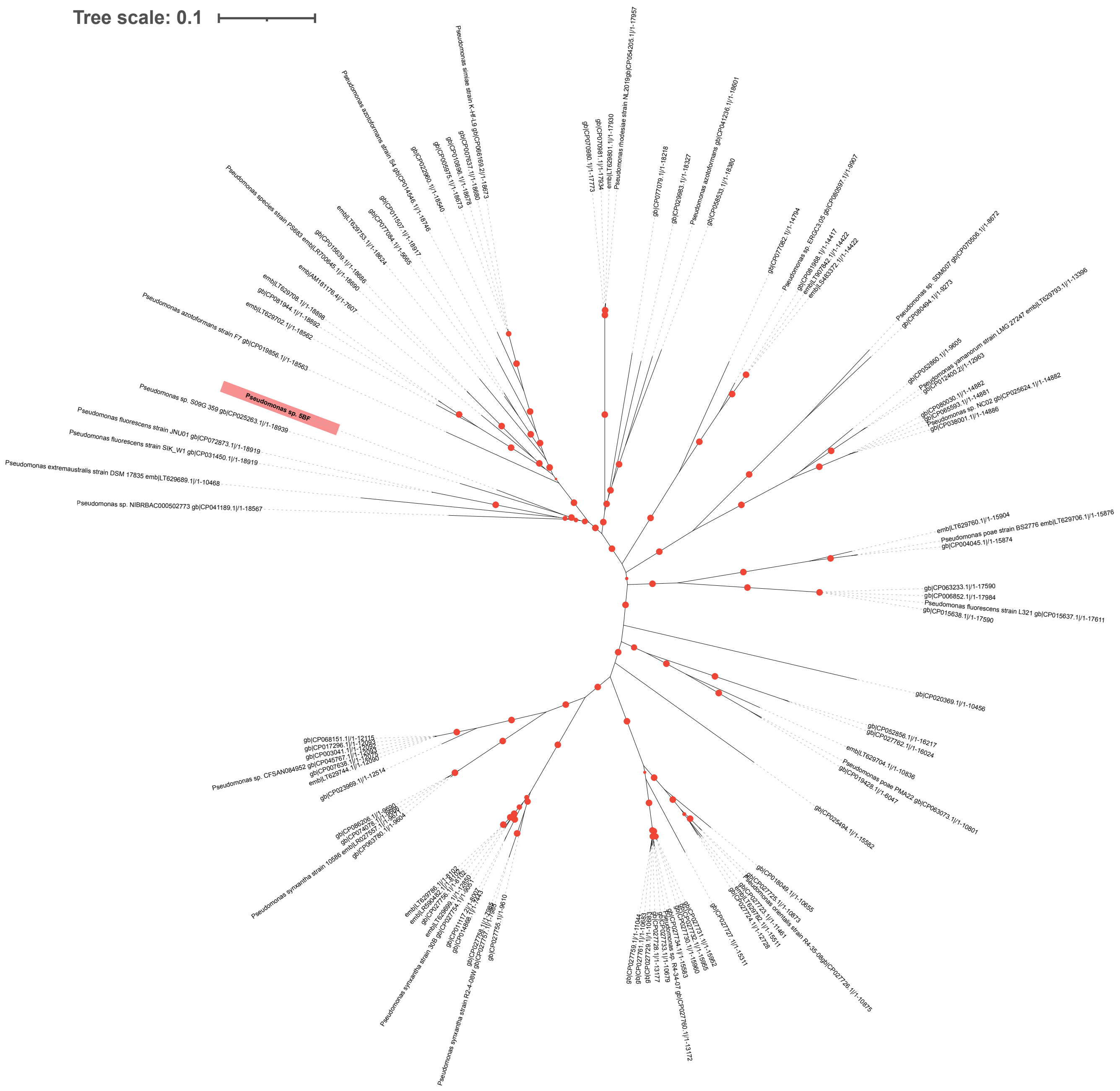


**Figure S21. NCBI phylogeny of the top 100 BLASTN hits queried with the *Pseudomonas nov.* 5BF type i1 T6SS locus.**

Maximum likelihood phylogeny of the type VI secretion system locus predicted by Secret6 between 2049345..2068484 of Pseudomonas nov. 5BF genome. and top 100 blastn hits. Red dots indicate bootstrap values >80.

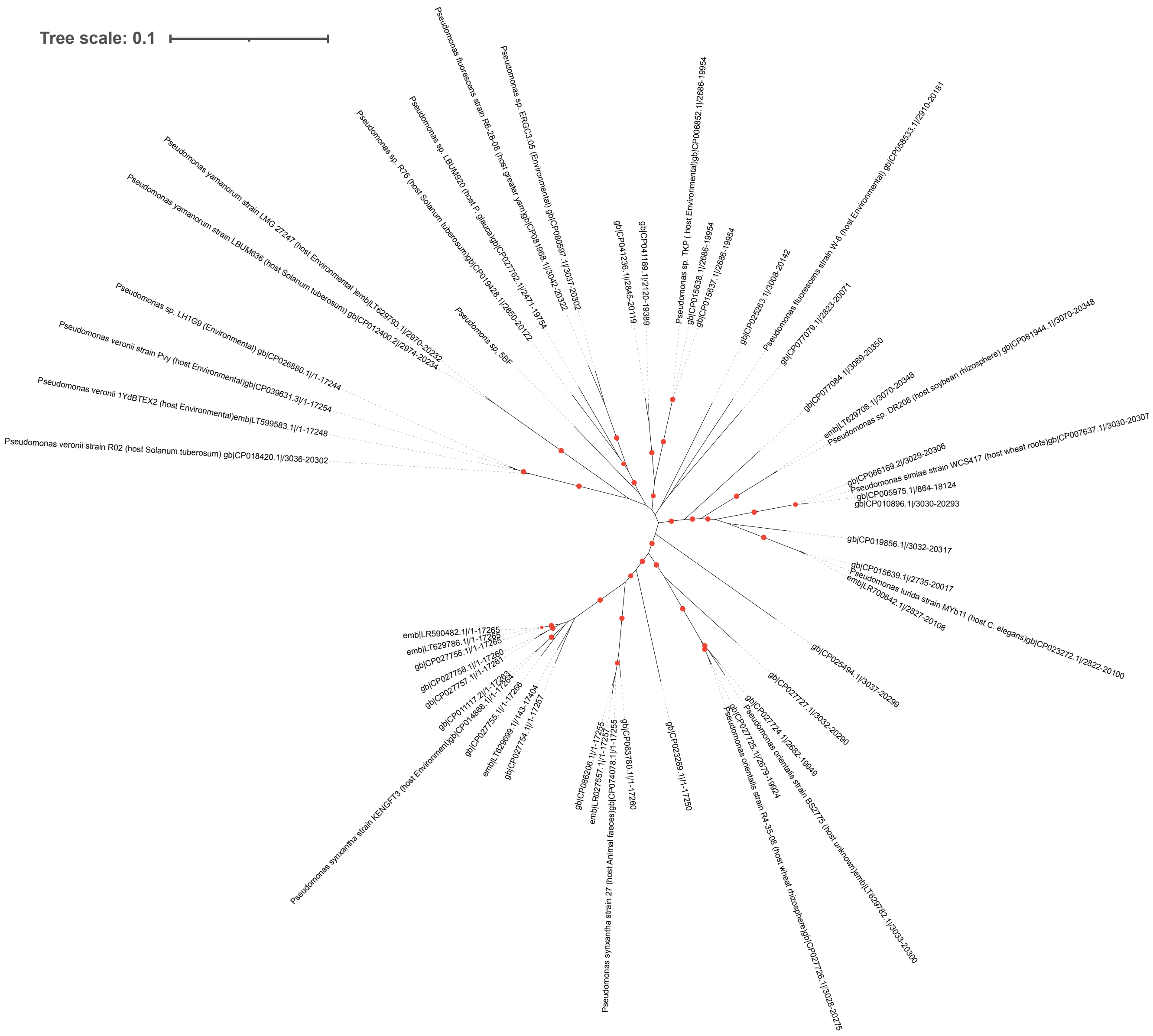


**Figure S22 NCBI phylogeny of the top 50 BLASTN hits to the *Pseudomonas nov.* 5BF type i3 T6SS locus**

Maximum likelihood phylogeny of the type VI secretion system locus with chromosomal location **5967898..5988283** predicted by Secret6 and top 50 blastn hits. Red dots indicate bootstrap values >80.

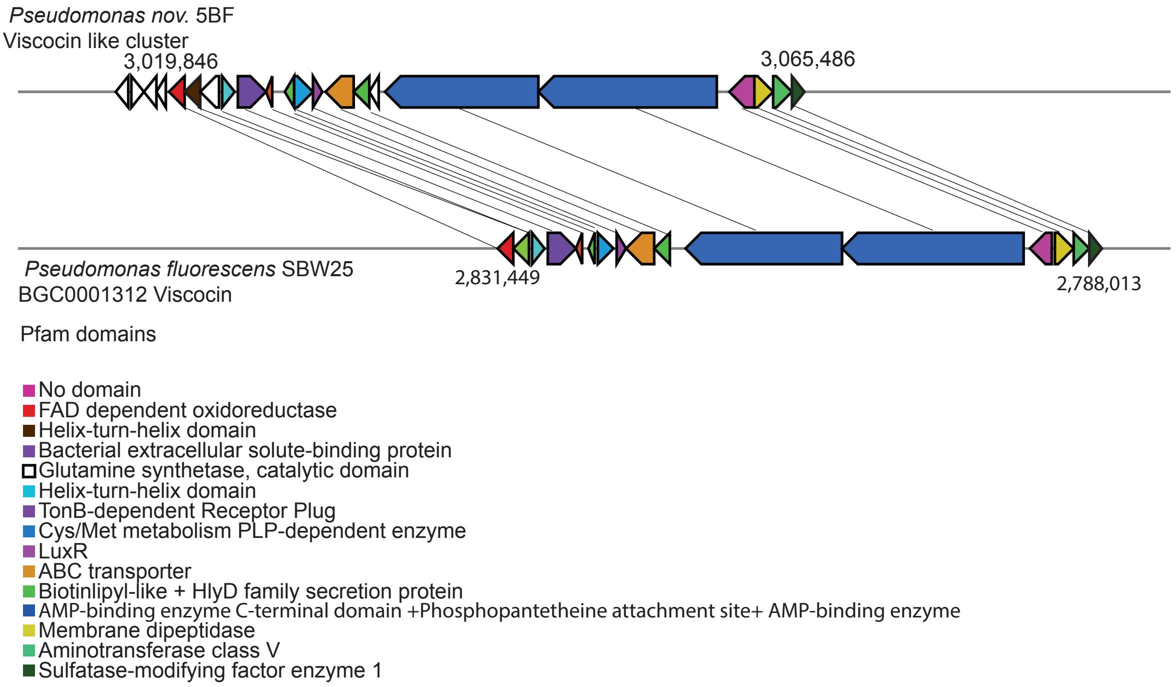


**Figure S23. Synteny between *Pseudomonas nov.* 5BF and *Pseudomonas fluorescens* SBW25 at the viscosin secondary metabolic islands**

Schematic diagram of gene synteny between P. nov 5BF and P. fluorescens SBW25. Genes are depicted as arrows, lines are drawn between best blast hit of genes from antismash5.0 known cluster blast. MIBiG repository accession number for gene clusters is BGC0001312 number. Colors of arrows indicate pfam domains of genes.

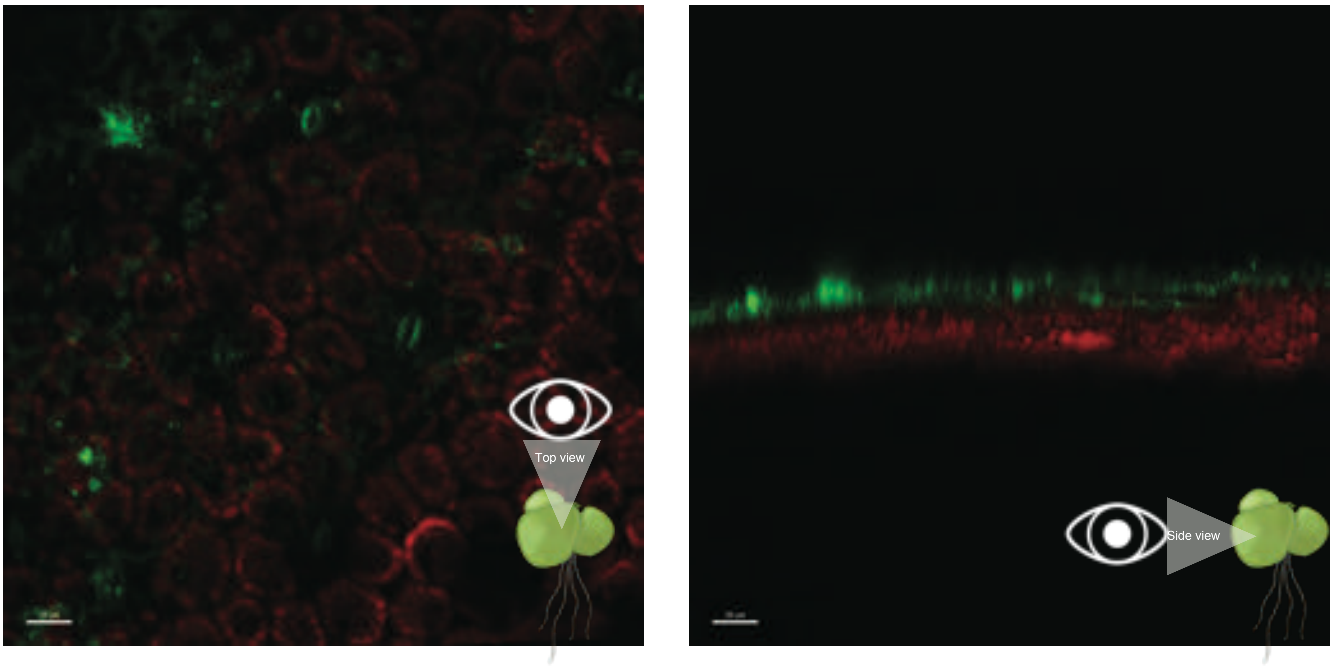


**Figure S24. Confocal microscopy of Z-stack of 5BT on *Spirodela polyrhiza* fronds..**
Green false coloring indicates *P. kielensis* 5BT and red shows the chloroplast fluorescence. A. is a top down view of the leaf surface and B is a view across the z-axis. Images were taken 7 days post inoculation with *P. kielensis* 5BT.
.

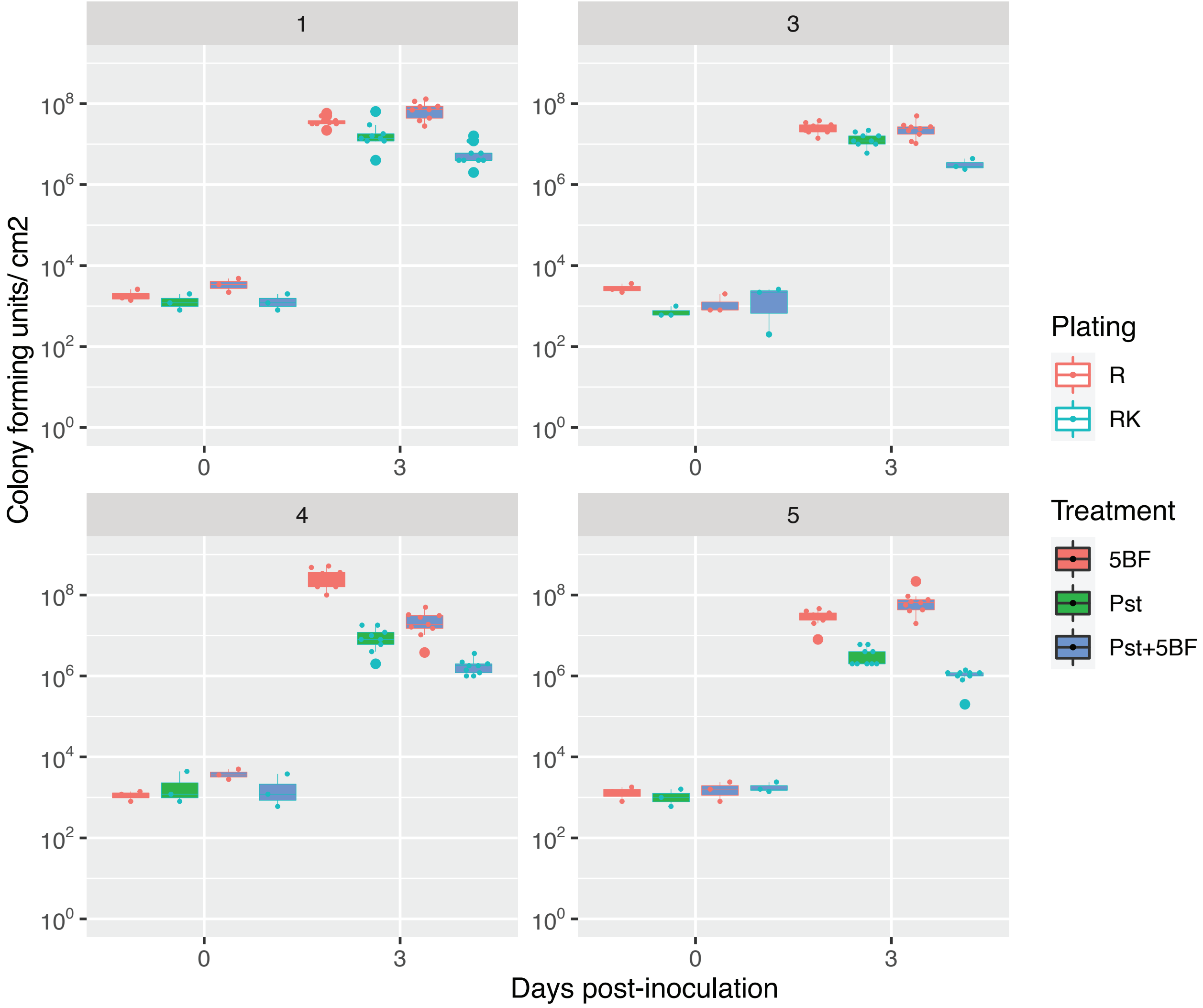


**Figure S25. Growth curve split by experiment *Spirodela polyrhiza* co-inoculated with low bacterial load of *Pseudomonas* nov. 5BF and or *Pst* DC3000.**
Images 10 days post-inoculation of *S. polyrhiza* with *Pst* DC3000 and/ or *Pseudomonas* nov. 5BF. Each well is an individual replicate and was sampled when treated for bacteria for the corresponding experiment growth curve.

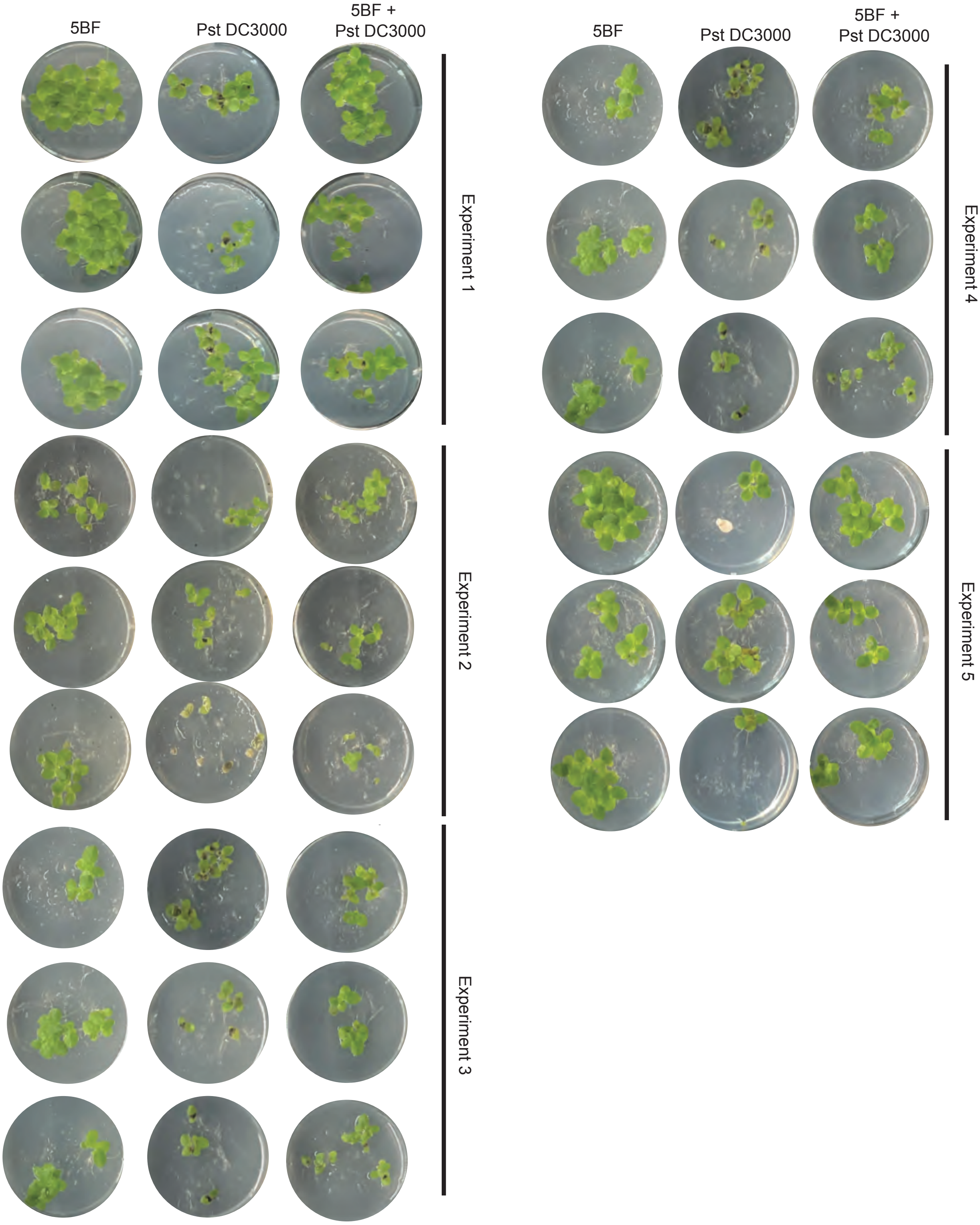


**Figure S26. Experiments 1-5 *Spirodela polyrhiza* co-inoculated with low bacterial load of *Pseudomonas* nov. 5BF and or *Pst* DC3000.**
Images 10 days post-inoculation of *S. polyrhiza* with *Pst* DC3000 and/ or *Pseudomonas* nov. 5BF. Each well is an individual replicate and was sampled when treated for bacteria for the corresponding experiment growth curve.

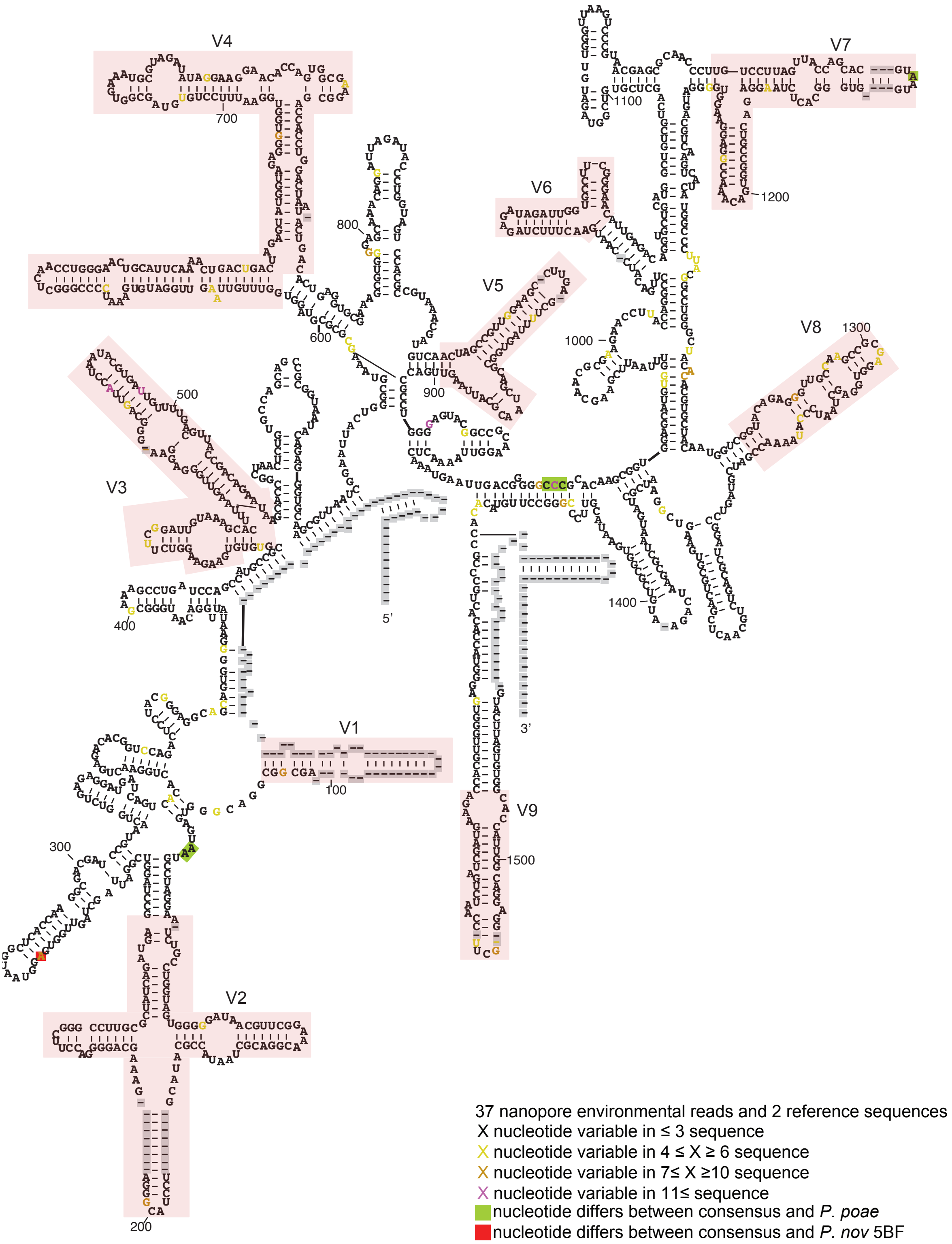


**Figure S27. Consensus sequence of nanopore reads and reference 16S rRNA sequence from secondary structure informed sequence alignment.**
Nucleotide sequence displayed is consensus sequence among the 37 nanopore reads that passed SNP error filter for similarity to *P. nov* 5BF and/ or *P. poae* and their reference sequence. Background colors indicate nucleotides that differ from consensus in reference strains. Colors of nucleotide text relate to the frequency of SNP at that position. Grey squares with dash indicate absence of sequence.

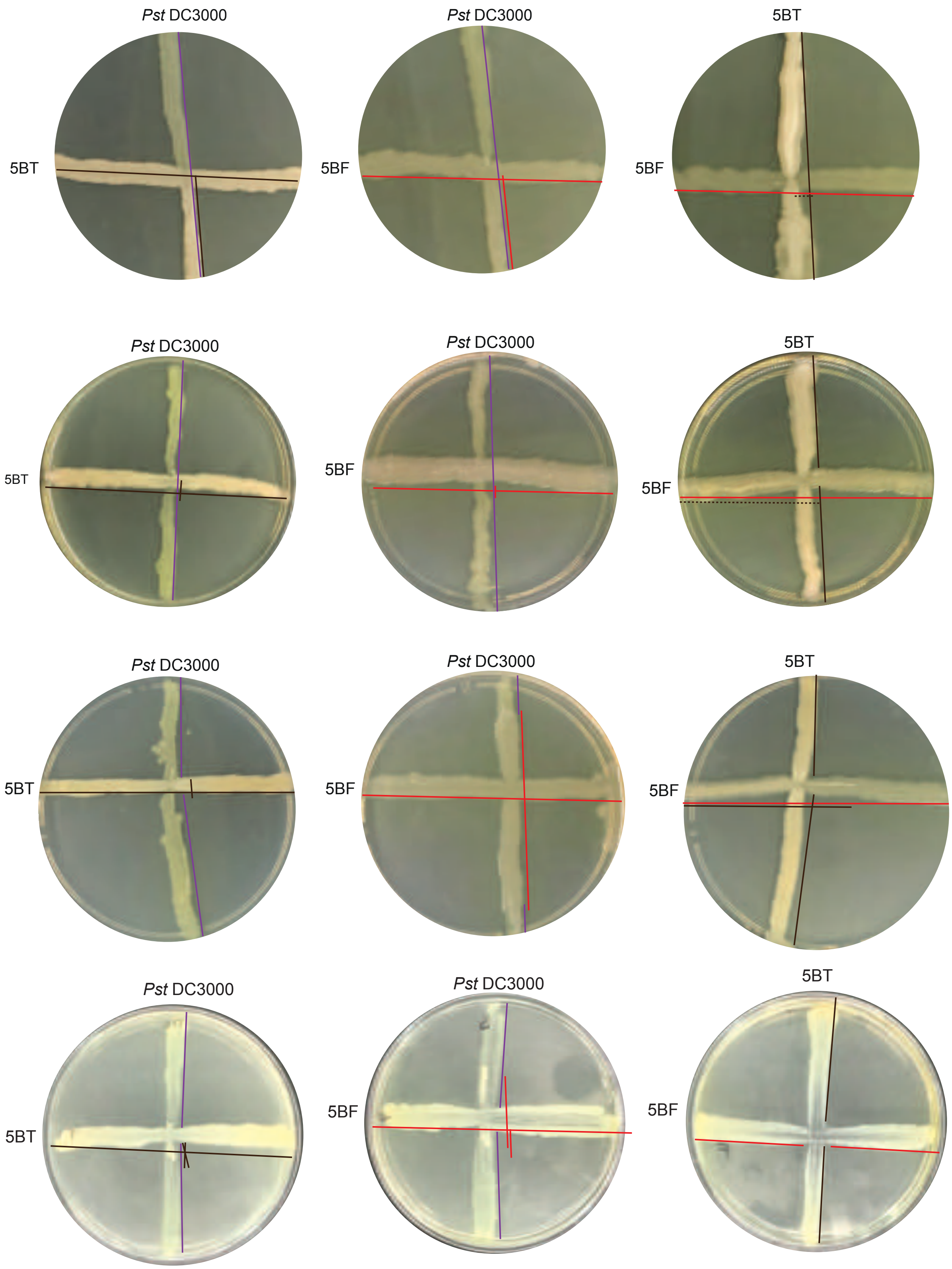


**Figure S28. *Pseudomonas* interaction streak replicates.**
Experiment 2-4 of cross-streaks of different *Pseudomonas* sp. on LB agar plates 3 days post plating. Colored lines shifted to side of streaks highlight colony growth pattern brown`- *Pst* DC3000, red *Pseudomonas nov.* 5BF and blue *Pseudomonas kielensis* 5BT.

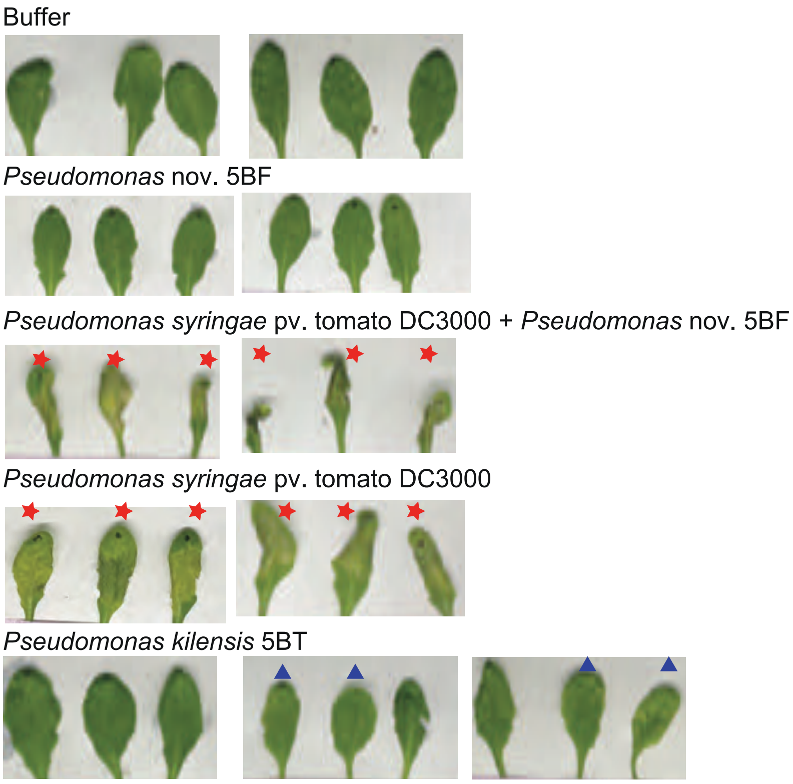


**Figure S29. High OD syringe infiltration of *A. thaliana* leaves with *Pseudomonas spp*. Experiment 1**
Experiment 1 additional biological replicates of high OD *Pseudomonas spp*. inoculations. Photographs of *A. thaliana* leaves 48 hrs post inoculation with various *Pseudomonas spp.* .

Buffer

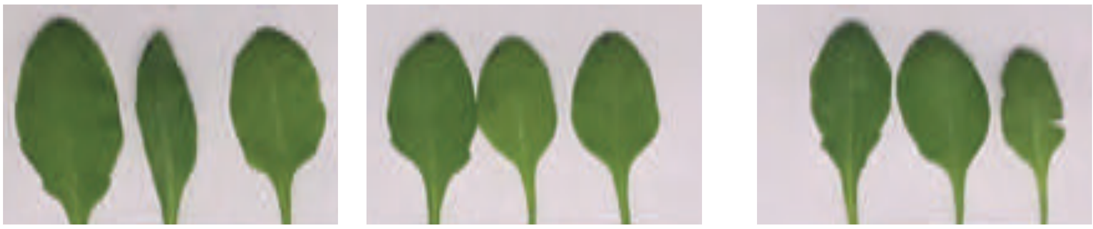

*Pseudomonas syringae* pv. tomato DC3000

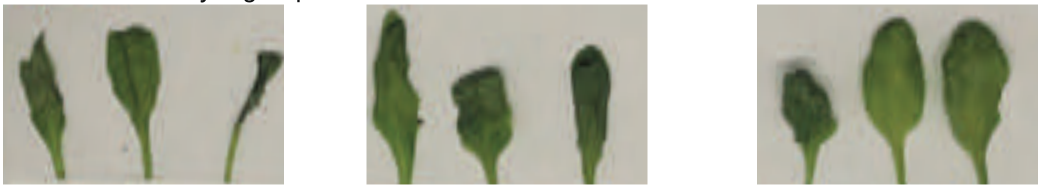

*Pseudomonas* nov 5BF

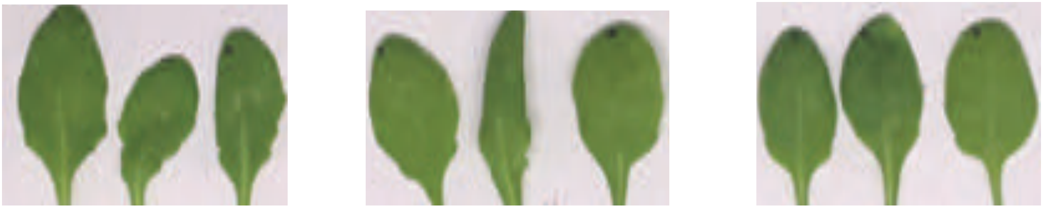

*Pseudomonas kielensis* 5BT

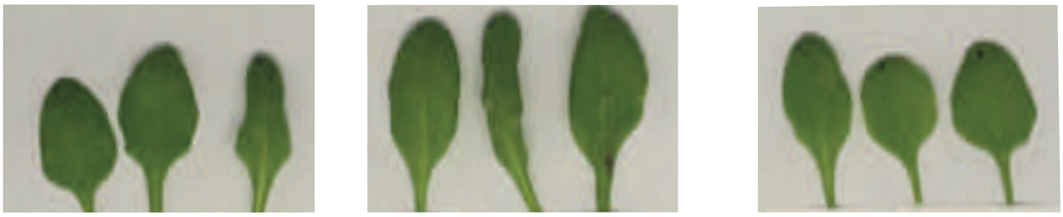

*Pseudomonas syringae* pv. tomato DC3000
+ *Pseudomonas* nov 5BF

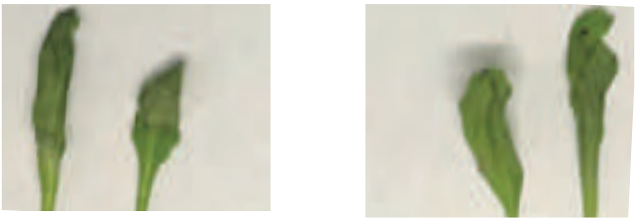

**Figure S30. High OD syringe infiltration of *A. thaliana* leaves with *Pseudomonas spp.***
Experiment 2 of high OD *Pseudomonas spp.* inoculations. Photographs of *A. thaliana* leaves 48hrs post inoculation with various *Pseudomonas spp.* .

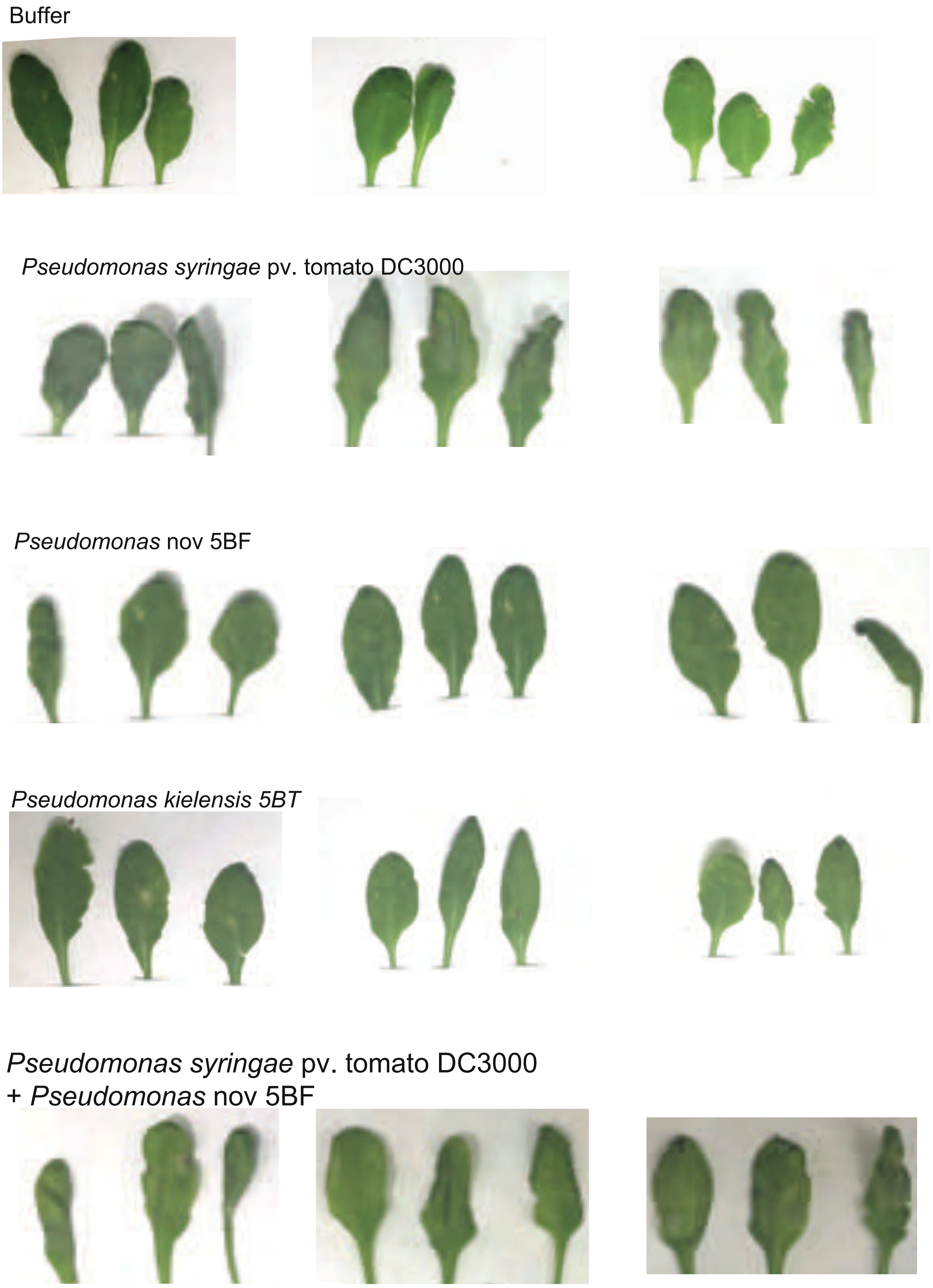


**Figure S31. High OD syringe infiltration of *A. thaliana* leaves with *Pseudomonas spp.***
Experiment 3 of high OD *Pseudomonas spp.* inoculations. Photographs of A. thaliana leaves 48hrs post inoculation with various *Pseudomonas spp*. .

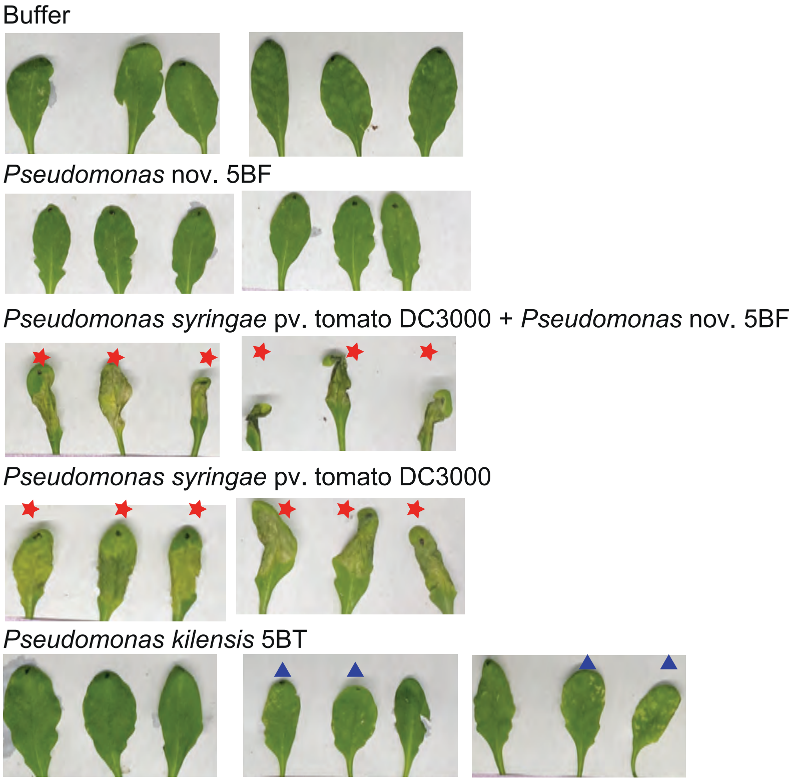


**Figure S29. High OD syringe infiltration of *A. thaliana* leaves with *Pseudomonas spp*. Experiment 1**
Experiment 1 additional biological replicates of high OD *Pseudomonas spp*. inoculations. Photographs of *A. thaliana* leaves 48 hrs post inoculation with various *Pseudomonas spp.* .

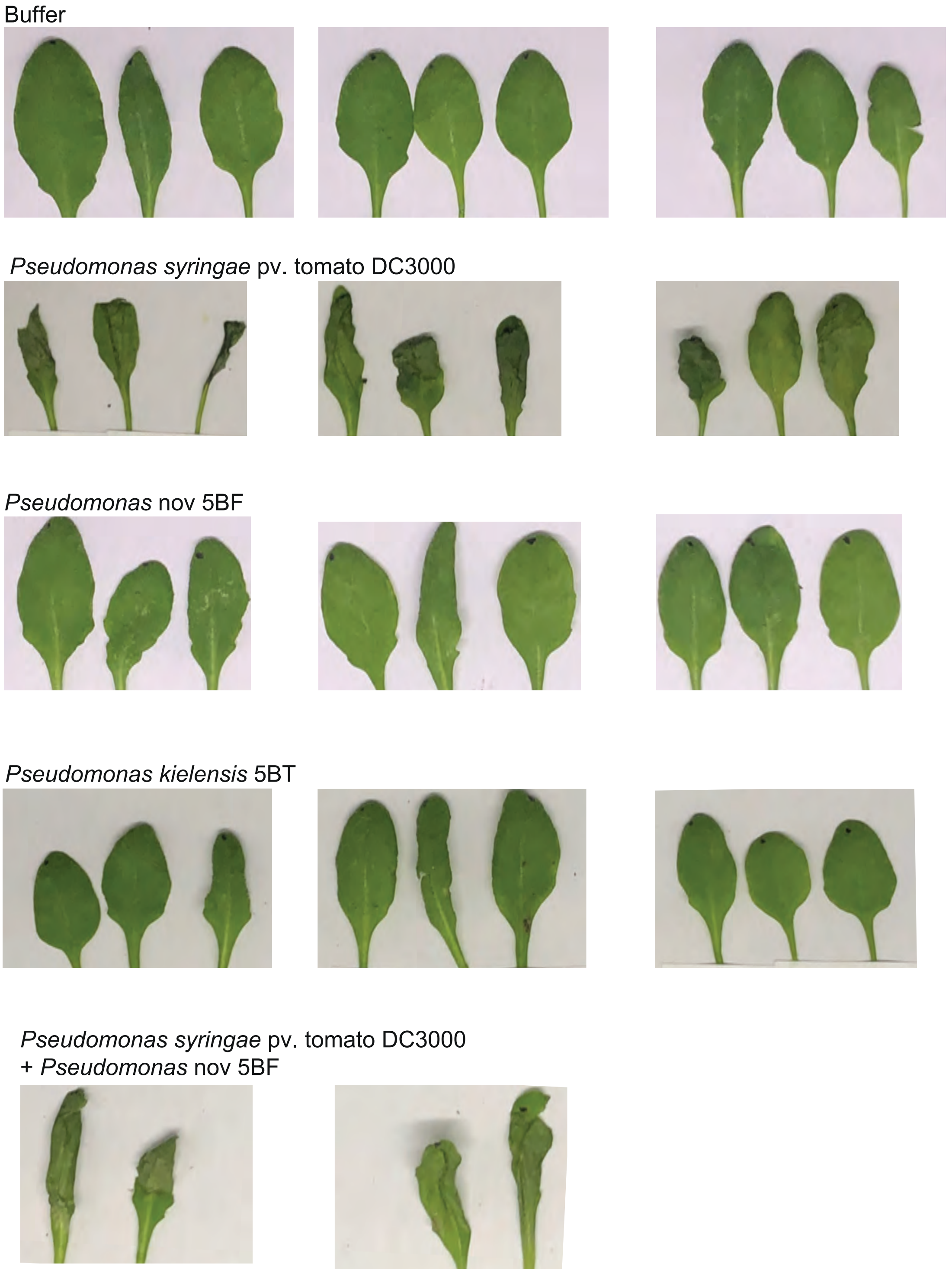


**Figure S30. High OD syringe infiltration of *A. thaliana* leaves with *Pseudomonas spp.***
Experiment 2 of high OD *Pseudomonas spp.* inoculations. Photographs of *A. thaliana* leaves 48hrs post inoculation with various *Pseudomonas spp.* .

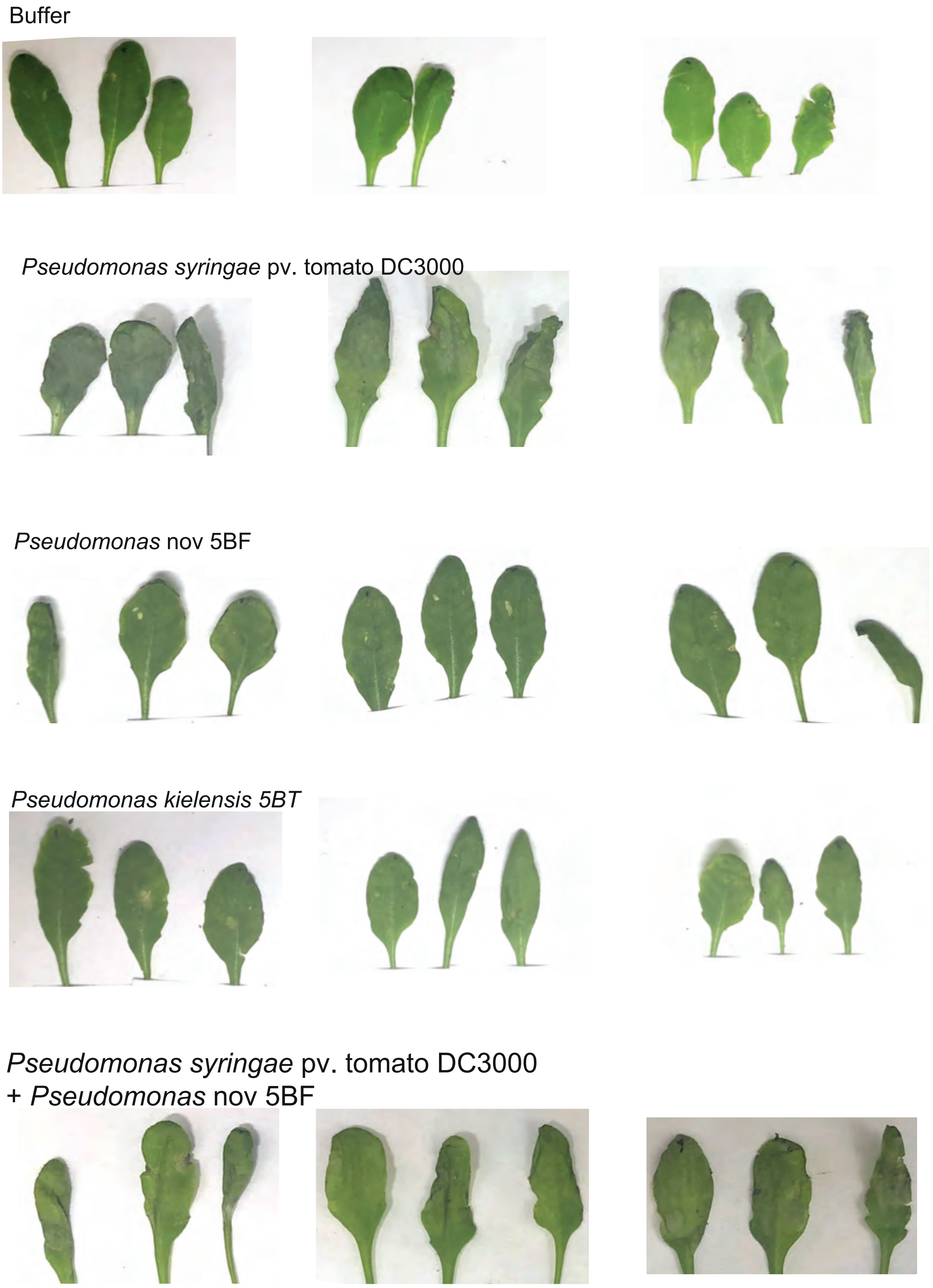


**Figure S31. High OD syringe infiltration of *A. thaliana* leaves with *Pseudomonas spp.***
Experiment 3 of high OD *Pseudomonas spp.* inoculations. Photographs of A. thaliana leaves 48hrs post inoculation with various *Pseudomonas spp*. .

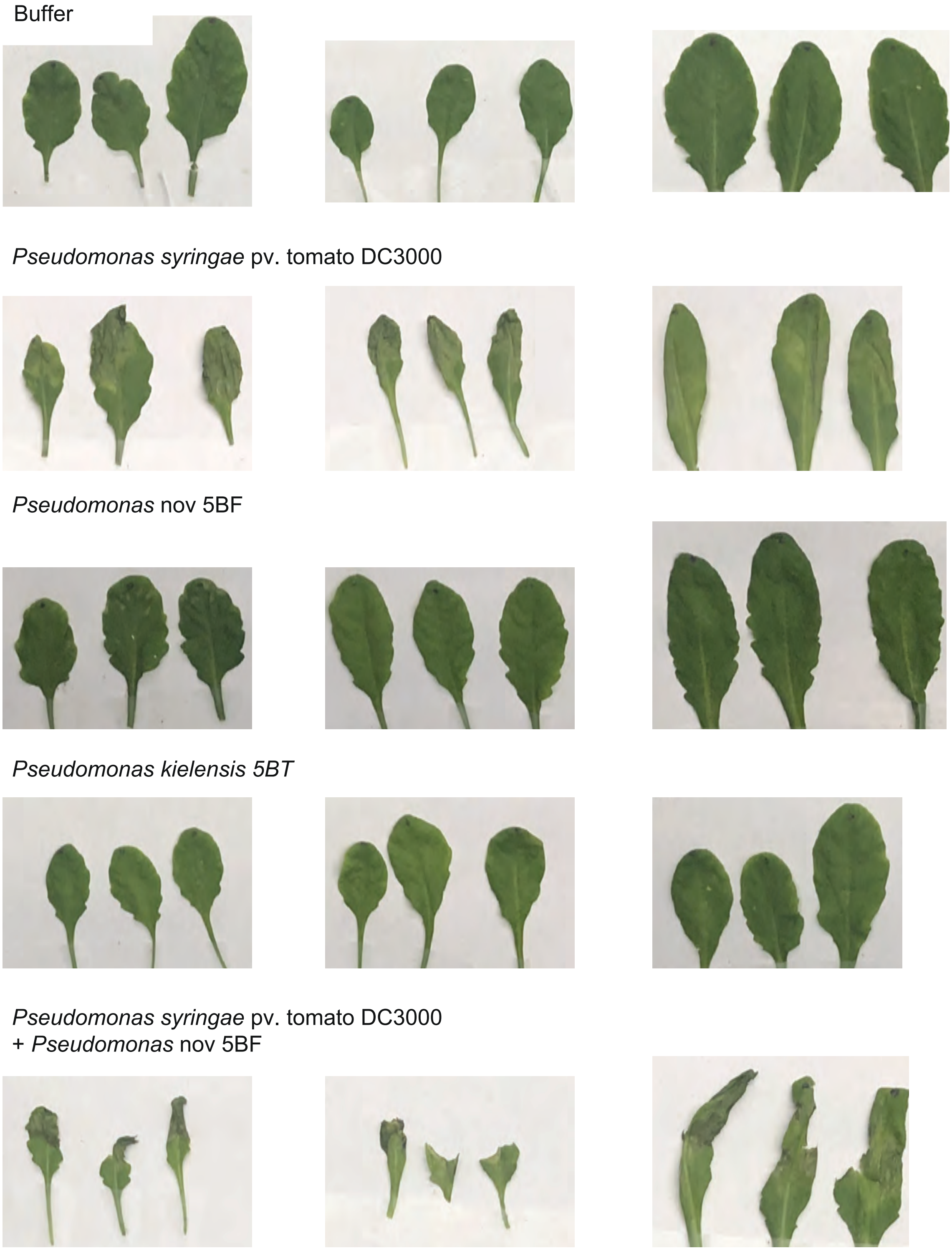


**Figure S32. High OD syringe infiltration of *A. thaliana* leaves with *Pseudomonas spp.*** Experiment 4 of high OD *Pseudomonas spp.* inoculations. Photographs of *A. thaliana* leaves 48hrs post inoculation with various *Pseudomonas spp.*

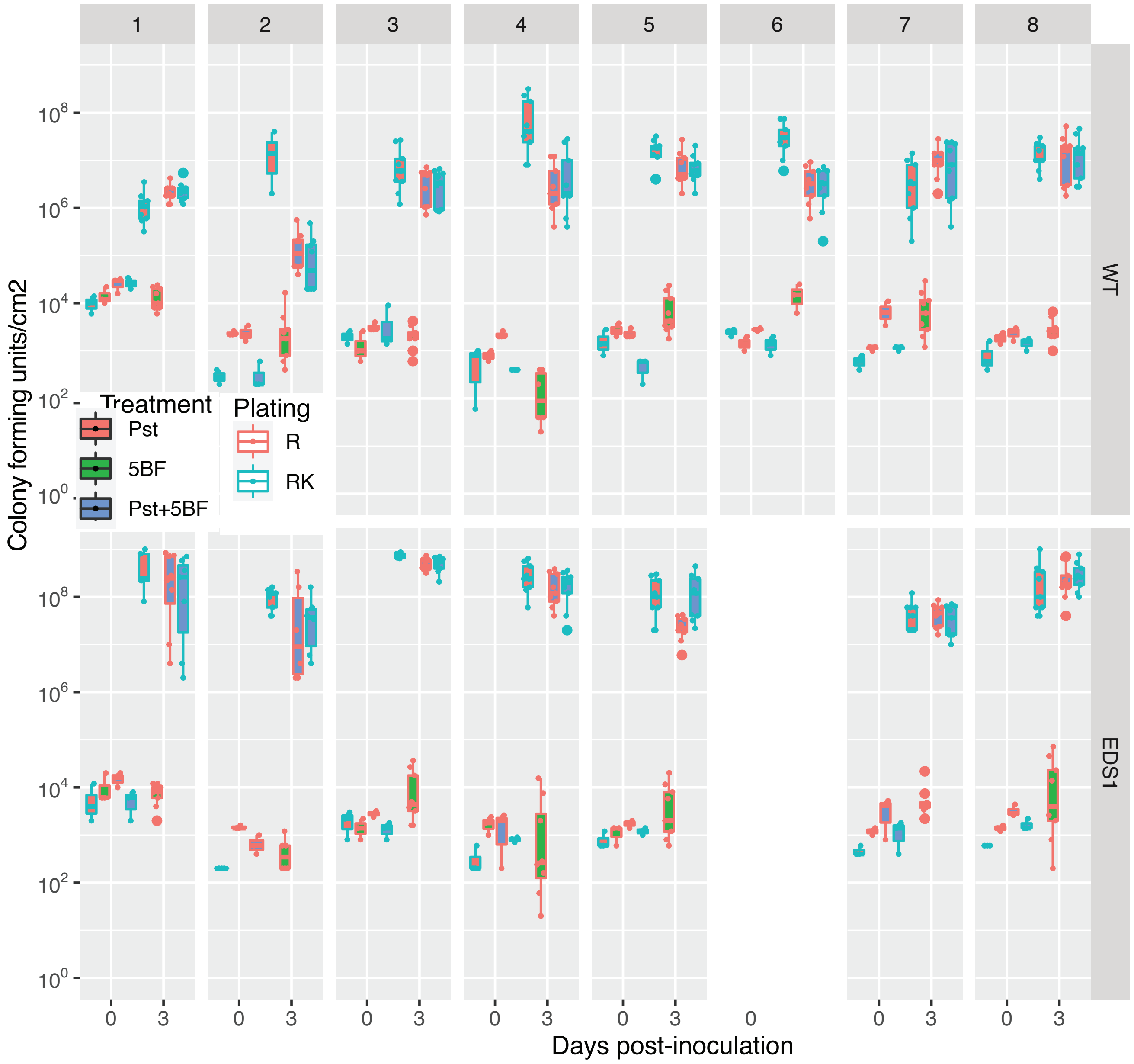


**Figure S33. Low OD syringe infiltration of *A. thaliana* leaves with *Pseudomonas spp*.**
Growth curves as in figure 7 separated by experiment for infiltrations of *Pseudomonas* 5BF and *Pseudomonas syringae* DC3000 together and individually on either A. thaliana WT or *eds1*.

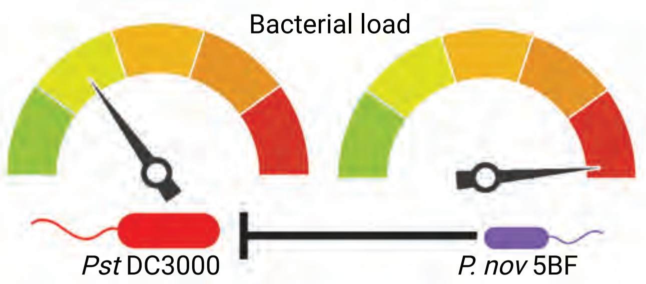


In conditions where *P. nov.* 5BF is able to grow it can suppress the growth of *Pst* DC3000

**Figure S34. Model of bacterial plant interactions**
